# Cortical sensory aging is layer-specific

**DOI:** 10.1101/2023.12.01.567841

**Authors:** Peng Liu, Juliane Doehler, Julia U. Henschke, Alicia Northall, Angela Serian, Laura C. Loaiza-Carvajal, Eike Budinger, Dietrich S. Schwarzkopf, Oliver Speck, Janelle M.P. Pakan, Esther Kuehn

## Abstract

The segregation of processes into cortical layers is a convergent feature in animal evolution. However, how changes in the cortical layer architecture interact with sensory system function and dysfunction remains unclear. We conducted functional and structural layer-specific in-vivo 7T-MRI of the primary somatosensory cortex in two cohorts of healthy younger and older adults. Input layer IV is enlarged and more myelinated in older adults, and associated with extended sensory input signals. Age-related cortical thinning is driven by deep layers and accompanied by increased myelination, but there is no clear evidence for reduced inhibition. Calcium imaging and histology in younger and older mice reveal increased sensory-evoked neuronal activity accompanied by increased parvalbumin expression as a potential inhibitory balance, with dynamic changes in layer-specific myelination across age groups. Using multimodal imaging, we demonstrate that middle and deep layers show specific sensitivity to aging across species.

## Introduction

Sensory processing is organized in a layered architecture with segregated input, output, and modulatory circuits. The layered architecture of sensory systems is a convergent feature in animal evolution^1^. A model of functional and dysfunctional sensory systems requires a detailed understanding of alterations in the layer-specific architecture and their associated phenotypes. Critically, this knowledge is so far lacking not only for sensory systems but for cortical dysfunction in general.

Sensory dysfunction has been associated with different cortical phenotypes, including functional overactivation^2^, increases in receptive field sizes^2,3^, decreases in lateral inhibition^2^, and structural alterations such as cortical thinning^4^. However, it is unclear how changes in the cortical layer architecture may contribute to the specific functional and structural alterations that characterize sensory cortices with reduced functionality. Cortical aging serves as a suitable model system to investigate this question as structural and functional plasticity is observed at different levels of the processing hierarchy and affects behavior^2^.

We tested four major hypotheses of how structural and functional alterations in the cortical layer architecture characterize cortical dysfunction. The ‘preserved layer hypothesis’ assumes that the structural layer architecture cannot be distinguished between younger and older adults with present methodology, motivated by the preserved layer architecture of primary motor cortex (MI) in older adults^5^. Conversely, the ‘altered input channel hypothesis’ assumes that age-related functional plasticity is characterized by a more (or less) pronounced input layer IV in older compared to younger adults. Previous studies have shown structural sensitivity of layer IV to sensory input statistics^6^, which are higher in older adults due to increased age, but also weakened due to reduced peripheral nerve receptor density^7^. Our third, ‘altered sensory modulation hypothesis’ assumes that older compared to younger adults are characterized by a degradation of deep layers V and VI, which mediate changes in the excitatory/inhibitory modulation of sensory inputs. In primates, deep layers of sensory cortex mediate inhibition^8^, whereas in older adults, inhibition is reduced^2^. Finally, the ‘degraded border hypothesis’ assumes that low-myelin borders that exist inferior to the thumb representation^9^, and occasionally between single finger representations^10^, in input layer IV degrade with cortical aging and are associated with less precise functional representations^3^.

To test these hypotheses, we employed a unique approach and acquired layer-specific functional and structural magnetic resonance imaging (fMRI) data using 7T-MRI of the primary somatosensory cortex (SI) of healthy younger and older adults together with behavioral assessments. To better understand the mechanistic underpinnings, we analyzed younger, older and senescent mice using *in-vivo* two-photon calcium imaging and histological analyses (see **Figure 1**). In humans, analyses were restricted to area 3b - the homologue of mouse SI^11^. This study inspires the precise investigation of cortical layer dynamics to reveal fundamental mechanisms that underlie cortical dysfunction.

**Figure 1.**
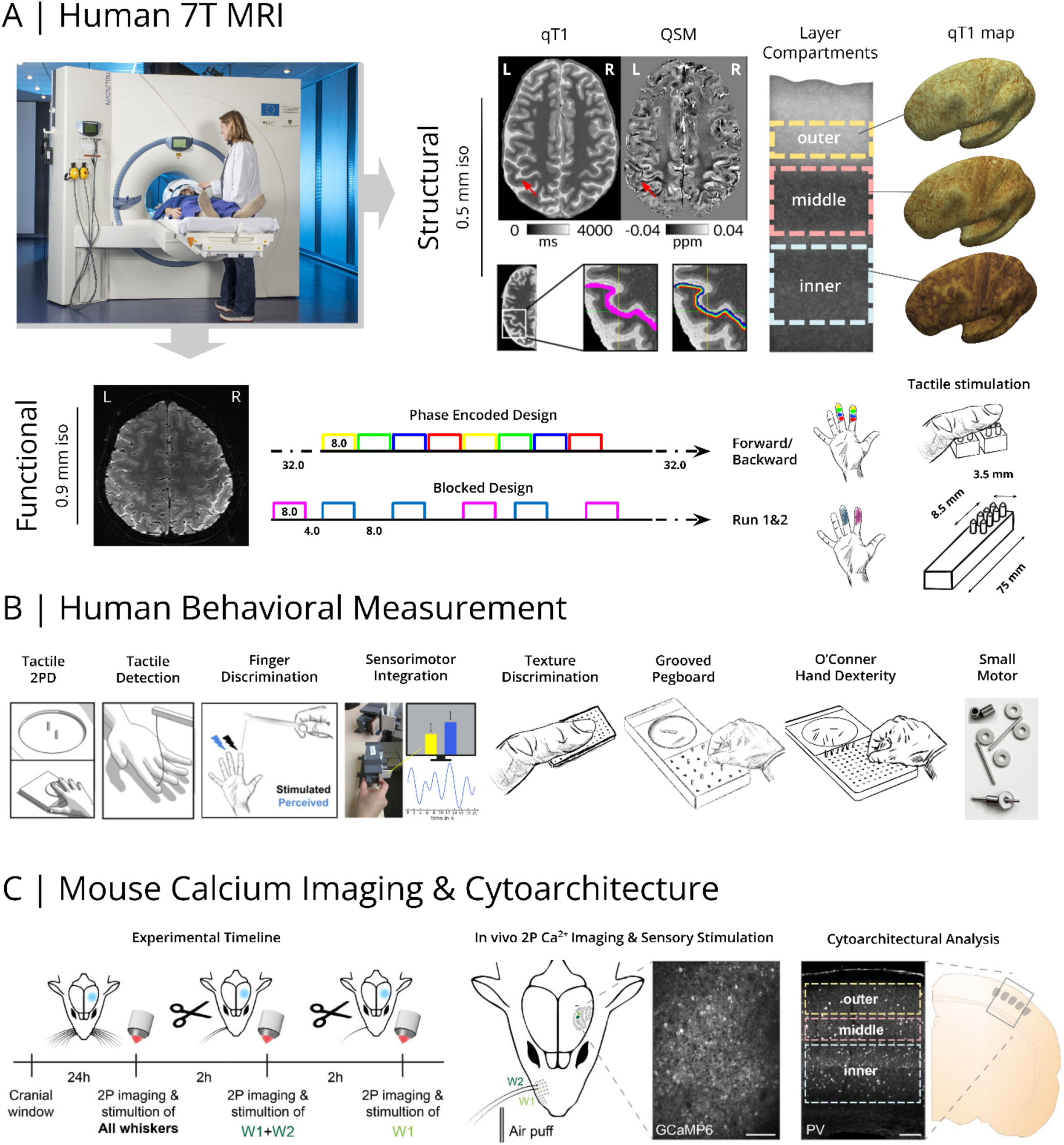
Overview Experimental Design. **(A)** 7 Tesla Magnetic Resonance Imaging (7T-MRI) was used to investigate the layer-specific structural architecture of primary somatosensory cortex (SI) in younger and older adults (red arrows indicate region of interest). MP2RAGE data were used to define SI layer compartments in-vivo. Quantitative T1 (qT1) and quantitative susceptibility mapping (QSM) values were mapped onto the individuals’ inflated, layer-specific cortical surfaces and used for further analyses. Myelin staining was remodeled according to Dinse et al.^12^. 7T functional MRI (7T-fMRI) was used to investigate age- and layer-specific functional changes in SI. Participants underwent several passive tactile stimulation paradigms (phase-encoded design and blocked design) in the scanner where fingers of the right hand were stimulated. L=left, R=right side of the human brain. **(B)** Younger and older adults underwent a behavioral test battery including tactile 2-point discrimination (2PD), tactile detection, tactile finger discrimination, sensorimotor integration, and tactile texture discrimination tasks as well as the Grooved Pegboard Test, the O’Connor Hand Dexterity Test and the Small Motor Test. (**C**) Barrel cortex two-photon calcium imaging (2PCI) of excitatory neurons expressing a genetically encoded calcium indicator (GCaMP6f) during airpuff whisker stimulation was used to investigate younger and older adult mice: i) during baseline conditions with all whiskers, ii) after all whiskers were cut except two on the right-side (W1+W2; double stimulation condition), iii) after another whisker was cut leaving only one (W1; single stimulation condition). Post-mortem histological analysis examined underlying cytoarchitectural differences across cortical layers and aging. Scale bars 100 µm.

## Results

### Age-related cortical thickness changes are layer-specific

Our first hypothesis expects the structural layer architecture to be similar between younger and older adults (‘preserved layer hypothesis’). According to this view, if cortical thinning occurs with aging, we expect homogenous thinning of all layers. To target this fundamental aspect of cortex organization, we analyzed structural and functional 7T-MRI data of younger (n=20) and older (n=20) adults’ SI (cohort 1) using the SI hand area as a model system. We localized the SI hand area using tactile population receptive field (pRF) modeling^3^. The SI layer definition followed a data-driven approach, assigning biological layers II and III to the outer compartment, input layer IV to the middle compartment, and layers V and VI to the inner compartment^10^. In agreement with previous findings^13^, the SI hand area is on average 2.00±0.10 mm thick (Mean±SD), but thinner in older compared to younger adults (older-younger=-0.12 mm; see **Table 1**)^4^. Critically, overall cortical thinning is driven by reduced thickness of the inner compartment, whereas the middle compartment is thicker in older compared to younger adults (see **Table 1** and Figure 2B,**D**, see **Table 1-supplemental table 1** for a different localization approach, see **Table 1-supplemental table 2** for a different layer compartmentalization scheme, see **Table 1-supplemental table 3** for cortical thickness analysis without outlier removal, see **Table 1-supplemental** figure 1 for the exact definition of cortical layer compartments, see Fig. 2**-supplemental** figure 1 for the distribution of layer-specific cortical thickness values).

**Figure 2.**
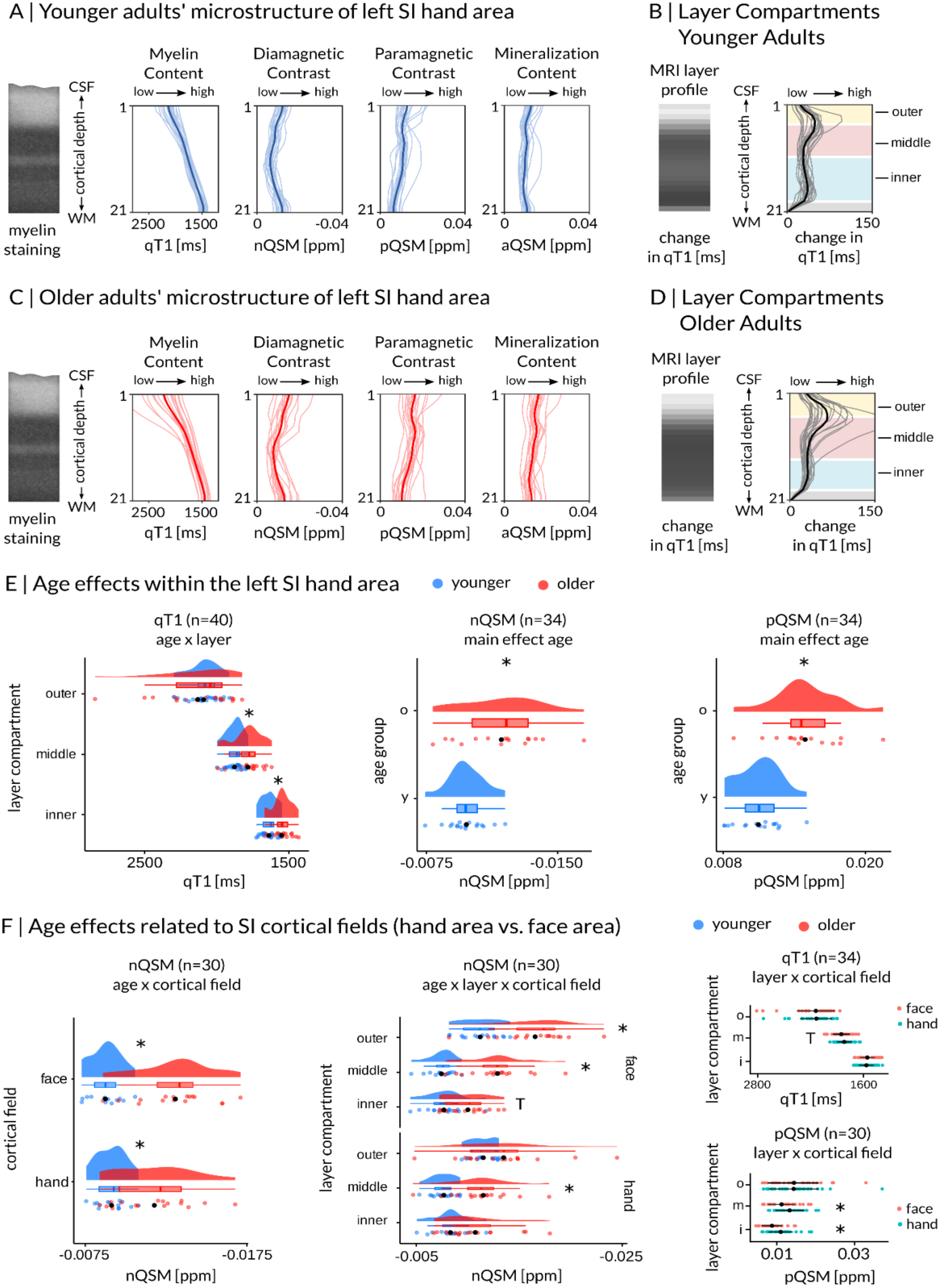
Age-Related Differences in the Structural Layer Architecture in Human SI. **(A)** Younger adults’ microstructure described by qT1-based myelin (n=20), nQSM-based diamagnetic contrast (indicative of calcium content/metabolism; n=18), pQSM-based paramagnetic contrast (indicative of iron content; n=18), aQSM-based mineralization (n=18) sampled between the CSF/GM border and the GM/WM border (group mean plotted in bold). Low qT1 and nQSM indicate high myelin content and high diamagnetic contrast, respectively. Myelin staining was remodeled according to Dinse et al.^12^. **(B)** Layer compartments of younger adults (cream, outer layer; light pink, middle layer; light blue, inner layer) calculated based on the mean local rate of change in intracortical qT1^10^. The middle layer compartment (light pink) is assumed to reflect the heavy-myelinated Baillarger band running through anatomical layer IV. MRI layer profiles visualise the local rate of change in intracortical qT1 as shades of grey. **(C)** Older adults’ microstructure described by qT1-based myelin (n=20), nQSM-based diamagnetic contrast (n=16), pQSM-based paramagnetic contrast (n=16), aQSM-based mineralization (n=16) sampled between the CSF/GM border and the GM/WM border (group mean plotted in bold). QSM values are given in parts per million (ppm). **(D)** Layer compartments of older adults calculated based on the mean local rate of change in intracortical qT1 (black line). The comparison between (B) and (D) visualizes age-related differences in the layer architecture of SI. **(E)** Age-related differences in qT1, nQSM and pQSM in the SI hand area (permutation mixed-effects ANOVAs), individual data shown as colored dots: younger adults [y] in blue, older adults [o] in red; group means shown as black dots. Box plots are drawn within the interquartile range (box), medians are shown as vertical lines, whiskers connect the minimum and the maximum with the lower and the upper quartiles. Significant differences at Bonferroni-corrected significance level of p < 0.0045 are marked by *. **(F)** Age-related differences in SI cortical fields. Shown are permutation mixed-effects ANOVAs with factors age (younger adults colored in blue, older adults colored in red), layer (outer [o], middle [m], inner [i]) and cortical field (hand area, face area) on residual qT1, nQSM and pQSM after regressing out the effect of map size. Significant differences at Bonferroni-corrected significance level of p < 0.002 are marked by a *. Trends above Bonferroni-corrected threshold are marked by a T. For similar data of n=1 participant with congenital loss of the right arm see **Table 1-supplemental** figure 3 for microstructural profiles and layer compartmentalization scheme in reference to younger and older adults, **Table 1-supplemental** figure 4 for microstructural profiles with map size adjusted ROIs, and **Table 1-supplemental table 5** for quantitative values of the microstructural composition of SI layers.

**Table 1.** Layer-specific cortical thickness values of the SI hand area. Shown are total and layer-specific (outer, middle, inner) mean cortical thickness values (Mean) and standard deviations (SD) in millimeters for the SI hand area. Independent-samples random permutation Welch t-tests were calculated to investigate group differences (t=test statistic, df=degrees of freedom, pperm=Monte-Carlo permutation p-value, CIperm=95% Monte-Carlo permutation confidence interval, number of permutations=100000, minimum value of pperm=1/number of permutations). Significant differences with Bonferroni-corrected p < 0.0125 (correcting for 4 tests) are marked by *. See **Table 1-supplemental table 4** for Bayesian independent-sample t-tests computed on these differences. Data of n=19 older adults are presented here after outlier removal. The full data set can be found in **Table 1-supplemental table 3**.

| hand area | All<br>n=39 | Younger<br>n=20 | Older<br>n=19 | Group Differences |  |  |  |
| --- | --- | --- | --- | --- | --- | --- | --- |
| | Mean ± SD | Mean ± SD | Mean ± SD | *t* | *df* | *p*<sub>perm</sub> | *CI*<sub>perm</sub> |
| total | 2.00 ± 0.10 | 2.06 ± 0.07 | 1.94 ± 0.08 | 5.0 | 35.8 | < 10<sup>-5</sup> * | 0.062, 0.183 |
| outer | 0.40 ± 0.02 | 0.41 ± 0.02 | 0.40 ± 0.02 | 1.3 | 34.7 | 0.196 | -0.004, 0.023 |
| middle | 0.70 ± 0.15 | 0.56 ± 0.02 | 0.85 ± 0.03 | -34.6 | 28.3 | < 10<sup>-5</sup> * | -0.386, -0.198 |
| inner | 0.90 ± 0.21 | 1.10 ± 0.06 | 0.69 ± 0.04 | 26.5 | 31.3 | < 10<sup>-5</sup> * | 0.275, 0.539 |

Bayesian independent-sample t-tests confirm this result, showing extreme evidence for reduced total cortical thickness (BF_10_=1383.79), reduced inner layer thickness (BF_10_=2.05×10^26^), and increased middle layer thickness in older adults compared to younger adults (BF_10_=8.93×10^21^), and anecdotal evidence for the null hypothesis of no group difference for the outer layer (BF_10_=0.635, see **Table 1-supplemental table 4**). This result of layer-specific cortical thickness differences between younger and older adults rejects the ‘preserved layer hypothesis’.

We also examined a healthy adult (male, age=52 years) with congenital arm loss on the right side to further investigate variations in cortical thickness in the area 3b hand area (see **Table 1-supplemental** figure 2 for functional localizer), detailed results are included in **Table 1-supplemental** figure 3 and **Table 1-supplemental** figure 4.

### Low-myelin borders have a similar architecture in younger adults, older adults, and with congenital arm loss

Next, we tested the ‘degraded border hypothesis’. This hypothesis assumes that the low-myelin borders that exist inferior to the thumb representation^9^, and occasionally between single finger representations^10^, in input layer IV of SI degenerate with age. We first confirmed that the overall structural topographic architecture of SI follows the expected pattern, i.e., individual finger representations do not significantly differ in their microstructural composition (see Fig. 2**-supplemental table 1,** Fig. 2**-supplemental table 2**), whereas the hand and the face areas show microstructural differences, indicating distinct cortical fields^10^. In particular, we show more pronounced diamagnetic contrast in the face area in older adults (see Figure 2F, Fig. 2**-supplemental table 3** and Fig. 2**-supplemental table 4**).

To test our hypothesis, we developed an automatic border detection algorithm (see Fig. 2**-supplemental** figure 2), which did not detect age-related differences with respect to the existence or composition of layer IV low-myelin borders between younger and older adults (see Fig. 2**-supplemental** figure 2). Statistical trends that indicate differences in iron and calcium content in the hand-face border between age groups (see Fig. 2**-supplemental** figure 2) are not specific to the border area but are generally observed in SI.

In an exploratory analysis, the same automatic border detection algorithm was applied to the data acquired from n=1 congenital one-hander, revealing that low-myelin borders exist in both hemispheres of this individual, i.e., contralateral and ipsilateral to the missing arm (see Fig. 2**-supplemental** figure 3). Low-myelin borders therefore have a similar architecture in younger adults, older adults, and in n=1 individual with congenital arm loss. This rejects the ‘degraded border hypothesis’, confirming a previous observation in human MI^5^.

### More pronounced sensory input signals in layer IV in older adults

The above-reported results reject both the ‘preserved layer hypothesis’ and the ‘degraded border hypothesis’. However, the increased thickness of input layer IV in older adults is in line with the ‘altered input channel hypothesis’. To follow this up, we investigated, in an independent cohort (cohort 2, n=11 younger adults, n=10 older adults), if there is evidence for extended sensory input signals in layer IV in older compared to younger adults. We extracted both resting state and tactile stimulation-induced %signal change of index and middle finger representations (see Figure 3, see Fig. 3**-supplemental** figure 1 and Fig. 3**-supplemental** figure 2 for data of all participants) in contralateral SI for each individual at each modeled cortical depth. In both younger and older adults, there is an antagonistic center-surround relationship between signals and cortical depth: signals peaked in the input layer IV but were minimal in neighboring depths. Given we only observed this relationship during sensory stimulation but not during the resting state (see Figure 3E versus **3F**), and the peak occurred at the expected layer compartment encompassing layer IV, the signal peak is a marker to measure input signals to layer IV in SI.

**Figure 3.**
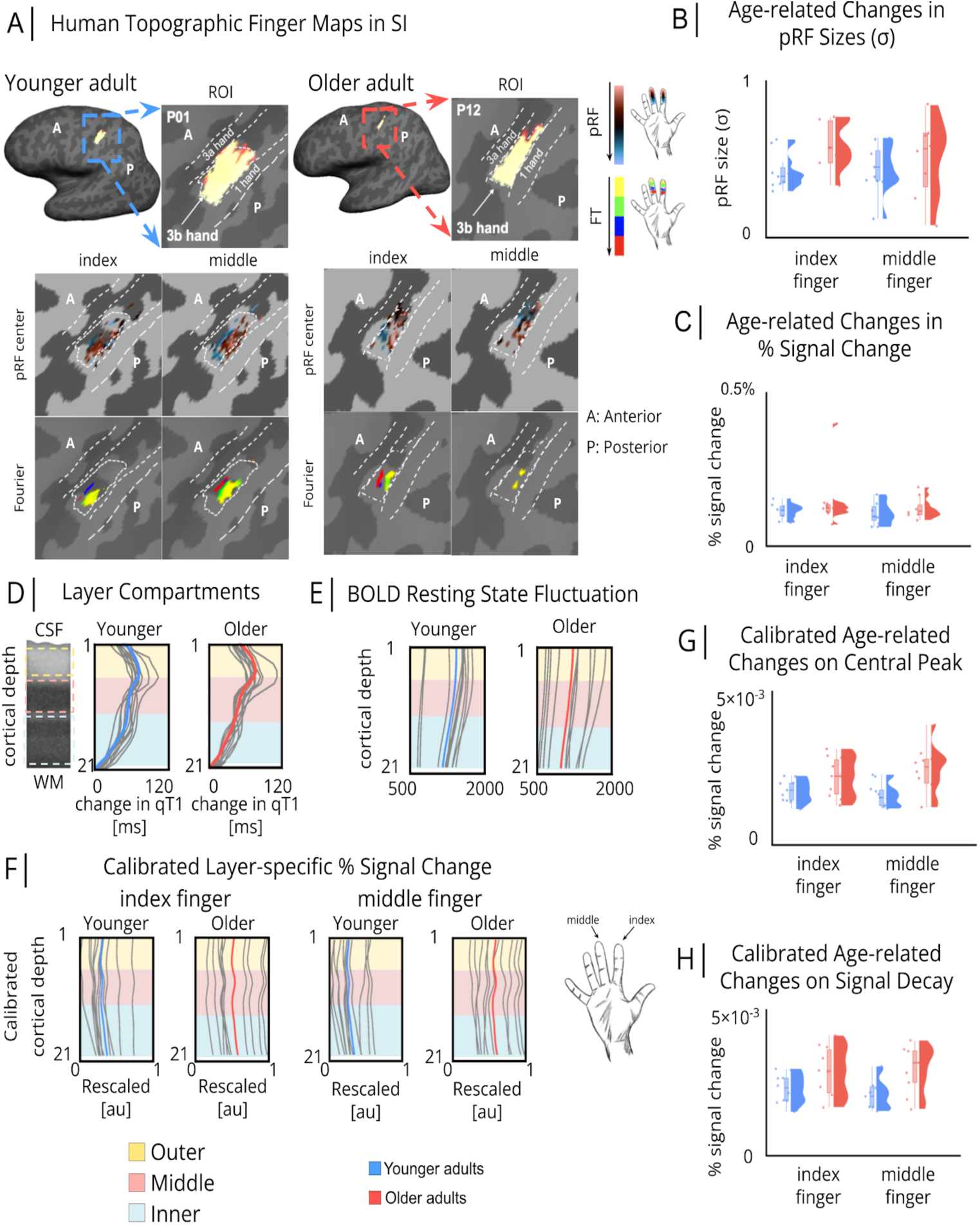
Layer-specific Functional Architecture of Human SI. **(A)** Individual topographic maps of the index finger and the middle finger, shown for one younger adult and one older adult (randomly chosen, younger: P01 and older: P12, see Fig. 3**-supplemental** figure 1 for all individual data). Shown are topographic maps extracted using population receptive field (pRF) modeling (first rows in each panel) and Fourier-based mapping (second rows in each panel). ROI=Region-of-interest (hand area of contralateral area 3b). **(B)** pRF size (σ) estimates of index and middle finger representations (with arbitrary units (au)) individual data shown as colored dots: younger adults [y] in blue, older adults [o] in red. Box plots are drawn within the interquartile range (box), medians are shown as vertical lines, whiskers connect the minimum and the maximum with the lower and the upper quartiles. **(C)** % signal change of index and middle finger representations, individual data shown as colored dots: younger adults [y] in blue, older adults [o] in red. Box plots are drawn within the interquartile range (box), medians are shown as vertical lines, whiskers connect the minimum and the maximum with the lower and the upper quartiles. **(D)** Layer compartments (cream: outer layer, light pink: middle layer, light blue: inner layer) for younger (n=11, blue) and older (n=10, red) adults defined based on rate of change in qT1^10^. (**E**) Resting state signal fluctuation extracted at different cortical depths for younger (blue) and older (red) adults. **(F)** %signal change of index and middle finger representations using calibrated-BOLD (Blood Oxygen Level Dependent) extracted at different cortical depths for younger (blue) and older (red) adults (see Fig. 3**-supplemental** figure 3 for %signal change without calibration). Values are scaled between 0 and 1. **(G)** Central peak of index and middle finger representations, individual data shown as colored dots: younger adults [y] in blue, older adults [o] in red. Box plots are drawn within the interquartile range (box), medians are shown as vertical lines, whiskers connect the minimum and the maximum with the lower and the upper quartiles. **(H)** Signal decay of index and middle finger representations, individual data shown as colored dots: younger adults [y] in blue, older adults [o] in red. Box plots are drawn within the interquartile range (box), medians are shown as vertical lines, whiskers connect the minimum and the maximum with the lower and the upper quartiles.

Older adults presented with a more pronounced sensory input peak compared to younger adults (see Figure 3F-H, for detailed statistics see Fig. 3**-supplemental table 1**, note that analyses were performed on calibrated BOLD, reducing the likelihood that effects are driven by age-related differences in neurovascular coupling). This provides further evidence towards the ‘altered input channel hypothesis’, according to which the cortical input circuit changes with aging, though reduced inhibition could also explain this effect (‘altered modulation hypothesis’).

### Altered functional response profile in older adults’ SI

So far, we have rejected both the ‘preserved layer hypothesis’ and the ‘degraded border hypothesis’, and have provided evidence for the ‘altered input channel hypothesis’. With respect to the third, ‘altered modulation hypothesis’, we hypothesized that deep layer degeneration (evidenced above by age-related deep layer thinning) is related to coarser functional selectivity and/or reduced coactivated inhibition in older adults^2^. To test for functional selectivity, we computed the sharpness of index and middle finger pRFs (sigma parameter (σ)). To test for coactivated inhibition, we compared mean %signal change and mean pRF sizes between a ‘double stimulation condition’ and a ‘single stimulation condition’^14^. Note that the double-versus-single-stimulation comparison implies comparing single finger versus double finger stimulation in humans, and single versus double whisker stimulation in mice (for the latter see section ‘Sensory-evoked neuronal activity and altered layer architecture in mice’).

Results show strong evidence for coarser spatial selectivity of the index finger representation in older compared to younger adults, whereas the middle finger shows anecdotal evidence favouring the null hypothesis (for detailed statistics see Fig. 3**-supplemental table 2**). Against our expectation, statistical evidence for age differences in markers of coactivated inhibition is inconclusive (for detailed statistics see Fig. 3**-supplemental table 3**). This suggests that middle layer expansion and deep layer thinning co-occur with coarser spatial selectivity of the index finger representation, but we cannot confirm that this co-occurs with decreased inhibition in older adults.

### Altered microstructural layer composition in older adults

Next, we investigated how the microstructural composition of SI layers changes with age. We extracted qT1 values as a proxy for cortical myelin^12^ and quantitative susceptibility maps (QSM) as proxies for cortical iron (positive values, pQSM) and calcium/metabolism (negative values, nQSM)^15^, as well as overall mineralization (absolute values, aQSM)^16^, from different depths of the human hand area. We confirm previously reported across-layer age effects^5,16^: (1) more negative nQSM values (older: −0.0118±0.0006 parts per million (ppm); younger: −0.0098±0.0003 ppm), (2) higher pQSM values (older: 0.0149±0.007 ppm; younger: 0.0110±0.0004 ppm), and (3) higher aQSM values (older: 0.0138±0.0006 ppm; younger: 0.0103±0.0003 ppm) in older compared to younger adults (see Fig. 2**-supplemental table 5**, Fig. 2**-supplemental table 6**).

Interestingly, age effects in qT1 values are layer-specific, with lower qT1 values (reflecting higher myelin) in the inner and middle compartments in older compared to younger adults (older: inner=1547.6±14.8 ms (Mean±SE); middle=1783.2±21.9 ms; younger: inner=1636.3±12.3 ms; middle=1874.4±12.9 ms; see Figure 2E, see Fig. 2**-supplemental table 5** and Fig. 2**-supplemental table 6** for exact results, see Fig. 2**-supplemental table 7** and Fig. 2**-supplemental table 8** for control analyses using a different compartmentalization scheme). Controlling for effects of cortical atrophy, there is no significant difference in hand mask (full hand map) size (qT1 (n=40): *t*(33.7)=-0.48, *p*=0.632; nQSM (n=34): *t*(31.8)=0.67, *p*=0.510; pQSM (n=34): *t*(30.1)=-0.39, *p*=0.696; aQSM (n=34): *t*(29.4)=0.02, *p*=0.981) between younger (qT1: 1249±90; nQSM: 512±53; pQSM: 717±62; aQSM: 1271±98 vertices, Mean±SE) and older adults (qT1: 1325±130; nQSM: 462±53; pQSM: 756±75; aQSM: 1268±124 vertices).

These results are in line with the ‘altered input channel hypothesis’: The input layer IV does not only appear thicker and shows more pronounced sensory input signals in older compared to younger adults, but it is also characterized by higher myelin content. These results are also in line with the ‘altered modulation hypothesis’: The deeper layers of SI are thinner in older adults and the functional spatial selectivity of the index finger representation is coarser, but this is not accompanied by evidence towards reduced functional inhibition. The reason may be the higher myelin content in deeper layers in older adults. To clarify the role of age-related alterations in layer-specific cellular composition more precisely, additional analyses were conducted in younger and older mice.

### Sensory-evoked neuronal activity and altered layer architecture in mice

While BOLD responses are difficult to relate to precise neural excitatory or inhibitory drive, and MRI data are difficult to relate to precise histological changes, the use of animal models can help us to clarify mechanistic aspects related to the cytoarchitectural changes observed in specific layers and associated functional changes. We examined the aging whisker barrel cortex in mice as an analogous sensory system that shares multiple topological features and aspects of cortical representation with the human hand and fingers^11^. We investigated the activity of individual neurons in the equivalent of younger adults (mice 19.6 ± 2.5 weeks, n=8, i.e., between 2-6 months) and older adults (mice 76.7 ± 2.2 weeks, n=8, i.e., between 12-20 months) and combined this with post mortem histological analysis across cortical layers and age.

To first examine sensory-driven neuronal activity in younger compared to older adult mice, a cranial window was implanted above the barrel cortex in transgenic mice expressing a genetically encoded calcium indicator (CGaMPf)^17^ driven by a Thy1 promoter^18^. We examined sensory-evoked responses of excitatory neurons across the outer (layer II/III) and inner (layer V) cortical layers following spontaneous activity and direct airpuff stimulation to all whiskers as well as subsets of whiskers (see Figure 1C, i.e., similar stimulation paradigm to humans: ‘single stimulation condition’, one whisker [W1], ‘double stimulation condition’, two neighboring whiskers coactivated [W1+W2]; see Fig. 4**-supplemental** figure 1). Changes in fluorescence (ΔF/F) of the calcium indicator were extracted and used as a proxy readout of neuronal activity. We observed a more pronounced increased amplitude of excitatory sensory-evoked responses in older adult mice during direct whisker stimulation of all whiskers as well as during voluntary whisking compared to spontaneous neuronal responses during rest (see Figure 4A-B, see also Fig. 4**-supplemental** figure 1; younger adult [n=1519 neurons] vs older adult [n=1958 neurons]: air-puff whisker stimulation: t(3475)=14.6, *p* < 0.001, Cohen’s *d*=0.498; natural voluntary whisking: t(3475)=16.9, *p* < 0.001, Cohen’s *d*=0.577; spontaneous neuronal responses: t(3475)=10.7, *p* < 0.001, Cohen’s *d*=0.367; independent-sample t-test, data from 8 mice per age group (see also Fig. 4**-supplemental** figure 1B)). This indicates more pronounced sensory input signals in older adult mice, analogous to that observed in humans (see Figure 3C,**F**).

**Figure 4.**
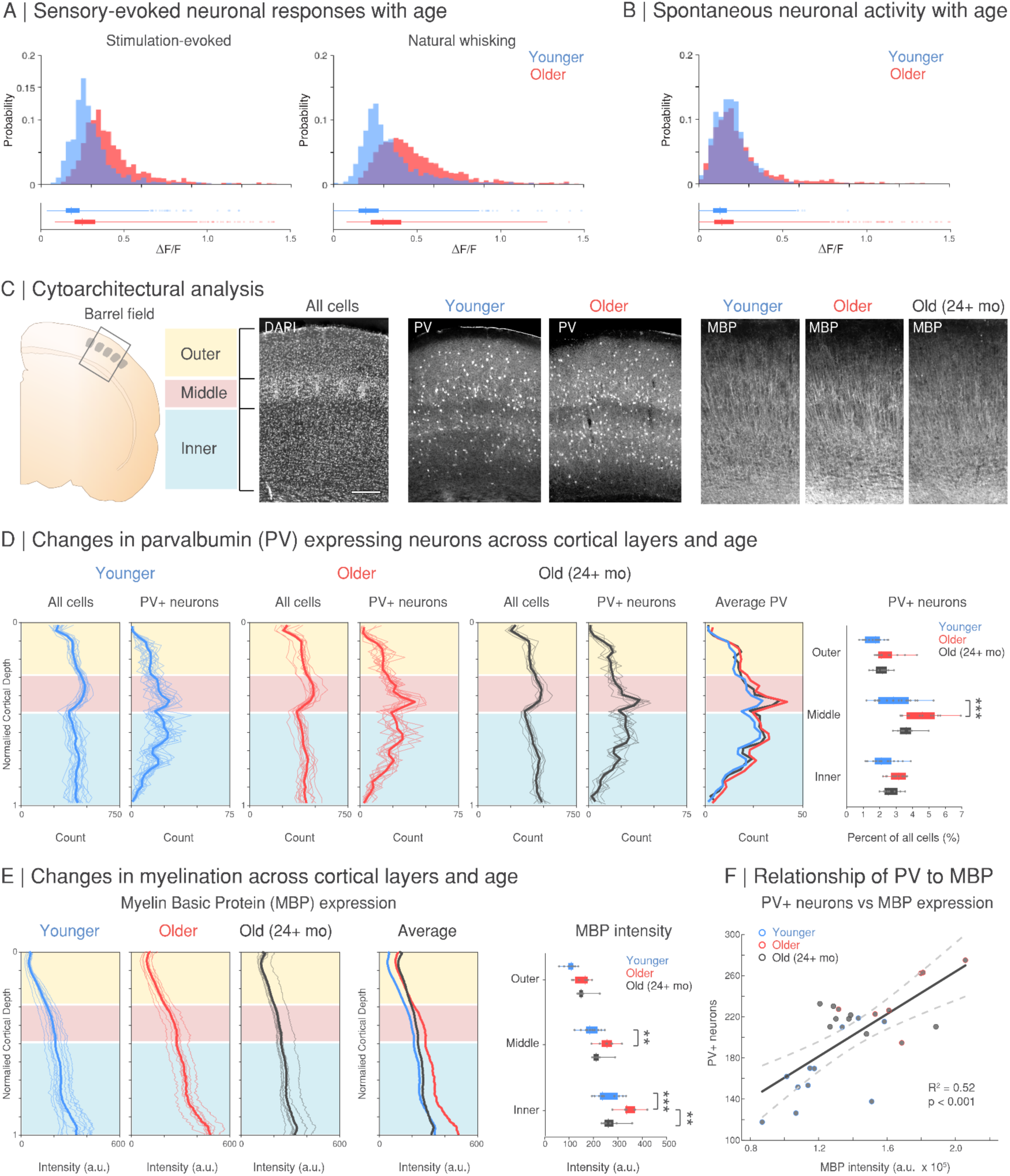
Age-Related Functional and Structural Changes in Mouse Barrel Cortex. Average change in fluorescence (ΔF/F) for all neurons across younger (n=1519 neurons) and older (n=1958 neurons) adult mice (from 8 mice for each age group) during **(A)** sensory-evoked neuronal activity following air-puff stimulation (left) and during periods of natural whisking (right) and **(B)** spontaneous neuronal activity during periods with no whisker pad movement. **(C)** Schematic of the mouse barrel field where images of 4′, 6-Diamidin-2-phenylindol (DAPI) staining of all cells and parvalbumin (PV) for younger adult mice [n=17, 2-6 months], older adult mice [n=13, 12-20 months], and mice in old age [n=8, +24 months]) and myelin basic protein (MBP) immunohistochemistry from the barrel cortex were collected for analysis in a subset of these mice (younger adult mice [n=11, 2-6 months], older adult mice [n=7, 12-20 months], and mice in old age [n=8, +24 months]) and across cortical layers (inner, layer 2/3; middle, layer IV; outer, layer V-VI). Scale bar 100µm. Quantification of the binned sum of PV-expressing cells **(D)** and intensity of MBP staining **(E)** is shown across the cortical depth, normalized from the brain’s surface (0) to the white matter (1); shaded lines, individual mice; thicker lines, average across mice for each age group. The percentage of PV expressing cells (**D**, right, normalized to the DAPI count per section) and mean intensity of MBP staining (**E**, right) was quantified as the average across cortical layers and age groups per mouse. **(F)** Linear regression R^2^ and 95% confidence intervals (dashed lines) for the intensity of MBP expression (average intensity per sample, summed across cortical depth) and number of PV+ cells (average count per sample, summed across cortical depth) per animal (R^2^=0.52, p < 0.001, n=26 mice, younger adult mice [n=11, 2-6 months], older adult mice [n=7, 12-20 months], and mice in old age [n=8, +24 months]). Asterisk indicates significance level of *p < 0.05, **p < 0.01, ***p < 0.001; in **(D, E)** values from Tukey-Kramer corrections.

The human data on coactivated functional inhibition is inconclusive, with no clear evidence for the expected reduced coactivated inhibition in older adults. In mice, we found suppressive influences of the coactivated double stimulation condition in comparison to the single stimulation condition in spatially averaged activity maps (see Fig. 4**-supplemental** figure 1B), although it should be noted that barrel fields were stereotaxically targeted to include the preserved whiskers without precise receptive field mapping (see methods). We observed changes in neuronal responses to whisker stimulation conditions on the single-cell level and in the percentage of responsive neurons across age and cortical layers (see Fig. 4**-supplemental** figure 1). Particularly in the outer cortical layers (layer II/III), younger adult mice show an increased proportion of individual neurons with additive responses (younger: 65%, older: 51%) while older adult mice show a higher proportion of neurons with reduced sensory-evoked responses (younger: 13%, older: 23%) in the double stimulation condition compared to the single stimulation condition, indicating increased inhibitory influence of the neighboring whisker in older adult mice. The more pronounced sensory input signals in older mice as well as increased functional inhibition of neighboring whiskers may help to interpret the human data mechanistically (see Figure 3).

Next, we examined cytoarchitectural changes across layers in the barrel cortex of younger adult (14.8±1.7 weeks, n=17, between 2-6 months) and older adult (68.7±3.8 weeks, n=13, between 12-20 months) mice as well as mice into further old age (114.7±1.1 weeks, n=8, > 24 months; see Figure 4C-E). To clarify the hypothesis that lower qT1 values (indicating higher myelination) in deep layers of older adults could prevent the decrease of coactivated functional inhibition (i.e., ‘altered modulation hypothesis’) and to assess changes specific to inhibitory drive, we examined the proportion of neurons expressing parvalbumin (PV+) in these mice, which is a calcium binding protein and molecular marker of the largest class of inhibitory interneurons in the cortex^19^. We found a significant effect of age driven by increased PV+ density in older mice (F_(2,105)_=21.7, *p* < 0.001) and layer (F_(2,105)_=39.5, *p* < 0.001), but no interaction (F_(4,105)_=1.48, *p*=0.214, two-way mixed-effects ANOVA).

To assess the neurophysiological basis for the finding in humans that cortical aging is associated with lower qT1 values (indicating higher myelination) in middle and deep SI layers, we also examined the intensity of myelin basic protein (MBP) immunohistochemistry staining across cortical layers and age in a subset of these mice (younger adult mice [n=11, 2-6 months], older adult mice [n=7, 12-20 months], and mice in old age [n=8, +24 months]). We found that older mice show increased MBP expression in layer IV and deeper cortical layers, however, in very old age, these cortical layers showed a loss of MBP expression (see Figure 4E; significant effect of age (F_(2,69)_=24.0, *p* < 0.001) and layer (F_(2,69)_=110.6, *p* < 0.001), but no interaction (F_(4,69)_=2.23, *p*=0.075, two-way mixed-effects ANOVA). These results are consistent with previous studies that showed an inverted U-shape relationship between myelination and age^20^. We also found a positive correlation between MBP expression and PV+ density across mice (R^2^=0.52, *p* < 0.001, n=26 mice, see Figure 4F).

Finally, the dynamics of microglia are also known to change with age and it has been shown that myelin debris accumulates within microglia with aging^20,21^. We found significant differences in microglia density (as measured by Iba1 expression) across cortical layers in this cohort of younger adult mice but not in older ages (with an increased microglial density in upper cortical layers in younger adults that was absent in older adults, see Fig. 4**-supplemental** figure 2). However, we found no significant correlation between microglia and MBP expression (R^2^=0.09, *p*=0.120) or PV+ neuron density (R^2^=0.05, *p*=0.278) across animals (Fig. 4**-supplemental** figure 2), suggesting that the specific functional changes we observed across aging and cortical layers are more likely linked to changes in excitatory-inhibitory balance and sensory-related altered modulation rather than specific neuroinflammatory dynamics.

### Relation between layer-specific changes and human sensorimotor impairments

Finally, we investigated whether interindividual variation in layer-specific microstructure and function is related to age-specific functional and behavioral decline in humans. We first confirm that older adults compared to younger adults show worse tactile and motor performance (for exact results see Fig. 5**-supplemental table 1** and Fig. 5**-supplemental table 2**). We then calculated Kendall’s tau correlations for older adults (see Figure 5), where the following correlations show a moderate relationship: (1) lower qT1 values (higher myelin content) in the superficial layer compartment are associated with lower network centrality of the index finger (τ=-0.30), and (2) lower qT1 values (higher myelin content) in middle and deep layer compartments of the index finger representation are associated with worse tactile 2-point discrimination performance of the index finger (τ=-0.26). Please note, however, that these correlations do not exceed the significance level (*p* > 0.1).

**Figure 5.**
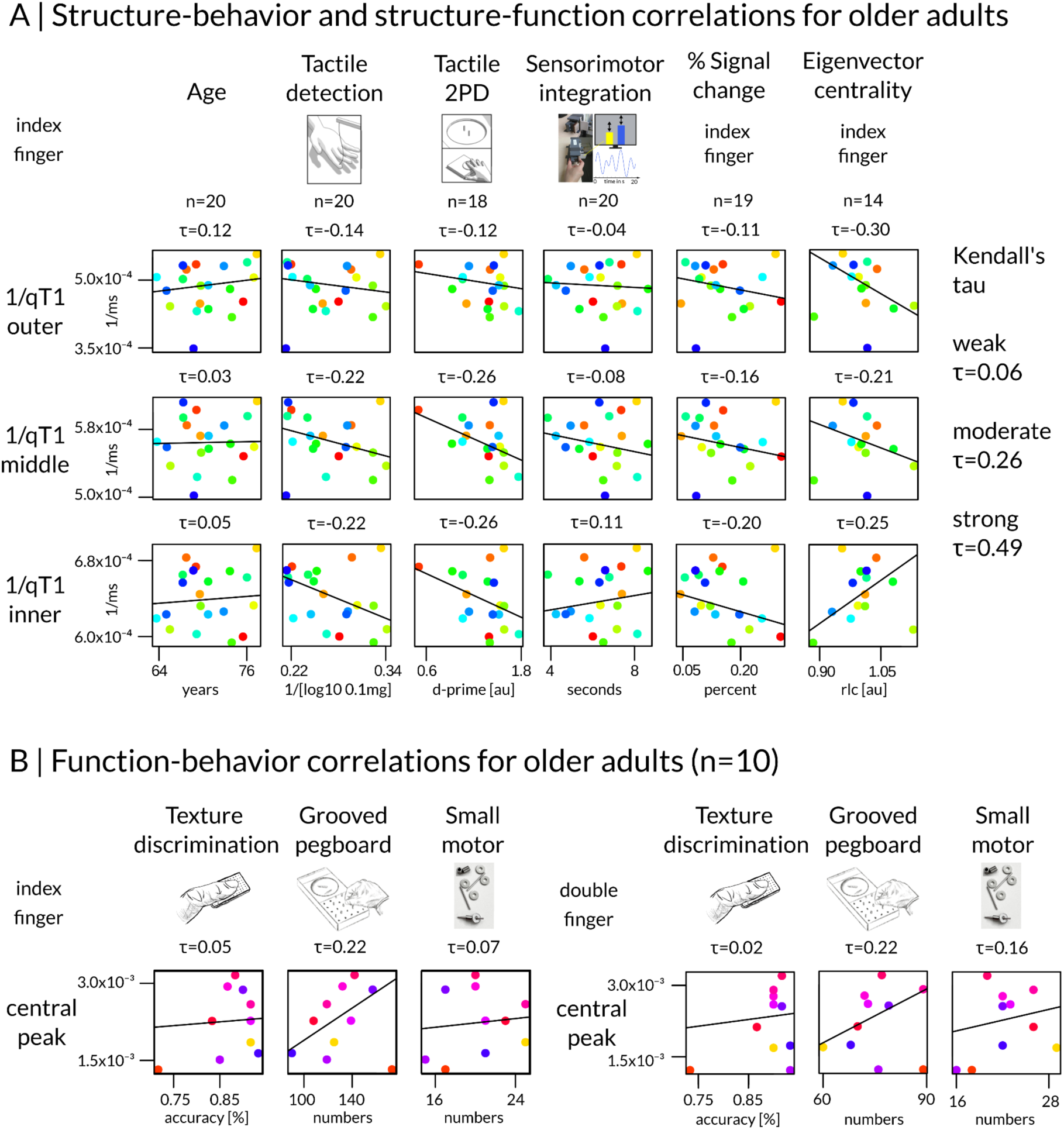
Relation between layer-specific changes and sensorimotor impairments in older adults. **(A)** Structure-function and structure-behavior correlations shown for the index finger representation only. Regression lines were generated based on robust Theil-Sen regression estimates^22,23^. Correlation coefficients are given as Kendall’s tau. Cut-offs for the interpretation of Kendall’s tau were approximated by applying the formula tau=2/π arcsin(r) to cut-offs for the Pearson correlation coefficient (r)^24,25^. To ensure that higher values indicate better performance or more myelin, tactile detection and qT1 values (given in milliseconds) were reversed. Network centrality is given as rectified linear unit correlation in arbitrary units (au), tactile 2-point discrimination performance (tactile 2PD) is given as dprime in arbitrary units (au). There are no significant correlations (uncorrected p-values > 0.1). **(B)** Function-behavior correlations shown for index finger and index-middle finger pair. Correlation coefficients are given as Kendall’s tau. There are no significant correlations (uncorrected p-values > 0.1).

## Discussion

To develop a detailed understanding of alterations in the layer-specific architecture and their associated phenotypes, we studied the layer-specific structural and functional architecture of SI using *in-vivo* ultra-high field 7T-MRI data of two cohorts of healthy younger and older adults. We also investigated SI whisker representations in younger and older mice using *in-vivo* 2PCI in combination with histology to derive more information on potential underlying neuronal mechanisms. Altogether, our data reject the idea that the principal layer architecture of the sensory cortex is preserved with aging. Rather, cortical thinning is driven by deeper layers, and the middle compartment, encompassing layer IV, appears thicker in older compared to younger adults. Whereas a previous study uncovered that primary motor cortex, MI, shows vulnerability in the output layer V and superficial layers to aging^5^, in SI, age-related changes are most pronounced in input and deep modulatory layers. These novel insights (summarized in Figure 6) suggest that layer-specific degeneration is dependent on the specific microstructural architecture of the cortex, and needs to be described separately for different cortical areas.

**Figure 6.**
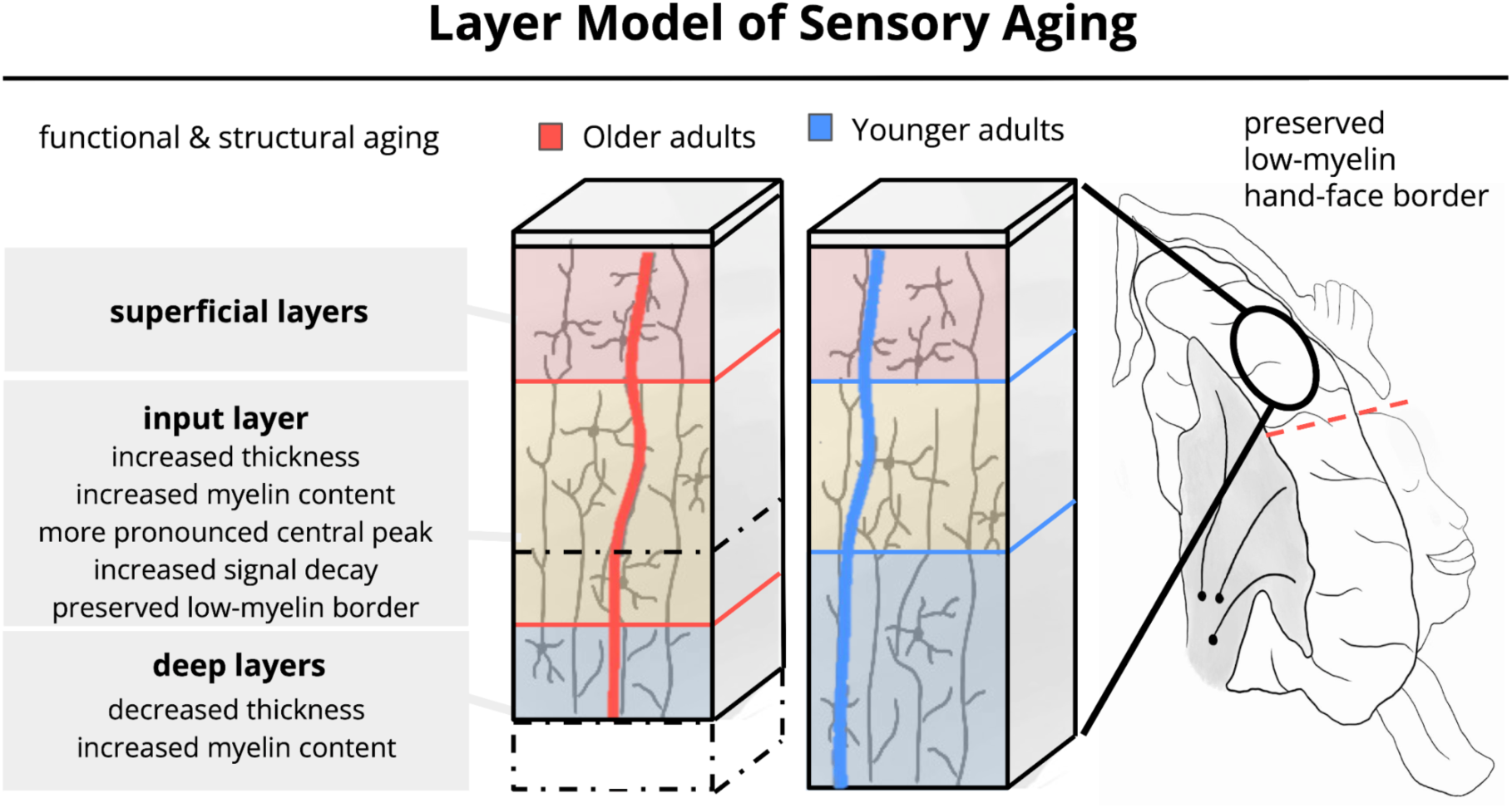
Layer Model of Sensory Aging. The layer model of sensory aging suggests that cortical aging, as one example of cortical dysfunction, has a unique layer-specific profile. Critical characteristics are a widened input channel (shown in cream color), a layer-specific profile of cortical thinning driven by deeper layer degeneration (shown in blue color, note that there is only anecdotal evidence for the null hypothesis of no group difference for the superficial layers (BF10=0.635)), and altered modulatory influences on functional representations. Black dotted lines indicate changes in layer-specific thickness in older adults compared to younger adults. Red and blue vertical lines schematically represent the layer-specific functional input signal for older and younger adults, respectively (with more input to the right, i.e. in x-direction).

Older adults present with a thicker and more myelinated input layer that exhibits a more pronounced sensory signal peak. Older mice also show increased MBP expression in layer IV, however, in more extreme old age, this cortical layer shows a loss of MBP expression, which is congruent with previous findings of changes in myelination across aging in animal models^20^ and cross-species studies^26^, as well as in relation to sensory activation^6,27^. A combination of age- and experience-driven mechanisms influencing the composition of layer IV is a potential explanation for these findings^6^. The more pronounced layer IV in older age may reflect plasticity over the individual lifespan, followed by degeneration in very old age. Layer IV structural alterations may allow individuals to compensate for peripheral signal loss, by shifting short-range connections towards inhibition to sharpen sensory signals and/or stabilize established functional networks^28,29^. This view confirms the ‘altered input channel hypothesis’ specifying a wider input channel in SI of older adults. An alternative view is that separate mechanisms underlie a thicker layer IV in older adults and potential experience-driven changes of layer IV architecture. Possible contributing mechanisms could be changes in synaptic plasticity^30^ and/or architectural differences of oligodendrocytes, microglia, and other neuroglial dynamics with aging^21,31,32^.

Second, we show that age-related cortical thinning in humans is layer-specific. Therefore, the usual across-layer estimate of cortical thickness, often used to describe the human cortex in health and disease, may not be as informative as previously assumed. This is true, in particular, given the middle compartment of SI is thicker despite overall cortical thinning in older adults. Layer-specific computations of cortical thickness therefore derive mechanistic information regarding the architecture of the cortex, such as plasticity-related expansion and age-related degeneration. Calculating the average across layers disregards this critical intracortical information.

Deeper layers are thinner but more myelinated in older adults. At the same time, the spatial selectivity of the index finger representation is coarser while there is no clear evidence that coactivated functional inhibition is reduced in older adults. These measures may be connected given deeper layers in SI are involved in subcortical signal integration, signal modulation, and inhibition^33,34^. Whereas the BOLD effect has limitations with respect to its interpretation as excitatory or inhibitory^33^, we replicate sensory-driven overactivation and found increased functional inhibition in older mice using single-neuron activity. In addition, we found higher PV+ cell density in older mice. Previous work in mice has shown that a large fraction of myelin in cortical layers II/III and IV ensheaths axons of GABAergic interneurons, particularly of PV+ basket cells in SI^34^. In humans, PV+ interneurons contribute substantially to the total myelin content in the cerebral cortex^35^. One explanation is to assume that the decreased qT1 signal reflects changes in myelination of cortical output with axons oriented through the deeper layers to enter long-range white matter tracts as well as an increase in PV+ density. Substantial evidence, and results from the current study in mice, show that cortical myelination continues throughout adulthood and does not begin to decline until well into old age^20,26,36^. This increase in myelination may also be linked to an increase in sensory-evoked activation and subsequent PV+ cell density, which may act as compensatory mechanisms for maintaining excitatory-inhibitory balance^37,38^. Thinner deep layers, on the other hand, may reflect processes of reduced overall cell density that are proceeding at these stages of age, and may even contribute to overactivation due to changes in local circuit dynamics and reduced inhibition. This may provide a mechanistic explanation for the altered sensory-evoked functional responses in SI in older humans and mice, and would support the ‘altered sensory modulation hypothesis’.

It is worth noting that in older compared to younger adult mice, PV+ cell density is increased as a main effect in all layers (most pronounced in layer IV), whereas in humans, increased myelination was restricted to middle and deep layers. If our interpretation of the ‘altered modulation hypothesis’ is correct, this suggests that cortical layer inhibition is differently affected in humans and mice with aging. An alternative explanation for the qT1 effect is that myelin sheaths are degrading (i.e., becoming less compact) due to sheath splitting and the formation of myelin balloons, which is accompanied by the production of redundant myelin and an increase in oligodendrocytes in older age^31^. Future studies focusing on cytoarchitectural changes in relation to changing myelin patterns, neurovascular coupling and sensory processing can address this, of particular interest would be cross-species longitudinal studies. Whereas longitudinal rs-MRI studies have shown changes in functional connectivity of somatosensory networks with age in mice^39^, detailed MRI approaches to assess layer IV microstructural changes^40^ have yet to be investigated in mouse models using a longitudinal approach across aging.

The present findings have multiple clinical implications. Our study reveals that the combined investigation of layer-specific thickness, microstructural layer compositions and layer-specific functional readouts, and the parallel investigation of humans and mice, uncovers key aspects of cortex organization and potential dysfunction. In Alzheimer’s Disease (AD), cortical thinning is related to cell loss and regarded as an early marker of the disease^41^. Whereas we show specific degradation of deep layers with aging, superficial layers may be altered earlier in AD. In multiple sclerosis (MS), where neuroinflammation causes myelin damage in specific cortical layers^42^, post-mortem investigations in humans and mice showed a decrease in PV+ interneurons^43,44^, suggesting an interaction between layer-specific myelination and excitation/inhibition modulation. Losing myelin progressively can result in epilepsy-like brain activity, as inhibition of slow brain waves decreases^45^, and epileptogenic hyperexcitability and lesions often present layer-specific^46^. Multimodal layer-specific investigations could uncover underlying mechanisms, also in MS^42^. Investigating participants with congenital or acquired hand loss may provide further insights into the mechanisms of layer-specific plasticity in SI, specifically to disentangle age-related from input-related changes. While we report a case study of a healthy, middle-aged participant with congenital arm loss in the **Supplemental Information**, extended and thorough investigations of this special population may derive key insights into underlying mechanisms. In addition, further studies with higher sample sizes will allow investigating subtle differences between age groups in more detail. The results presented here motivate the investigation of high-precision cortical pathology, allowing for the tailoring of interventions to specific patients.

## Methods

### General procedure

Human data acquisition and analyses were organized in two cohorts: cohort 1 and cohort 2. In Fig. 1**-supplemental table 1**, a detailed overview over which data was used for which specific analyses is provided.

In brief, the data of cohort 1 was used to derive the insights reported in the sections ‘Age-related cortical thickness changes are layer-specific’, ‘Low-myelin borders have a similar architecture in younger adults, older adults, and with congenital arm loss’, ‘Altered microstructural layer composition in older adults’ and ‘Relation between layer-specific changes and human sensorimotor impairments’. Younger and older adults of cohort 1 attended altogether five experimental sessions (numbered 1-5): (1) structural 7T-MRI scanning session for layer-specific structural analyses (see **Table 1** and Figure 2 for results), (2) functional 7T-MRI scanning session including both the blocked-design and phase-encoding paradigms where the fingertips were stimulated (design and analyses reported under the heading ‘*Functional localizer task: Individual fingers*’), (3) functional 7T-MRI scanning session with hand, face, and foot movements ^5,10^ (design and analyses reported under the heading ‘*Functional localizer task: hand and face areas*’), (4) behavioral testing for finger discrimination task (results shown in Fig. 5**-supplemental table 1**), and (5) behavioral testing for tactile 2-point discrimination task, tactile detection task, sensorimotor integration task (results shown in Figure 5 and Fig. 5**-supplemental table 1**).

The data of cohort 2 was used to derive the insights reported in the sections ‘More pronounced sensory input signals in layer IV in older adults’, ‘Altered functional response profile in older adults’ SI’ and ‘Relation between layer-specific changes and human sensorimotor impairments’. Younger and older adults of cohort 2 attended four sessions (numbered 6-9): (6) combined structural and functional 7T-MRI scanning session with blocked-design paradigm where the full index and middle fingers were stimulated, and used as localizer for the layer-specific functional analyses (result shown in Figure 3), (7) functional 7T-MRI scanning session with a phase-encoding paradigm where the full index and middle fingers were stimulated (design and analyses reported under the heading ‘*Functional task used for layer-specific analyses*’, see Figure 3 for results), (8) structural 3T-MRI scanning session for individual cortical surface reconstruction, and (9) behavioral measurement (including tactile detection task, texture roughness discrimination task and hand dexterity tasks, results shown in Figure 5 and Fig. 5**-supplemental table 2**, see Figure 1 for experimental design and analysis pipeline).

### Human participants

#### Younger and older adults

Cohort 1 was composed of 46 healthy volunteers who underwent structural 7T MRI, functional localizers of individual fingers, as well as behavioral tests of tactile finger performance. Due to severe motion artifacts in the imaging data, 6 participants were excluded from the study, leaving a total of 40 participants for analyses of cohort 1 (20 younger adults: 10 females, 21-29 years, Mean±SD: 25.1±2.7 years; and 20 older adults: 10 females, 63-77 years, Mean±SD: 70.5±4.0 years).

Cohort 2 was composed of 21 healthy, right-handed younger adults and 17 healthy older adults who underwent behavioral measurements, of which, 14 younger adults and 13 older adults underwent 7T fMRI with a detailed investigation of the index and middle finger. Due to severe head motion during scanning, 3 younger adults and 3 older adults had to be excluded, leaving 11 younger adults (5 female, 25-35 years, Mean±SD: 28.18±3.06 years) and 10 older adults for analyses of cohort 2 (3 female, 60-80 years, Mean±SD: 68.40±6.20 years).

Participants of both cohort 1 and cohort 2 were right-handed. In cohort 1, this was investigated via the Edinburgh handedness questionnaire^47^ (laterality index ranging from +40 to +100; Mean±SD: 83.93±19.56). In cohort 2, this was investigated in a semi-structured interview (i.e., excluding left-handed and ambidextrous candidates). Chronic illness, central acting medications and MRI contraindications (e.g., active implants, non-removable metallic objects, tattoos, claustrophobia, tinnitus, consumption of alcohol and drugs, and pregnancy) were a priori exclusion criteria for both cohorts. A study-specific health screening revealed no anomalies of sensory perception (e.g., numbness, tingling sensations, hypersensitivity, hyperalgesia) and motor movement (e.g., loss of motor control, restricted finger movement) in distal extremities. Given enhanced cortical hand representations and superior tactile perception in string and piano players^48^, no professional musicians were enrolled in either cohort. We note that 2/20 older participants of cohort 1 underwent successful carpal tunnel surgery on either the right hand only or on both hands. However, their tactile performance was absent of clinical signs (no outliers detected in the data), leading us to keep the data in the analysis. Besides, participants reported no other medical conditions. Finally, none of the participants of cohort 1 showed signs of cognitive impairments as indicated by the ‘Montreal Cognitive Assessment’ (MoCA; score ≥ 26 used as criterion for healthy aging; younger adults Mean±SD: 29.0±1.1; older adults Mean±SD: 27.8±2.2)^49^, except for one older adult with a MoCA score of 21. Because this participant performed equally well in the behavioral tasks compared with all other older adults, we kept the data in the analysis.

Younger and older adults from both cohorts were recruited from the database of the German Center for Neurodegenerative Diseases (DZNE) Magdeburg, Germany. All participants gave written informed consent and were paid for attendance. The study was approved by the Ethics committee of the Otto-von-Guericke University Magdeburg. Data of younger and older adults of cohort 1 were partly published in previous studies^3,10^.

#### Participant with congenital arm loss

We also collected data of one healthy adult with congenital arm loss (male, age: 52 years, affected side: right, level of deficiency: complete arm missing from the shoulder downwards, no experience of phantom sensations and pain, cosmetic prosthesis worn > 8 hours per day, prosthesis never involved in daily life routines).

The participant with congenital arm loss was recruited from the database of the Central Institute of Mental Health (CIMH) Mannheim, Germany. He gave written informed consent and was paid for his attendance. The study was approved by the Ethics committee of the Otto-von-Guericke University Magdeburg.

### Human MR scanning

#### Human 7T MRI

We used a whole-body 7-Tesla MAGNETOM scanner (Siemens Healthcare, Erlangen, Germany) equipped with a 32 Channel Nova Medical head coil to acquire MP2RAGE^50^ images with whole brain coverage for both cohorts (session 1 and session 6, see Fig. 1**-supplemental table 1** for session indexing) (0.7 mm isotropic resolution, 240 sagittal slices, FoV read=224 mm, TR=4800 ms, TE=2.01 ms, inversion time TI1/TI2=900/2750 ms, flip angle (α)=5°/3°, bandwidth=250 Hz/Px, GRAPPA 2). In addition, for cohort 1 (session 1), we acquired MP2RAGE images with part brain coverage (targeting the sensorimotor cortex; 0.5 mm isotropic resolution, 208 transversal slices, FoV read=224 mm, TR=4800 ms, TE=2.62 ms, inversion time TI1/TI2=900/2750 ms, flip angle (α)=5°/3°, bandwidth=250 Hz/Px, GRAPPA 2, phase oversampling=0%, slice oversampling=7.7%), and susceptibility-weighted images with part brain coverage (targeting the sensorimotor cortex) using a 3D gradient-recalled echo (GRE) pulse sequence^51^ (0.5 mm isotropic resolution, 208 transversal slices, FoV read=192 mm, TR=22 ms, TE=9.00 ms, flip angle=10°, bandwidth=160 Hz/Px, GRAPPA 2, phase oversampling=0%, slice oversampling=7.7%). Structural scanning lasted in total approximately 60 minutes in session 1 and about 20 minutes in session 6 (because no 0.5 mm resolution images were acquired).

To acquire fMRI data, we first performed shimming and acquired two echo-planar images (EPIs) with opposite phase-encoding (PE) polarity before the functional time-series were collected using GRE EPI pulse sequences (cohort 1, sessions 2 and 3: 1 mm isotropic resolution, FoV read: 192 mm, TR=2000 ms, TE=22 ms, GRAPPA 4, interleaved acquisition, 36 slices; cohort 2, session 6 and 7: 0.9 mm isotropic resolution, 30 slices, interleaved acquisition, FoV read=216 mm, TR=2000 ms, TE=22 ms, GRAPPA 4). The same sequence was used for all functional tasks (see below).

#### Human 3T MRI

For cohort 2 (session 8), 3T MRI data were acquired at the Philips 3T Achieva dStream MRI scanner, where a standard structural 3D MPRAGE was acquired with the following parameters: resolution: 1.0 mm, 192 slices, FoV read=192 mm×256 mm, slab thickness=256 mm, TI=650 ms, echo spacing=6.6 ms, TE=4.73 ms, flip angle=8°, bandwidth=191 Hz/Px.

#### Human physiological data recording

A pulse oximeter (NONIN Pulse Oxymeter 8600-FO) clipped to the participant’s left index finger (no stimulation module attached) captured the pulse, and a breathing belt captured respiration during fMRI (cohort 1: session 2; cohort 2: session 6 and 7). Signals were digitally recorded with a sampling frequency of 200 Hz using an in-house developed setup (National Instruments USB 6008 module with pressure sensor Honeywell 40PC001B1A).

### Human fMRI tasks

#### Functional localizer task: Individual fingers

In session 2 of cohort 1, we applied a previously established tactile stimulation paradigm to localize the fingers of the right hand in contralateral SI (localizer used for cortical thickness analyses, microstructural composition analyses, low-myelin border analyses, structural topography analyses)^3,10^. During the tactile task, five independently-controlled MR-compatible modules were used to stimulate the five fingertips of the right hand (each with 8 piezoelectric-controlled pins arranged in a 2×4 matrix, covering 3.5×8.5 mm^2^ of skin; Quaerosys, http://www.quaerosys.com, see Figure 1)^3^. Individually-adjusted vibrotactile stimulation was applied at a frequency of 16 Hz using 16 pin combinations per second^3^, raising 2 pins at a time. First, a phase-encoded protocol was applied (2 runs of 20 cycles; each fingertip stimulated 20 times for 5.12 seconds) in forward (thumb to little finger, 50% forward-run first) and reverse order (little finger to thumb, 50% reverse-run first). One run took 8 minutes and 32 seconds (256 scans, TR of 2 seconds). Participants counted randomly occurring stimulation pauses and had to report them after each run. A blocked-design protocol was used to stimulate the fingers in a pseudo-random way (2 runs; 6 conditions: stimulation to thumb, index, middle, ring, little finger and no stimulation), using the same instructions and stimuli as described for the phase-encoded protocol. One run took 6 minutes and 56 seconds (each fingertip was stimulated 10 times for 2 seconds followed by a 22 seconds resting phase; inter-stimulus intervals of 2 seconds in 70% of trials or 6 seconds in 30% of trials were counterbalanced between fingers; 208 scans). Finally, we acquired resting-state data in a 5-minute scan while participants looked at a fixation cross and were asked to think about nothing in particular. Total scan time was approximately 40 minutes.

#### Functional localizer task: hand and face areas

In session 3 of cohort 1, younger and older adults (n=34) as well as the participant with congenital arm loss underwent the same motor paradigm as described in Northall et al.^5^ to locate the hand and face areas in BA 3b (used to investigate cortical thickness in major body part representations, and low-myelin borders between the hand and the face). In short, we used a blocked-design paradigm where participants carried out motor movements of the left and the right hand, the left and the right foot (investigation not part of this study, but used in Northall et al.^5^) and the tongue. Participants were trained outside the MR scanner and wore fingerless braces to stabilize the hand while carrying out precise movements. Movements were carried out for 12 seconds followed by a 15 seconds rest period. Movements were repeated four times each resulting in 20 trials and taking approximately 9 minutes in total. For the one-handed adult, we applied a modified version of the original motor paradigm using mental imagery of finger movements for the missing limb.

#### Functional tasks used for layer-specific analyses

In session 6 and 7 of cohort 2, the same independently-controlled MR-compatible stimulation modules as described above for session 2 were used to stimulate the distal phalanges and intermediate phalanges of the index and middle finger of the right hand. One module was attached to each phalanx using a custom-build, metal-free applicator that fitted individual finger sizes (see Figure 1A). The vibrotactile stimulation was applied at a frequency of 16 Hz, and the stimulation intensity of each finger was adjusted to two times of the individual tactile detection threshold of the respective finger. To minimize adaptation-related differences in map activity between individuals, three randomly chosen pins were raised once at a time, yielding 16 pin combinations per second. In session 6, participants underwent two blocked-designed runs to localize the distal and intermediate phalanges of the index and middle finger. For each run, participants were asked to look at a centrally presented fixation cross. The blocked-design run comprised three conditions, including stimulation of the index finger, the middle finger, and a rest condition with no stimulation. Each finger was stimulated for 8 seconds in a pseudo-random sequence, where one finger was stimulated maximally two times in a row. In 70% of the trials, there was a 4 seconds pause between two subsequent stimulations, in 30% of the trials, there was a 8 seconds pause between two subsequent stimulations. This was counterbalanced across fingers. Each finger was stimulated 20 times. One run comprised 264 scans, and lasted for 8 minutes and 48 seconds. The blocked-design run was repeated twice, lasting around 20 minutes in total.

In session 7, participants underwent twelve phase-encoded runs, which were used for % signal change and pRF modeling analyses. The phase-encoded runs included three different conditions: (1) stimulation of only the index finger, (2) stimulation of only the middle finger, (3) stimulation of both the index and the middle finger. Each condition comprised four runs, each consisting of 8 stimulation cycles and two rest conditions of 32 seconds (one before and one after stimulation). Each stimulation cycle lasted 32 seconds, and stimulation was applied to each section of the phalanx four times for 8 seconds. Half of the stimulation runs of each condition were delivered in a forward order (finger top→ finger bottom) and the other half in a reverse order (finger bottom→ finger top) (see Figure 1A). Half of the participants of each age group started with the forward order-run, the other half started with the reverse order-run. One run comprised 160 scans (128 scans for stimulation and 32 scans for rest), lasting 320 seconds (TR=2 seconds). Participants were instructed to covertly count short randomly distributed interrupts embedded in the tactile stimulation (duration 180 ms, see Liu et al.^3^). There were the same number of gaps in each run (32 gaps in total). All phase-encoded runs took around 60 minutes.

### Human MRI analyses

7T structural data processing for layer-specific cortical thickness, microstructural composition, low-myelin border and structural topography analyses (cohort 1)

### Reconstruction of QSM images

In cohort 1 (session 1), quantitative susceptibility mapping (QSM) data were reconstructed from susceptibility-weighted images (i.e., magnitude and phase images, collected in session 1 using a 3D GRE pulse sequence) using the Bayesian multi-scale dipole inversion (MSDI) algorithm^52^ as implemented in QSMbox (version 2.0, freely available for download: https://gitlab.com/acostaj/QSMbox). No normalization was applied to QSM values, because aging effects are assumed to be similar for normalized and non-normalized 7T MRI data^16^.

### Image registration and cortex segmentation for layer-specific structural analyses

Two independent raters evaluated structural data quality of QSM and qT1 images (with the effect of excluding data of n=6 participants with severe motion artifacts) which were collected in session 1 of cohort 1. Only data showing no artifacts or mild truncation artifacts (not affecting SI) were processed. We used the same structural image preprocessing pipeline as described previously^10^, employing CBS Tools (v3.0.8) as a plugin for MIPAV (v7.3.0). In short, we co-registered qT1 slab images to upsampled qT1 whole brain images (combining linear and non-linear registration using ANTs version 1.9.x-Linux, embedded in CBS Tools) in one single step (nearest neighbor interpolation). To register QSM slab images to qT1 images, a combination of rigid and affine automated registration was applied using ITK-SNAP (v3.8.0). Registered qT1 slab and upsampled qT1 whole brain images were fused. Resulting images were skull stripped and dura removed. Dura estimates were manually refined where required to ensure complete removal of non-brain matter from the region of interest. We used the TOADS algorithm^53^ to segment the cortex from the rest of the brain before we applied the CRUISE algorithm^54^ to estimate tissue boundaries using the level set framework^55^. Cortex estimates were thresholded between the maximum of the inner and outer level set images (−2.8 and −0.2, respectively). Afterwards, the cortex was divided into 21 cortical depths using the validated equivolume model^56^. Intra-cortical qT1 values were (used as proxy for myelin)^12^ and QSM values (used as proxies for iron [positive QSM values, pQSM]^15^; calcium/diamagnetic contrast [negative QSM values, nQSM]^57^; and mineralization [absolute QSM values, aQSM]^16^) were sampled along the extracted cortical depths in reference to individual cortical folding patterns. Individual cortical surfaces were estimated based on level set images of the middle cortical depth. Layer-specific quantitative values of the non-merged high-resolution qT1 and QSM slab images were mapped onto the inflated cortical surfaces (method of closest point)^58^.

### Extracting microstructural profiles and cortical layer compartments

For cohort 1, layer-specific qT1, nQSM, pQSM and aQSM values were sampled along 21 cortical depths perpendicular to the cortical sheet. Sampled values were extracted from the five fingertip representations (pRF center location maps), and from the face-hand representation (hand and face activation maps; see section *Localizing body part representations in area 3b for layer-specific structural analyses*) in area 3b. Layer-specific quantitative values were averaged across vertices within individuals (giving one value per body part and depth). First and second derivatives of layer-specific qT1 profiles were calculated using the gradient function as implemented in MATLAB (R2017b).

The layer-specific structural profiles (including 21 cortical depths) were then further averaged based on a previously introduced approach^10^, using *ex-vivo in-vivo* validated myelin profiles of area 3b^12^ to identify anatomically-relevant layer compartments in *in-vivo* MRI data (see Figure 2B and D). In short, after removing the two deepest layers (where qT1 stabilized) in both age groups, minima and maxima of the first derivative of raw qT1 profiles indicated three data-driven layer compartments: an inner, middle and outer compartment. Based on Dinse et al.^12^, we assume that the input layer IV is located in the middle compartment and deep layers V/VI are located in the inner compartment. However, we note that these compartments are based on *in-vivo* MRI data and may therefore not match exactly with the anatomical layers as described by e*x-vivo* myelo- and cytoarchitecture.

### Definition of area 3b for layer-specific structural analyses

For cohort 1, area 3b was manually delineated based on an operational definition using anatomical landmarks extracted from cytoarchitectonic^59^, fMRI^60^ and multimodal parcellation studies^61^, i.e., following a standardized procedure that has been used previously^5,10,62^. Resulting masks cover the anterior wall of the postcentral gyrus (mainly covering area 3b and parts of area 3a). Finally, all masks were plotted in reference to co-registered Freesurfer labels (normalized probabilistic maps of area 3a and area 3b) on the individual cortical surfaces to ensure that the locations of the manual delineations overlap with those outlined by automated approaches (see Fig. 2**-supplemental** figure 4).

### Localizing body part representations in area 3b for layer-specific structural analyses

For cohort 1, pRF center location maps (restricted to area 3b) resulting from the tactile stimulation paradigm (session 2, see section *Functional data preprocessing for layer-specific cortical thickness, microstructural composition, low-myelin border and structural topography analyses* for details on pRF modeling) served as localizer for individual finger representations (used for cortical thickness and layer-specific structural composition analyses). A “winner-takes-it-all” approach was applied to sample vertices only once. Overlapping vertices (introduced by mapping splitted single finger pRF center location maps) were exclusively assigned to the finger map with the highest variance explained (obtained from pRF modeling). Combining the five single finger maps to one ROI defined the hand area in area 3b of younger and older adults.

Hand and face activation maps resulting from the motor paradigm (cohort 1, session 3) were used to localize the hand-face area in younger and older adults and in the participant with congenital arm loss (layer-specific cortical thickness, low-myelin border and structural topography analyses). A “winner-takes-it-all” approach was applied to the extracted values to sample vertices only once. For the participant with congenital arm loss, the lowest 30% of t-values were removed to ensure comparable map sizes within and between participants. To exclude spatial outliers, vertices located more than two standard deviations away from the location of the main cluster (in z-direction and in y-direction) were removed from the final data.

### Registration and surface mapping of functional data for layer-specific structural analyses

For cohort 1, all functional data were registered to the qT1 image using the automated registration tool in ITK-SNAP (version 3.6.0, non-rigid transformation, 9 degrees of freedom). Manual refinement was applied where required to ensure registration accuracy. Registration matrices were applied to statistical maps (i.e., t-maps, pRF maps, Eigenvector centrality maps) in a single step (ANTs, version 2.1.0, nearest neighbor interpolation). We note that all ROI analyses were performed on non-smoothed functional data (in original individual space) before statistical maps were registered to individual structural data space. Area 3b masks were applied to all functional data. Registered individual functional parameter maps (e.g., t-value maps, pRF maps, EC maps) were mapped onto the individual cortical surfaces (method of closest point). The most superficial 20% of cortical values were excluded (to account for superficial veins affecting the BOLD signal). The mean of the remaining cortical values (covering 20-100% of cortical depth) were used to compute statistics independent of cortical depth.

### Cortical thickness estimation

For cohort 1, the profile geometry module from the CBS Tools (v3.0.8) for MIPV (v7.3.0) was used to calculate overall cortical thickness across cortical depths (after removing the two deepest cortical depths where qT1 values stabilized) and layer-specific cortical thickness of extracted layer compartments.

### Automatic detection of low-myelin borders

To investigate low-myelin borders in human area 3b in cohort 1, a multimodal surface-based mapping approach was applied to each individual dataset^10^. First, multidimensional sampling (inferior to superior, anterior to posterior) of layer-specific qT1 values was performed within area 3b (Pyvista implementation of the Dijkstra algorithm^63^). The peak activation of the thumb (as identified by blocked design data) served as seed region to sample geodesic paths in inferior-to-superior direction, connecting the upper face representation with the little finger representation. Start and end points were defined along the y-axis of D1 and D5 activation peaks and 10 mm (geodesic distance on shortest path) below the D1 activation peak (estimation based on the location of the forehead as described previously^64^). Considered vertices were scattered within one vertex-to-vertex distance of appr. 0.28 mm around the y-axis. Only in cases where the underlying qT1 pattern did not match y-axis sampling, start and end points were defined along the x-axis. Five equally-distant geodesic paths were sampled for each participant (see Fig. 2**-supplemental** figure 2A) to extract qT1 values from middle cortex depth (where the detection of low-myelin borders in area 3b is expected based on previous findings^9^). Second, peak detection was performed on the five extracted qT1 signals (see Fig. 2**-supplemental** figure 2A). To control for a possible gradient in cortical myelin content along the inferior-to-superior axis^9^, all qT1 signals were detrended before the find_peaks algorithm from the SciPy signal processing toolbox (version 1.10.1) was applied to find the most prominent peaks (local maxima with a prominence > 2 SD from the mean of absolute detrended qT1 values) in each detrended qT1 signal by comparing neighboring values. Resulting peaks were considered a low-myelin border (reflected by a row of high qT1 values) when they occurred in at least 3/5 detrended qT1 signals based on a nearest neighbor approach. The nearest peaks on first and second neighbor signals were grouped together based on geodesic distances. Peaks with a geodesic distance < 5 mm were considered near. Third, resulting low-myelin borders were back-projected to individual cortical surfaces and visualized in reference to individual pRF finger maps to categorize low-myelin borders in hand-face borders and within-hand borders (i.e., by visual inspection). Finally, a number of different features were extracted for each peak including prominence, full width at half maximum, qT1 intensity, Eigenvector centrality, signed QSM intensity, nQSM intensity, pQSM intensity and aQSM intensity. For each feature, all values belonging to a single border were averaged to obtain one feature vector for each border. Feature vectors of within-hand borders were further averaged across all within-hand borders. In this way, we extracted two feature vectors (one for within-hand borders and one for the hand-face border) in each individual which were used to calculate group statistics (between younger and older adults).

### Blocked design analyses to locate individual fingers and hand and face in area 3b for layer-specific structural analyses

For cohort 1, opposite polarity (PE) EPIs were distortion-corrected using point spread function (PSF) mapping^65^. We applied a weighted image combination to combine the distortion-corrected PE EPIs while controlling for differences in spatial information. Resulting EPIs were motion corrected to time point zero. PSF mapping was performed on motion-corrected EPIs to allow geometrically accurate image reconstruction. Evaluating quality of blocked-design and phase-encoded time series, there were no severe artifacts. Resulting data were slice-time corrected using SPM8 (Statistical Parametric Mapping, Wellcome Department of Imaging Neuroscience, University College London, London, UK). Slice-time corrected data of phase-encoded protocols were concatenated.

We used tactile stimulation to localize the finger representation of the right hand in contralateral SI (session 2, used for low-myelin border detection). We calculated 1st level fixed-effects models for the two blocked-design runs using a general linear model (GLM) on individual data as implemented in SPM 8. BOLD activation driven by each finger’s stimulation was modeled as an independent measure, because each finger was treated individually^66^. Five regressors of interest (stimulation to thumb, index, middle, ring, little finger) were modeled per session, resulting in five linear contrasts (e.g., the contrast [-1 4 −1 −1 −1] for stimulation to index finger). Peak clusters of t-values were extracted (*p* < 0.01, minimum cluster size of k=3). Resulting t-value maps were taken forward for surface mapping (see section *Registration and surface mapping of functional data*).

For the motor paradigm (session 3), we used the GLM as implemented in SPM 12 to estimate functional activation maps (t-value maps) of tongue and finger movements (first-level analysis using contrast estimates for each body part, layer-specific cortical thickness, low-myelin border and structural topography analyses). We thresholded peak clusters at *p* < 0.01 with a minimum size of k=3 (for details see Northall et al.^5^). Resulting t-maps were taken forward for surface mapping). In the participant with congenital limb loss, we used mental imagery of finger movements to locate the hand area in area 3b in the hemisphere contralateral to the missing limb, while the rest of the motor paradigm was equal to the one used for younger and older participants. Again, we used the GLM as implemented in SPM 12 to estimate t-value maps based on contrasts for each body part. We note that the motor paradigm also included movements of the left and right foot (which were not investigated here), resulting in five different contrasts (e.g., the contrast [1 0 0 0 0] for moving/imagining movements of the left hand). Resulting t-maps were taken forward for surface mapping and were thresholded in surface space.

### Population receptive field modeling to locate the fingertips of the right hand in contralateral area 3b for layer-specific structural analyses

For cohort 1, we used the same method as described previously to estimate Bayesian population receptive fields (pRF) of the five fingertips^3,10^. In short, phase-encoded fMRI data (collected in session 2) were used to perform a two-stage analysis using the BayespRF Toolbox (freely available for download from https://github.com/pzeidman/BayespRF) written for Matlab (SPM12, Matlab R2017b): First, we conducted a 1st level GLM analysis, constructing five regressors for the five fingers of the right hand. Only voxels passing a threshold of *p* < .05 uncorrected were included in pRF modeling^3^, which was performed on a voxel-by-voxel basis on the inferior-to-superior dimension (x-dimension) of topographic alignment, using a Gaussian response function and a posterior model probability > 0.95 (code available for download at https://gitlab.com/pengliu1120/bayesian-prf-modelling.git)^3^. The analysis was restricted to area 3b to reduce processing time. We extracted pRF center location (x) to locate activated finger-specific voxels (finger-specific ROIs) and pRF width (σ, standard deviation of the Gaussian response function) to estimate the pRF size of activated voxels in one-dimensional stimulus space. The most superficial 20% of the remaining data points were disregarded to minimize the effects of superficial veins.

### %signal change analysis for structure-function correlation analyses

For cohort 1, statistical analyses were conducted on the averaged individual time series of the averaged forward- and reversed-order runs from the phase-encoded paradigm (session 2)^3,66^ to calculate mean response amplitudes, using the program Fourier as implemented in csurf (http://www.cogsci.ucsd.edu/~sereno/.tmp/dist/csurf). Discrete Fourier transformations were performed on the time course of each 3D voxel, before calculating the phase and the significance of the periodic activation. Cycles of 20 stimulations were used as input frequencies. Frequencies below 0.005 Hz (known to be dominated by movement artifacts) were excluded, while higher frequencies up to the Nyquist limit (1/2 the sampling rate) were included. For display, a vector was generated whose amplitude was the square root of the F-ratio calculated by comparing the signal amplitude at the stimulus frequency to the signal amplitude at other noise frequencies, and whose angle was the stimulus phase. To estimate mean response amplitudes of the five finger ROIs (in %), we estimated the discrete Fourier transform response amplitude (hypotenuse given real and imaginary values) for each voxel, within each finger’s pRF center location area. This value was multiplied by two to account for positive and negative frequencies, again multiplied by two to estimate peak-to-peak values, divided by the number of time points over which averaging was performed to normalize the discrete Fourier transform amplitude, and divided by the average brightness of the functional data set (excluding air). Finally, the value was multiplied by 100 to estimate the percentage response amplitude^3,66^. No spatial smoothing was applied to the data. The data was sampled onto the individual Freesurfer surface for each participant. To minimize the effect of superficial veins on BOLD signal change, superficial points along the surface normal to each vertex (upper 20% of the cortical thickness) were disregarded. The individual maps were calculated based on the mean value of the remaining depths (20-100% cortical depth). Clusters that survived a surface-based correction for multiple comparisons of *p* < .05 (correction was based on the cluster size exclusion method as implemented by surfclust and randsurfclust within the csurf FreeSurfer framework)^67^ and a cluster-level correction of *p* < .001 were defined as significant. For each participant and condition, the complex-valued phasing-mapping data (real and imaginary values) was sampled onto the individualized inflated 3D cortical surface, and the values within the ROI were extracted. Mean % signal change of the tactile maps of each condition were calculated.

### Resting-state data processing for structure-function correlation analysis

For cohort 1, resting-state functional data (collected in session 2) were corrected for pulse- and respiration-induced noise. To prepare the physiological data for noise correction and to remove acquisition artifacts, we used the open-source Python-based software ‘PhysioNoise’^68^. Resulting respiratory and cardiac phase data were then used to correct the resting-state time series for pulse- and respiration-induced noise by performing RETROspective Image CORrection (RETROICOR) on a slice-by-slice basis^69^. Residuals were taken as cleaned data to regress out motion-related noise parameters (extracted from the raw data) using the program vresiduals as implemented in LIPSIA (freely available for download at <u>github.com/lipsia-fmri/lipsia</u>)^70^. The resulting data were high-pass filtered at 0.01 Hz (allowing frequencies faster than 0.01 Hz to pass) and smoothed (Gaussian kernel with a FWHM of 2 mm) using the program vpreprocess implemented in LIPSIA. For n=6 participants, physiological data were not successfully recorded due to loosening the pulse oximeter and/or breathing belt during scanning, which interrupted successful data sampling. For n=5 participants, severe motion artifacts were detected in the resting-state data. Therefore, resting-state analyses are presented for n=29 (14 younger, 15 older) participants only.

Eigenvector centrality (EC) maps were calculated in native space as a measure of network centrality (i.e. maps reflecting the degree of connectedness of nodes within a network^71^) using the program vecm, implemented in LIPSIA^70^. Thereby, the method of rectified linear unit correlation (RLC)^72^ was applied, which is suitable for high-resolution fMRI data.

### Structural data processing for layer-specific %signal change analyses and pRF modeling (cohort 2)

#### 7T structural data processing for layer-specific %signal change analyses

For cohort 2, 7T MP2RAGE structural data were collected in session 6. The data quality was evaluated first for each participant. Data showing severe artifacts (i.e., n=2 participants) were excluded. Only data showing mild truncation artifacts (not affecting SI) or no artifacts at all were used. All structural MP2RAGE images were sent to a pipeline including background noise cleaning, inhomogeneity correction, brain segmentation, cortical reconstruction and layer surface mapping. Background noise in all T1-weighted images was removed using the code of José P. Marques (https://github.com/JosePMarques/MP2RAGE-related-scripts) and the method described in O’Brien et al.^73^. Skull stripping was performed on T1-weighted images using Freesurfer (v7.3.0) mri_synthstrip routine. Based on the white matter (WM), gray matter (GM), and cerebrospinal fluid (CSF) segmentations, a brainmask was created and applied to the qT1 images for consistency. Inhomogeneity correction was done for both qT1 and T1-weighted images using the segment routine as implemented in SPM12 (Statistical Parametric Mapping, Wellcome Department of Imaging Neuroscience, University College London, London, UK) in MATLAB (R2018b, The MathWorks Inc., Natick, MA, 2018). Finally, resulting images cortex segmentation and cortical depth sampling was performed following the same steps as described for cohort 1 (see section ‘Extracting microstructural profiles and cortical layer compartments’).

For layer-specific analyses, functional time series (i.e., resting state data, blocked-design data, phase-encoded data) were registered to the MP2RAGE image using ANTs (v2.1.0, nearest neighbor interpolation). The resulting registration matrices were applied to the corresponding functional parameter maps (i.e., GLM estimates for localiser images and fourier transform estimates).

#### 3T structural data processing for cross-layer %signal change analyses and pRF modeling

Csurf (http://www.cogsci.ucsd.edu/~sereno/.tmp/dist/csurf) recon-all (implanted from Freesurfer (v7.3.0) (http://surfer.nmr.mgh.harvard.edu/) was used for brain segmentation and cortical surface reconstruction using the T1-weighted 3D MPRAGE image. As a fully automated processing pipeline, recon-all performs steps including intensity correction, transformation to Talairach space, normalization, skull-stripping, subcortical and white matter segmentation, surface tessellation, surface refinement, surface inflation, sulcus-based nonlinear morphing to a cross-subject spherical coordinate system, and cortical parcellation^74^. Skull stripping, segmentation and surface inflation quality were checked for each participant.

For cross-layer functional analyses, functional volumes were registered to the T1-weighted 3D MPRAGE image (same image used for recon-all brain segmentation) using csurf tkregister (12 degrees of freedom, non-rigid registration). And the x, y and z location of each surface vertex was mapped into functional voxel coordinates with the obtained registration matrix. The floating point coordinates of points at varying distances along the surface normal to a vertex were used to perform nearest neighbor sampling of the functional volume voxels (i.e., the 3D functional data were associated with each vertex on the surface by finding which voxel that point lay within).

### Functional tasks used for layer-specific analyses

For cohort 2, motion distortion correction was performed using the same method as described for cohort 1. Data quality was evaluated after acquisition for each participant, and data showing severe artifacts (i.e., n=4 participants) were excluded. Slice timing correction was performed on the remaining functional data to correct differences between slices during image acquisition using SPM12 (Statistical Parametric Mapping, Wellcome Department of Imaging Neuroscience, University College London, London, UK). Spatial smoothing was performed only to the blocked-design data with 1mm kernel width using SPM12.

### Blocked design analysis to locate index and middle finger in area 3b for cross-layer and layer-specific %signal change analyses and pRF modeling

For cohort 2, area 3b and the hand area were defined for each individual based on the atlas provided in csurf^75^. This approach was taken because finger-specific functional localizers were used here to detect index and middle finger representations. The functional representations of each finger were localized based on the pre-processed blocked-design data collected in session 6 (see section *Analysis of blocked design data*). We calculated 1st level fixed-effects models for the blocked-design runs using the general linear model (GLM) on individual data as implemented in SPM12. Each finger was treated individually and independently, therefore BOLD activation elicited by each finger’s tactile stimulation was treated as an independent measure^3^. Two regressors of interest were modeled explicitly (stimulation to index finger and stimulation to middle finger) and the rest condition was modeled implicitly. The linear contrast estimate for index and middle finger was computed: F contrast [1 0 / 0 1], replicated for each session. On the individual subject level, voxels that survived a significance threshold of *p* < .05 (uncorrected) were mapped onto individual cortical surfaces generated by csurf. The overlapping area between the hand area^75^ and the index and middle finger representation in area 3b of each individual was used as ROI for %signal change analyses and pRF modeling.

### Cross-layer and layer-specific %signal change analyses for sensory input signals in layer IV and altered functional response profile

Phase-encoded data collected in session 7 were averaged time point by time point by reversing the direction of time on a scan-by-scan basis, as the time series of different cycle directions (forward or reversed) were mirror-symmetric (see above) to each other. The time-reversed cycle direction (down→top cycle) was time-shifted before averaging by 4 seconds (2 TRs) to compensate for hemodynamic delay. Averaging was performed in 3D without any additional registration. Neither normalization nor smoothing was performed on the data during this procedure.

For cohort 2, we followed the same process as section ‘*%signal change analysis for structure-function correlation analyses*’, performing the %signal change analysis on the phase-encoding runs (only scans with stimulation cycles were used) for the cross-layer %signal change. There were 8 stimulation cycles for each run, serving as input frequencies. In addition to the above outlined procedure, the complex-valued phasing-mapping data (real and imaginary maps) were registered to the structural MP2RAGE images and segmented into three layer compartments using MIPAV. %signal changes were calculated at each layer compartment for each participant and for each condition.

### Population receptive field modeling for sensory input signals in layer IV and altered functional response profile

For cohort 2, pRF mapping of the index and middle fingers was performed on phase-encoded data collected in session 7 using a Matlab-based toolbox SamSrf v9.4 (freely available for download from https://github.com/samsrf/samsrf). The toolbox provides a generic framework to model pRFs with stimulus space in any dimension^76^. The phase-encoding dataset was used here for pRF mapping, with an additional rest condition to provide baseline information. The forward-order runs and reversed-order runs were first converted into Freesurfer MGH format and projected onto the Recon-all reconstructed surface. Superficial points along the surface normal to each vertex (upper 20% of the cortical thickness) were disregarded to minimize the effect of superficial veins on BOLD signal change. Only the mean value of 20-100% cortical depth was used. After converting from MGH to SamSrf format, the data were concatenated and averaged to increase the signal-to-noise ratio. The data were z-scored and linearly detrended. A stimulus aperture was created to define the somatosensory space. To represent the finger areas that were stimulated during fMRI scanning, the aperture was set up with the dimension of 100×100×320 (320=number of TR). Four finger areas were constructed (two sections in distal phalanx and two sections in intermediate phalanx), with each one associated with each of the stimulation units, based on the stimulation timing. Modeling ROI was defined manually by choosing vertices in the inflated surface model with restricting y-coordinates from −55 to 5 and z-coordinates from 30 to 80 in Freesurfer space, which covered the full SI area. Since the stimulation was applied along one dimension of the fingers, a 2D Gaussian tuning curve vertical model was used to perform the model fitting only on the y-dimension (within-digit dimension), with the x-dimension set as a constant. The receptive field profile determines the predicted response for a given location. The predicted time series is convolved with a canonical haemodynamic response function (HRF).

The parameter modeling employed a two-stage coarse-to-fine procedure to obtain the best possible fit of the predicted time series with the observed data. First, a coarse fit process was applied, which involved an extensive grid search by correlating the observed time series against a set of predicted time series derived from a combination of y_0_ and σ, covering the plausible range for each parameter. The search space was set at [-1.25 1.25], which covers more than the stimulus space [-1 1]. The parameters giving rise to the maximal correlation can survive and be sent to the second stage. The fine fit is an optimized procedure to refine the parameters by further maximizing the correlation between observed and predicted time series. The noise ceiling was calculated as an estimate of the maximum goodness-of-fit that could theoretically be achieved from the data of each voxel^76^, using a similar procedure as Urale et al.^77^. The Spearman-Brown prophecy formula^78^ was used to account for the accurate estimate of the true reliability of the time series. Six other parameters were estimated during modeling, including the center location (μ), the sizes (σ), goodness-of-fit (R^2^), normalized goodness-to-fit (nR^2^), baseline and beta (β). The R^2^ is the coefficient of determination of the correlation between the observed and predicted time series, and nR^2^ is the normalized coefficient of determination relative to the noise ceiling. During modeling, baseline (intercept) and β parameters (i.e., BOLD response amplitudes) were estimated for each voxel. After extracting the modeled parameters within the ROI, the resulting data were cleaned by preserving only those vertices with positive β values. The average pRF size (σ) of each condition and of each participant was calculated based on the cleaned data.

### Statistical analyses of Human MRI data

For cohort 1, all statistical analyses were carried out in R (version 4.2.2, R Core Team, 2022). All sample distributions were analyzed for outliers using boxplot methods and tested for normality using Shapiro-Wilk’s test. Homogeneity of variances was tested with Levene’s test. We applied two-sided tests. The significance level was set to p=0.05. Bonferroni-corrected significance levels were applied for multiple testing to correct for familywise error accumulation. To account for skewed and heteroscedastic data, we calculated robust permutation tests^79^. Group differences between younger and older adults (i.e., cortical thickness and septa analyses) were investigated with independent-samples random permutation Welch t-tests (100000 Monte-Carlo permutations, two-sided, equal-tail permutation *p*-value) using the MKinfer package (version 1.1, https://cran.r-project.org/package=MKinfer). Permutation *p*-values (*p_perm_*) are set to a minimum value of 10^-5^, limited by the number of permutations (i.e., minimum value of *p_perm_*=1/number of permutations). To investigate interaction effects between the factors age (between-subjects factor, levels: younger, older) and layer (within-subjects factor, levels: outer, middle, inner) as well as between the factors, age, layer and body-part (with-subjects factor, levels: thumb, index, middle, ring, little finger, or levels: hand, face) we calculated mixed-effects permutation ANOVAs (type III) using the “Rd_kheradPajouh_renaud” method^80^ for random effects models as implemented in the permuco package (version 1.1.2) written for R (version 4.2.2, R Core Team, 2022). The number of permutations was set to 100000. For comparability reasons we report both, results of parametric mixed-effects ANOVAs (using Greenhouse-Geisser corrected degrees of freedom and p-values in case of sphericity violations) and non-parametric permutation mixed-effects ANOVAs. Generalized eta squared (*η_G_²*) was calculated as an effect size estimator for parametric ANOVAs. To follow-up significant interaction effects, we calculated Welch t-tests using bootstrapped samples to estimate confidence intervals and p-values (as implemented in the MKinfer package version 1.1)^81^, accounting for non-normality in underlying conditions. For comparability we report both results, i.e., with and without bootstrapping. To regress out the effect of map size on layer-specific structural values, we calculated group-wise linear regression models using the lme 4 package (version 1.1.33) while controlling for the effect of individuals (random intercept model, e.g. middle qT1 ∼ map size + 1/participant). Intercepts were re-added to the resulting residuals before they were taken forward for between-finger and hand-face comparisons. Bayesian independent-sample t-tests were used to compare cortical thickness between younger and older adults using two-tailed tests, the Bayes Factors are specified as BF_10_ (detailed explanation on Bayesian analyses see below).

For cohort 2, the JASP software package (Version 0.17.1, JASP Team, 2023) was used to calculate Bayesian independent-sample t-tests for each stimulation condition on % signal change and pRF size (σ) to compare younger with older adults. Shapiro-Wilk tests were used to test on data normality. Student t-tests were used when data were normally distributed, and Mann-Whitney tests with 5 chains of 1000 iterations were used when data were not normally distributed. The choice of priors (Cauchy, scale=0.707) and the Markov chain Monte Carlo settings were the default as implemented in JASP (for all Bayesian tests)^82^. For pRF size and activation comparisons between younger and older adults, one-tailed tests were used to test for the alternative hypothesis σ_younger_ < σ_older_, as previous studies have found both enlarged pRF sizes and greater activation in older adults compared with younger adults^2,3^. As reduced coactivated inhibition has been evidenced with aging^2^, the alternative hypothesis was set as σ_older_ < σ_younger_ for pRF size difference between the double finger condition and the average of single finger conditions, and %_older_ < %_younger_ for % signal changes difference between double finger condition and the sum of single finger conditions. The Bayesian Factors are specified as BF_+0_ for one-tailed tests. Following Lee and Wagenmakers (2014)^83^, we interpret the Bayes Factor in support of H1 (BF_10_) in the following way: BF_10_=1-3 anecdotal evidence, BF_10_=3-10 moderate evidence, BF_10_=10-30 strong evidence, BF_10_=30-100 very strong evidence, BF_10_ > 100 extreme evidence, and in support of H0: BF_10_=0.33-1 anecdotal evidence, BF_10_=0.1-0.33 moderate evidence, BF_10_=0.03-0.1 strong evidence, BF_10_=0.01-0.03 very strong evidence, BF_10_ < 0.01 extreme evidence.

### Human behavioral measurements

#### Cohort 1

##### Tactile Detection Task

Tactile detection of touch to the surface of the fingertip was assessed using fine hair stimuli (sessions 4 and 5). We used a subset (0.008 g, 0.02 g, 0.04 g, 0.07 g, 0.16 g, 0.4 g, 0.6 g, 1.0 g, 1.4 g, 2.0 g, 4.0 g, 6.0 g) of standardized tactile monofilaments (Semmes-Weinstein monofilaments; Baseline, Fabrication Enterprises Inc., White Plains, NY, USA) to apply different mechanical forces to the skin surface of the fingertips (see Fig. 1). Stimuli were manually applied to a predefined skin area (circle with a diameter of approximately 2 mm), touching the skin surface at an angle of approximately 90 degree, for one second^84^. Stimulus application was guided by auditory instructions via headphones which were controlled by the Psychophysics Toolbox extension in MATLAB (R2017b). All participants sat in front of a screen that signaled the beginning and ending of stimulus intervals and listened to white noise via headphones. The right hand (palm facing upwards) was fixated on a small pillow behind a paper wall, preventing participants from seeing their own hand and the experimenter.

In a two-alternative forced choice paradigm, participants chose one of two possible time intervals that contained the stimulation (randomly applied in the first or second interval)^85^. Application of tactile monofilaments followed a 3-down/1-up staircase approach with two interleaved staircases, one starting at a weight of 0.02 g and the other starting at a weight of 0.4 g. The stimulus weight was increased by one step after each error, and decreased by one step after every three correct responses, until stable performance was reached. This procedure was separately applied for each of the two staircases^86^. The experiment was finished when for the last 30 trials the standard variation in stimulus intensity was 1 step or less^85^, or when the maximum number of 100 trials was reached. The participant‘s tactile detection threshold was defined as the mean stimulus intensity across reversal points (change of response from correct to incorrect or incorrect to correct) within the period of stable performance (i.e., the last 10 trials). The experiment took approximately 12 min per finger.

Before averaging, stimulus intensities were transformed logarithmically on a 1/10th milligram scale (log 10 0.1mg), yielding approximately equal intervals between filaments. Lower values indicate higher tactile sensitivity to mechanical forces. Additionally, we estimated the skin indentation in mm based on the examined detection threshold. Detection thresholds were taken as proxies of corresponding values in milliNewton (mN) that were provided by the manufacturer. Afterwards, the skin indentation δ was calculated according to the following equation^87^:

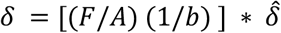

F is the estimated force in Newton (N), A (=0.2368 N) and b (=2.0696) are material/structural constants and δ^^^(=1.00 mm) is the reference indentation^87^. Finally, the result (δ) was multiplied by 3 to get an indentation value clearly above threshold, which was used to increase the amplitude of pin movement in the 2PD task. All calculations were performed in Matlab (R2017b).

##### Finger Discrimination Task

Again, Semmes Weinstein monofilaments were used to apply tactile stimulation, targeting the same stimulation sites as described for the tactile detection task. In a five-alternative-forced-choice design, tactile stimulation (lasting 1 second) was applied to one of five possible fingertips (session 5). Participants were asked to name the finger where they felt the touch. Answers were given verbally within a limited response interval (lasting 7 seconds). In case participants perceived no touch (note that tactile stimulation was applied at individual detection thresholds and was therefore expected to be perceived only in approximately 50% of the cases), they were motivated to guess. Each fingertip was stimulated 20 times, using unique pseudo-randomized sequences (with fingertips being stimulated not more than two times in a row). To extract the finger discrimination sensitivity (sensitivity of one finger being correctly discriminated from other fingers), we applied signal detection theory and calculated the d-prime as bias-free index of discrimination sensitivity^88^ by computing the amount of times a specific finger was touched and detected (hit), or was not touched but falsely detected (false alarm). Hits and false alarms were first converted to z-scores before subtracting false alarms from hits. D-primes were obtained for each finger separately.

##### 2-Point Discrimination Task (2PD Task)

We assessed the tactile discrimination performance of the right index finger (session 4)^89^. Stimulation was applied by two rounded pins (diameter=0.4 mm) simultaneously touching the surface of the fingertip. A custom-made, fully automatic stimulation device moved the pins up and down, controlled by the software package Presentation (version 16.5, Neurobehavioral Systems, Inc., Albany, CA, USA). The amplitude of pin movement was adjusted to the individual detection threshold (as assessed before in the tactile detection task), but was at least set to 1.2 mm. Spacing between pins ranged from 0.7 to 2.8 mm (in steps of 0.3 mm) for younger adults and from 0.7 to 6.3 mm (in steps of 0.8 mm) for older adults. Additionally, a single pin was included as control condition. Pin spacing was vertically adjusted by rotating a disc containing all possible pin spacing conditions (n=9). In a two-alternative forced-choice paradigm, pin spacing conditions were pseudo-randomly presented. Participants indicated whether they perceived one or two single pins touching their fingertip. They were instructed to give the answer ‘two pins felt’ only if they were certain. The right index finger was fixated on the stimulator, and the hand was covered by a white box during the task to prevent effects caused by seeing the stimulated finger^90^. Each task block included 90 trials (10 repetitions per pin condition). To prevent order effects, unique sequences of pin spacing conditions were used per participant and run. All participants completed two runs. Intertrial intervals were pseudo-randomized and varied between 1 to 5 seconds. 2PD thresholds were calculated per participant and run. Answers “two pins felt” were fitted as percentages across ascending pin distances (e.g. 0.7-2.8 mm). A binary logistic regression was used to fit the data using the glmfit function (iterative weighted least square algorithm) from the Statistics Toolbox as implemented in MATLAB R2017b. The 2PD threshold was taken from the pin distance where the 50 percent level crossed the fitted sigmoid curve^89^. Lower values indicate higher spatial acuity.

Z-transformed false alarm and hit rates were calculated for each participant to derive d-prime values as bias-free indices of 2PD sensitivity. Hit rates were calculated as the proportion “2 pins felt” responses when the stimulus consisted of two pins. False alarm rates were calculated as the proportion “2 pins felt” responses when the stimulus consisted only of one pin. False alarm rates were adjusted to 0.1 by default, if no false alarm was detected^89^.

##### Precision Grip Task

Sensorimotor integration performance was assessed with a custom made pressure sensor that was held between the thumb and index finger of the right hand, adjusted to individual strength (session 4, see Fig. 1)^91^. Reference forces that were to be matched ranged from 5% to 25% of the individual maximum grip force to avoid muscle fatigue^91^. Participants solved a visuo-motor matching task^91^, demanding them to continuously adjust the grip force. Applied forces were sampled at a frequency of 100 Hz and projected on screen at a refresh rate of 60 Hz. The task was controlled by the software package Presentation (version 16.5,

Neurobehavioral Systems, Inc., Albany, CA, USA). Each task repetition contained a unique pseudo-randomized sequence of 10 position changes at five different amplitudes (5%, 10%, 15%, 20%, 25% of maximum grip force), leading to a mean frequency of 0.25 Hz. After a period of task familiarization^91^, all participants performed the task for a total duration of 20 seconds. One run contained 15 trials divided by intertrial intervals of 10 seconds, leading to a total duration of about 8 minutes per run. All participants performed two runs that were separated by a 5-minute resting period. After each trial, participants received feedback about their individual performance level on screen. We monitored the time (in seconds) the controllable bar was within a given percentage above (2.5%) and below (2.5%) the target line (upper edge of the reference bar)^91^. Higher values reflect better sensorimotor integration.

### Cohort 2

#### Tactile detection task

We measured the tactile threshold of the index and middle finger of the right hand. During testing, participants were asked to sit on a chair with the right hand positioned on a custom-build, metal-free applicator with an independently-controlled MR-compatible piezoelectric stimulator (same device used during fMRI task) set underneath the tested finger. The tested hand was occluded from view. The tactile threshold was detected with a two-alternative forced task. At each trial, two intervals were presented with only one of them containing a stimulation, which was one pin rising up during a stimulation with a certain amplitude, lasting for 1 second. Participants were asked to detect the stimulation interval by pressing the respective key on the keyboard in a self-paced manner (“1” or “2”). A randomized sequence (different for each participant) was used to determine which interval contained the stimulation. The adaptive thresholding procedure followed a 3-down/1-up staircase algorithm. For each finger, the stimulation amplitude started at 0.73 mm. Every time the participant chose the correct interval three times in a row, the amplitude went down for 0.03 mm, whereas if the participant chose the wrong interval, the amplitude went up for 0.03 mm. There were 30 trials to start with, and the participants were expected to reach an accuracy above 80%. If the accuracy was below 80%, there would be more trials until the accuracy reached the requirement. The task took approximately 45 to 60 minutes.

#### Texture roughness discrimination

The texture roughness discrimination test was used to detect individual tactile sensation of surface roughness. The test comprises a plastic board with 15 columns, and the size of each column is 1.0×1.5 cm. Each column consists of small pins arranged with densities at different levels, resulting in a roughness ranking from rough (level 1) to smooth (level 10). During testing, participants were asked to sit on a chair with the arm positioned on a foam cushion. The tested hand was occluded from view. At each trial, two intervals were presented containing stimulation with two different roughness stimuli. Participants were asked to detect the column with higher roughness (i.e., less density). They were asked to press the respective key on the keyboard in a self-paced manner (“1” or “2”). For stimulus application, the experimenter followed auditory instructions via headphones. Neither the hand nor the experimenter was visible to the participant during testing. The plastic board was positioned under the participants’ right hand during the stimulation and withdrawn afterwards. The test was composed of two conditions, each included 30 trials. During the first testing condition, the tactile sensation of the index finger was tested and during the second testing condition, the tactile sensation of index and middle finger combined was tested. The stimulation pairs were chosen under three different conditions, with the roughness level increasing consecutively (e.g., level 3 and level 4) or in increments of two (e.g., level 3 and level 5) or three (e.g., level 3 and level 6). Stimulation pairs too easy or too difficult to distinguish were excluded. The stimulation order for each pair and for the sequence were randomized for each participant. The test took approximately 45 to 60 minutes.

#### Hand dexterity

Two standard tests (i.e., the Grooved Pegboard Test and The O’Conner Finger Dexterity Test) were used to test individual levels of hand motor function^3^, each consisting of two conditions. During one testing condition, participants were asked to only use their thumb and index finger of the right hand, whereas during the other testing condition, they were asked to use thumb, index finger and middle finger.

We also performed the Small Motor Test. It was employed to explore the haptic object sensation and coordinated motor movement function, which also consisted of two conditions. During one testing condition, participants were asked to only use their thumb and index finger of the right hand to pick up the pins, whereas during the other testing condition, they can use thumb, index finger and middle finger. It was composed of hexagonal-shaped nuts and small metal pins. The task was to pick up a pin using two fingers or three fingers to fill in the hole on a hexagonal-shaped nut held by the left hand, after success then move on to the next pair. As there is no requirement on how they could hold the nut with the left hand, it requires coordinating fingers of the right hand for holding the pin, and coordinating both left and right hand to aim for the hole. We measured the number of pins and nuts that were successfully paired with each other (n).

### Statistical analyses of human behavioral data

For cohort 1, all behavioral measures (tactile detection thresholds, 2PD thresholds and 2PD sensitivity, finger discrimination sensitivity, precision grip accuracy) and functional outcome variables (%signal change, network centrality) were compared between age groups using independent-samples random permutation Welch t-tests (100000 Monte-Carlo permutations, two-sided, equal-tail permutation *p*-value) as implemented in the MKinfer package (version 1.1) for R (version 4.2.2, R Core Team, 2022).

Tactile detection thresholds and 2PD sensitivity of the right index finger, precision grip accuracy (i.e. sensorimotor integration) involving both the right thumb and the right index finger as well as %signal change (i.e. responsivity) and network centrality (i.e. EC) of the contralateral index finger representation were correlated with layer-specific qT1 values of the contralateral index finger representation. Correlation analyses were performed using Kendall’s tau correlation coefficient. Uncorrected results are reported. Before calculating correlations, the data was partly transformed (using the reciprocal of tactile detection thresholds and qT1 values), so that in the final scatterplots higher values always indicate better performance in behavior, higher responsivity, and more connectivity in fMRI markers, as well as higher substance concentration in structural MRI markers. Regression lines in the scatterplots (see Fig. 5) were generated based on robust linear Theil-Sen regression estimates^22^, because the estimation of the regression coefficients is based on Kendall’s tau^23^. Cut-offs for the interpretation of Kendall’s tau correlation coefficients were approximated with the following equation, where *r* denotes the Pearson correlation coefficient^24^:

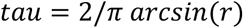

To derive the cut-off values, we applied the equation to a previously stratified convention for the Pearson correlation coefficient^25^.

For cohort 2, behavioral statistics included data of 21 healthy younger adults and 17 healthy older adults. Nonparametric independent sample Mann-Whitney U tests were used to compare texture roughness and hand dexterity measures between age groups. Effect sizes are given by the rank biserial correlation.

The functional signal central peak at layer IV of the index finger and double-finger (index and middle finger together) representation were correlated with the texture roughness discrimination accuracy, the number of holes filled in the Grooved Pegboard Test and the number of pairs completed in the Small Motor Test for the index finger and the double finger condition, respectively. Correlation analyses were performed using Kendall’s tau correlation coefficient. Uncorrected results are reported.

### Animal experiments

Calcium imaging experiments were performed in younger adult (2-6 months; n=8) and older adult mice (12-20 months; n=8; ages chosen for equivalent ranges to the human cohorts 1 and 2, see also Wang et al.^92^), both male and female of the transgenic line expressing genetically encoded calcium indicators (GCaMP6f; C57BL/6J-Tg (Thy1-GCaMP6f) GP5.5Dkim/J; RRID: IMSR_JAX:024276). Mice were housed in standard cages at a 12 h/12 h light dark cycle, food and water were provided ad libitum. Histological analysis was performed in 12 of these mice in relation to the expression of PV positive (PV+) neurons and in an additional 26 mice in relation to the expression of PV+ neurons, MBP expression as an indication of myelination, and Iba1 expression as a marker for microglia (younger adult mice [n=11, 2-6 months], older adult mice [n=7, 12-20 months], and mice in old age [n=8, +24 months]). All experiments were performed according to the NIH Guide for the Care and Use of Laboratory animals (2011) (National Research Council (US) Committee for the Update of the Guide for theCare and Use of Laboratory Animals, 2011) and the Directive of the European Communities Parliament and Council on the protection of animals used for scientific purposes (2010/63/EU) and were approved by the animal care committee of Sachsen-Anhalt, Germany.

### Mice cranial window

For cranial window implantation, mice were anesthetized with isoflurane (4% for induction and 2% maintenance during surgery). Eye cream was applied to protect the eyes (Bepanthen, Bayer), analgesics and anti-inflammatory drugs were injected subcutaneously (carprofen, 0.15 mg, and dexamethasone, 2 mg). After shaving and disinfection of the scalp, a section of skin was removed. A craniotomy was then made over the left S1-barrel field, stereotaxically targeting the D1-D3 barrel fields (from bregma: AP −1.22 mm; ML 3 mm ^94^), the window was 3 mm in diameter. A glass coverslip was then placed on top of the craniotomy and fixed with glue (gel control, Loctite). A custom-built headplate was implanted on the exposed skull with glue and cemented with dental acrylic (Paladur, Heraeus Kulzer).

### Mice two-photon calcium imaging

Two-photon calcium imaging was performed using an 8 kHz resonant scanning microscope (HyperScope, Scientifica) with an Ultrafast laser (InSight X3 Dual output laser; Spectra-Physics; < 120 fs pulse width, 80 MHz repetition rate) tuned to 940 nm. Images were acquired at 30 Hz (using a 16X objective 0.8 NA, Nikon; tilted 10° from vertical for S1-barrel field imaging with ScanImage software [Vidrio Technologies]). Imaging of layer II/III S1-barrel field, stereotaxically targeted to the D1-D3 barrels and spanning a 500-700 µm field of view (FoV), was performed at cortical depths between 150-250 μm and layer V imaging between 420-550 µm, each FoV was imaged under 3 whisker stimulation conditions. The size of the imaging FoV (500-700 µm) potentially spanned ∼2 neighboring barrel fields including dividing septa within the stereotaxically targeted cranial window. Note that the imaging FoV was delineated based on the optimal imaging parameters to maximize the number of neurons but without precise spatial receptive field mapping of the individual barrel fields. For all imaging sessions, mice were awake, head restrained, and placed on an air-suspended polystyrene 20 cm ball (Jet Ball system, PhenoSys GmbH, sampling frequency 30 Hz). The sampling frequency of the calcium imaging was resampled to precisely match the sampling frequency of the optical encoders, sensory stimulation, 30Hz camera recordings (ImagingSouce, DMK22BUC03) from a side-view of the face region, and behavioral readouts. Whisker stimulation was performed by an implemented air puff system (PhenoSys GmbH). Animals were in the dark for all trials. Regions of interest from the calcium imaging and whisker pad estimations (see below) from the behavioral video tracking (DeepLabCut^95^) were matched to the behavioral data file. This was done by downsampling and interpolating, ensuring that the aligned datasets had the same total number of frames, with an overall sampling frequency of approximately 30 Hz.

### Mice whisker stimulation

All three stimulation conditions consisted of randomized airpuffs directed to the right side whiskers and two-photon imaging performed in the left barrel field. Two FoVs were imaged for each mouse under each whisker stimulus condition, one in the outer layers (layer II/III) and one in the inner layers (layer V). First, all whiskers were stimulated (all whisker condition); after trimming the whiskers except for only two whiskers remaining on the right whisker pad (two of D1-D3 whiskers) the second session was performed (double stimulation condition; W1+W2), and after trimming one of the remaining two whiskers the last session of stimulating was performed (single stimulation condition; W1). The stimulation protocol consisted of ∼30 airpuffs (200 ms duration) delivered at randomized inter-stimulus intervals (range 6-20 seconds) for each field-of-view flanked by 2 minutes imaging of spontaneous activity (10 min total per FoV). Whisker-pad estimations and frame-by-frame tracking were conducted on the video recordings captured during the session. The side view of the animal’s face was analyzed using DeepLabCut^95^. In brief, we trained a model for pose-estimation on the side-view videos to track the whisker pad. The pose-estimation tracking was subsequently used to identify frames in which the animal was moving its whisker pad, with whisker movement defined on a frame-by-frame basis as periods with an instantaneous speed ≥ 0.5 cm/s, 0.25 Hz lowpass filtered speed ≥ 0.5 cm/s, and an average speed ≥ 0.5 cm/s over a 1 s window centered at this point in time. Any inter-movement interval shorter than 500 ms was also labeled as movement.

### Mice histology

Animals were deeply anesthetized with ketamine (20 mg/100 g body weight, ip) and xylazine (1 mg/100 g body weight, ip) and perfused transcardially with 20 mL of 0.1 M phosphate-buffered saline (PBS, pH 7.4) followed by 200 mL of 4% paraformaldehyde. The brains were extracted, post-fixed overnight in 4% paraformaldehyde at 4°C, and then cryoprotected in 30% sucrose in PBS for 48 h. Brains were cut into 50 μm thick coronal sections (CryoStar NX70, Thermo Scientific, USA) and collected in PBS (free-floating). For immunohistochemistry, serial series of floating sections were then blocked in normal donkey serum (10% and 0.4% triton in PBS) for 1 h and incubated in primary antibodies overnight at 4°C to visualize parvalbumin (PV, monoclonal mouse anti-parvalbumin, 1:4000, Swant, PV 235, RRID: AB_10000343; MBP, monoclonal mouse anti-myelin basic protein, 1:500, Santa Cruz, RRID:AB_10655672; Iba1, rabbit recombinant monoclonal anti-Iba1 antibody, 1:2000, Abcam, RRID: AB_2636859). After rinsing in PBS, sections were incubated for 2 h with donkey anti-mouse Cy3 (1:200, Jackson ImmunoResearch Labs Cat# 715-165-150, RRID: AB_2340813, United Kingdom) or for the Iba1 staining donkey anti-rabbit Cy3 (1:200, Jackson ImmunoResearch Labs, RRID: AB_233800). Finally, sections were rinsed again in PBS, mounted on gelatin-coated slides, and coverslipped with mounting media containing DAPI (4′,6-diamidino-2-phenylindole) counterstain (Vectashield). High resolution images were captured using an epifluorescence slide scanner microscope (Axioscan 7, Zeiss, Germany) under a 10x objective and merged (resulting in a ∼15000 × 11000 pixel image per coronal section).

### Mice data analysis

Two-photon calcium imaging datasets (30Hz) were motion corrected and cell detection and signal extraction performed using Suite2p^96^. To calculate the change in fluorescence (ΔF/F), the pixel intensity is averaged within each ROI, representing the change in intensity for a single neuron, to create a raw fluorescence time series F(t). Baseline fluorescence F0 was computed for each neuron by taking the 5th percentile of the smoothed F(t) (1 Hz lowpass, zero-phase, 60th-order FIR filter) and the change in fluorescence relative to baseline (ΔF/F0) was calculated (F(t)-F0/F0). We used nonnegative matrix factorization (NMF), in order to remove neuropil contamination as implemented in FISSA^97^. All further analyses were performed using custom-written scripts in MATLAB (MathWorks, MA, USA). Histological analysis was performed on 3 representative sections of a 1 mm medio-lateral extent for each brain across the entire cortical depth. Images were aligned to the cortical depth matching the surface, layer IV and the ventral white matter border as demarcated by DAPI. DAPI staining was also used to demarcate the cortical layers in the aligned images, namely outer layers, supragranular (I/II/III); middle layers, granular (IV); and inner layers, infragranular (V/VII) of the barrel cortex (using Fiji, ImageJ software)^98^. Cells were then automatically counted using the open-source software cellpose^99^ with the nuclei model (DAPI) and Cyto model (PV neurons or Iba1+ microglia) or the raw intensity summed across each row of pixels along the axis perpendicular to the cortical surface (MBP expression). For PV+ and Iba1+ counts, cell density was then calculated by normalizing to the area for the given demarcated cortical depths for each layer and across the given anterior posterior sections.

## Author contribution

Conceptualization: E.K., P.L., J.D., J.M.P.P.

Methodology: P.L., J.D., E.K., J.M.P.P., J.U.H., D.S.S., O.S.

Formal Analysis: P.L., J.D., J.M.P.P., J.U.H., A.N., L.C.L-C., E.B.

Investigation: E.K., P.L., J.D., J.M.P.P

Writing - original draft preparation: P.L., J.D., J.M.P.P., E.K., J.U.H.

Writing - review and editing: P.L., J.D., E.K., J.M.P.P., J.U.H., A.N., A.S., E.B., D.S.S., O.S

Visualization: P.L., J.D., J.M.P.P., J.U.H. Supervision: E.K., J.M.P.P.

Funding acquisition: E.K., J.M.P.P., O.S.

All authors contributed to the article and approved the submitted version.

## Data availability

Source data that underlie Figures for human studies are available here: 10.57754/FDAT.2davw-rgq96. Source data that underlie Figures for mice studies are available here: https://github.com/pakanlab/Liu_et_al_NatureNeuroscience2025. Any additional information required to reanalyze the data reported in this paper is available upon request. Due to data protection policies, raw MRI data from human studies are available upon request, requirements are a formal data sharing agreement and the need to submit a formal project outline.

## Competing interests

No competing interests to declare.

## Acknowledgments

This project was funded by the German Research Foundation (Deutsche Forschungsgemeinschaft, DFG) (KU 3711/2-1, project number: 423633679 to E.K., Project-ID 425899996 - SFB 1436/B06 to E.K and J.M.P.P., and project number 362321501 – RTG 2413 to J.M.P.P., and SFB 1158/B07, project number 255156212 to A.S.). P.L. and E.K. were supported by the European Research Council (ERC) under the European Union’s Horizon 2020 research and innovation programme (grant agreement No 949609).

We thank Prof. Dr. Herta Flor for collaborating with us on the congenital limb loss participant at the Central Institute of Mental Health (CIMH) Mannheim, Germany. We also thank Elnaz Khosroshahi, Melina Trigoussis, Lilith-Sophie Lange, Anastasia Chrysidou, Johanna Nolle and Sophie Christine Busalt for their support in participant recruitment and data collection, and Ridhim Joshi and Cathleen Knape for technical assistance and data collection in mice.

## Notes

### Competing Interest Statement

The authors have declared no competing interest.

### Summary of Updates

We updated the Bayesian statistics performed in this manuscript, and optimised with the interpretation.

