## Supplementary material for "Cortical sensory aging is layer-specific": Figure 1-supplemental table 1

### Supplemental Information

**Figure 1 - supplemental table 1. Overview of collected human 7T MRI data and its use for conducted analyses.**

| | Session number | Collected data | Purpose / Conducted analyses | Result sections |
|---|---|---|---|---|
| **Cohort 1** | Session 1 | qT1, QSM | cortical thickness, microstructural layer composition, low-myelin border analysis, structural topography analysis | 'Cortical thickness changes are layer-specific in older adults and with congenital arm loss', 'Altered microstructural layer composition in older adults and with congenital arm loss', 'Preserved low-myelin borders but greater structural changes in the face area compared to the hand area characterize the aging SI topography' |
| | Session 2 | Task-related blocked-design and phase-encoded fMRI data (tactile stimulation paradigm), Resting-state fMRI data | Localization of the fingers in SI: cortical thickness, microstructural layer composition, low-myelin border analysis, structural topography analysis | 'Cortical thickness changes are layer-specific in older adults and with congenital arm loss', 'Altered microstructural layer composition in older adults and with congenital arm loss', 'Preserved low-myelin borders but greater structural changes in the face area compared to the hand area characterize the aging SI topography' |
| | Session 3 | Task-related blocked-design fMRI data (motor movement paradigm) | Localization of the hand and face in SI: cortical thickness, microstructural composition analysis, low-myelin border analysis, structural topography analysis | 'Cortical thickness changes are layer-specific in older adults and with congenital arm loss', 'Altered microstructural layer composition in older adults and with congenital arm loss', 'Preserved low-myelin borders but greater structural changes in the face area compared to the hand area characterize the aging SI topography' |
| | Session 4 | Tactile Detection task (all fingers),<br>Finger Discrimination task (all fingers) | adjustment of stimulation intensity in session 4, age group comparison of behavior | 'Relation between layer-specific changes and human sensorimotor impairments' |

| | | | | |
|---|---|---|---|---|
| | Session 5 | Tactile 2-point discrimination task (index finger), Tactile detection task (index finger), Precision grip task (thumb+index finger) | Structure-behavior correlation, age group comparison of behavior | 'Relation between layer-specific changes and human sensorimotor impairments' |
| **Cohort 2** | Session 6 | qT1, Task-related blocked-design fMRI data and resting-state data | Microstructural layer composition, localization of index and middle finger in SI, BOLD signal calibration | 'More pronounced sensory input signals in layer IV in older adults', 'Altered functional response profile in older adults' SI', 'Relation between layer-specific changes and human sensorimotor impairments' |
| | Session 7 | Task-related phase-encoded fMRI data | Topographic maps, cross-layer and layer-specific % signal change and pRF modeling | 'More pronounced sensory input signals in layer IV in older adults', 'Altered functional response profile in older adults' SI' |
| | Session 8 | 3T structural image | Surface reconstruction for topographic maps and pRF modeling | 'More pronounced sensory input signals in layer IV in older adults', 'Altered functional response profile in older adults' SI' |
| | Session 9 | Tactile Detection Task (index finger and middle finger), O'Conner Hand Dexterity Test (thumb+index finger and thumb+index+middle finger), Grooved Pegboard Test (thumb+index finger and thumb+index+middle finger), Small Motor Test (thumb+index finger and thumb+index+middle finger), and Texture Discrimination Test (index finger and index+middle finger) | Structure-behavior correlation, age group comparison of behavior | 'Relation between layer-specific changes and human sensorimotor impairments' |
