## Supplementary material for "Cortical sensory aging is layer-specific": Table 1-supplemental table 1

**Table 1 - supplemental table 1. Cortical thickness comparison for the SI (i.e. BA 3b) hand and face areas localized by motor movements in relation to anatomically-relevant layer compartments.** Given are layer-specific (outer, middle, inner) mean cortical thickness values (Mean) and standard deviations (SD) in millimeters for hand and face regions of younger and older adults (localized by movements of corresponding body parts). Independent-samples random permutation Welch t-tests were calculated to investigate group differences (t=test statistic, df=degrees of freedom, $p_{perm}$=Monte-Carlo permutation p-value, $CI_{perm}$=95% Monte-Carlo permutation confidence interval, number of permutations=100000, minimum value of $p_{perm}$=1/number of permutations). Significant differences with Bonferroni-corrected $p < 0.006$ (correcting for 8 tests) are marked by *.

| | All<br>n = 34 | **Younger**<br>n = 16 | **Older**<br>n = 18 | **Group Differences** | | | |
|---|---|---|---|---|---|---|---|
| | Mean ± SD | Mean ± SD | Mean ± SD | $t$ | *df* | $p_{perm}$ | $CI_{perm}$ |
| **hand total** | 2.01 ± 0.10 | 2.06 ± 0.05 | 1.96 ± 0.11 | 3.24 | 25.3 | 0.003* | 0.03, 0.16 |
| hand outer | 0.47 ± 0.06 | 0.42 ± 0.02 | 0.52 ± 0.04 | -10.1 | 22.1 | $<10^{-5}$* | -0.14, -0.06 |
| hand middle | 0.67 ± 0.10 | 0.56 ± 0.01 | 0.75 ± 0.04 | -18.6 | 20.5 | $<10^{-5}$ * | -0.26, -0.12 |
| hand inner | 0.87 ± 0.20 | 1.08 ± 0.05 | 0.69 ± 0.05 | 24.4 | 31.4 | $<10^{-5}$ * | 0.25, 0.52 |
| **face total** | 2.13 ± 0.16 | 2.22 ± 0.10 | 2.05 ± 0.16 | 3.74 | 29.1 | $6.0^{-4}$ * | 0.06, 0.28 |
| face outer | 0.51 ± 0.06 | 0.46 ± 0.04 | 0.56 ± 0.05 | -6.95 | 31.1 | $<10^{-5}$ * | -0.14, 0.05 |
| face middle | 0.70 ± 0.11 | 0.60 ± 0.03 | 0.78 ± 0.08 | -9.46 | 23.0 | $<10^{-5}$ * | -0.25, -0.11 |
| face inner | 0.92 ± 0.23 | 1.16 ± 0.05 | 0.71 ± 0.07 | 22.01 | 31.8 | $<10^{-5}$ * | 0.29, 0.60 |
