## Supplementary material for "Cortical sensory aging is layer-specific": Table 1-supplemental table 2

**Table 1 - supplemental table 2. Thickness comparison for the SI (i.e. BA 3b) hand area (localized by pRF center location maps) using equal layer compartments (layer definition of younger adults)**. Given are layer-specific (outer, middle, inner) mean cortical thickness values (Mean) and standard deviations (SD) in millimeters for the BA 3b hand region (localized by vibro-tactile stimulation to the five fingertips of the right hand). Independent-samples random permutation Welch t-tests were calculated to investigate group differences (t=test statistic, df=degrees of freedom, $p_{perm}$=Monte-Carlo permutation p-value, $CI_{perm}$=95% Monte-Carlo permutation confidence interval, number of permutations=100000, minimum value of $p_{perm}$=1/number of permutations). Significant differences with Bonferroni-corrected $p < 0.016$ (correcting for 3 tests) are marked by *. Trends above Bonferroni-corrected threshold are marked by a T.

| | All<br>n = 40 | **Younger**<br>n = 20 | **Older**<br>n = 20 | **Group Differences** | | | |
|---|---|---|---|---|---|---|---|
| | Mean ± SD | Mean ± SD | Mean ± SD | $t$ | *df* | $p_{perm}$ | $CI_{perm}$ |
| hand outer | 0.41 ± 0.03 | 0.41 ± 0.02 | 0.40 ± 0.03 | 0.5 | 30.8 | 0.640 | -0.01, 0.02 |
| hand middle | 0.55 ± 0.03 | 0.56 ± 0.02 | 0.55 ± 0.04 | 0.7 | 28.0 | 0.538 | -0.01, 0.02 |
| hand inner | 1.05 ± 0.07 | 1.10 ± 0.06 | 1.00 ± 0.05 | 5.7 | 36.7 | $<10^{-5}$ * | 0.05, 0.14 |
