## Supplementary material for "Cortical sensory aging is layer-specific": Table 1-supplemental table 3

**Table 1 - supplemental table 3. Layer-specific cortical thickness of the SI hand area without outlier removal.** Shown are total and layer-specific (outer, middle, inner) mean cortical thickness values (Mean) and standard deviations (SD) in millimeters for the SI hand area. Independent-samples random permutation Welch t-tests were calculated to investigate group differences (t=test statistic, df=degrees of freedom, $p_{perm}$=Monte-Carlo permutation p-value, $CI_{perm}$=95% Monte-Carlo permutation confidence interval, number of permutations=100000, minimum value of $p_{perm}$=1/number of permutations). Significant differences with Bonferroni-corrected p < 0.0125 (correcting for 4 tests) are marked by *.

| hand area | All n = 40 | **Younger** n = 20 | **Older** n = 20 | **Group Differences** | | | |
|---|---|---|---|---|---|---|---|
| | Mean ± SD | Mean ± SD | Mean ± SD | *t* | *df* | $p_{perm}$ | $CI_{perm}$ |
| total | 2.01 ± 0.10 | 2.06 ± 0.07 | 1.95 ± 0.10 | 3.8 | 33.4 | $5.4 \times 10^{-4}$ * | 0.04, 0.17 |
| outer | 0.46 ± 0.06 | 0.41 ± 0.02 | 0.51 ± 0.04 | -10.1 | 26.8 | $< 10^{-5}$ * | -0.14, -0.07 |
| middle | 0.65 ± 0.10 | 0.56 ± 0.02 | 0.75 ± 0.04 | -18.7 | 26.0 | $< 10^{-5}$ * | -0.26, -0.13 |
| inner | 0.89 ± 0.21 | 1.10 ± 0.06 | 0.69 ± 0.04 | 26.6 | 31.0 | $< 10^{-5}$ * | 0.27, 0.54 |
