## Supplementary material for "Cortical sensory aging is layer-specific": Table 1-supplemental table 4

**Table 1 - supplemental table 4. Bayesian independent-sample t-tests on layer-specific cortical thickness differences of the SI hand area between younger and older adults.** Shown are total and layer-specific (outer, middle, inner) mean cortical thickness values (Mean) and standard deviations (SD) in millimeters for the SI hand area. Bayesian independent-sample t-tests were performed on younger (n=20) and older (n=19) adults. The alternative hypothesis $H_1$ is specified as $thickness_{young} \neq thickness_{old}$ for total, outer, middle and inner, and the null hypothesis $H_0$ is specified as no difference between younger and older adults on total cortical thickness and the thickness of each cortical layer compartment.

| | Age | | | | |
|---|---|---|---|---|---|
| | Younger adults (n=20) | Older adults (n=19) | | | |
| | Mean ± SD | Mean ± SD | $BF_{10}$ | error% | 95% Credible Interval |
| total | 2.06 ± 0.07 | 1.94 ± 0.08 | 1383.792 | $7.538 \times 10^{-9}$ | 0.41, 1.82 |
| outer | 0.41 ± 0.02 | 0.40 ± 0.02 | 0.635 | 0.005 | 0.04, 1.05 |
| middle | 0.56 ± 0.02 | 0.85 ± 0.03 | $8.931 \times 10^{21}$ | $5.925 \times 10^{-28}$ | -2.21, -0.68 |
| inner | 1.10 ± 0.06 | 0.69 ± 0.04 | $2.045 \times 10^{26}$ | $4.522 \times 10^{-24}$ | 0.72, 2.25 |
