## Supplementary material for "Cortical sensory aging is layer-specific": Table 1-supplemental table 5

**Table 1 - supplemental table 5. Microstructural layer composition of SI hand and face areas of a participant with congenital arm loss after controlling for map size.** Cortical fields (i.e. hand and face regions) were localized by motor movements or imagery of motor movements (for the missing limb condition). Given are layer-specific (outer, middle, inner) qT1 (in milliseconds), nQSM (in parts per million), and pQSM (in parts per million) values. Lower qT1 and nQSM values indicate higher substance concentration. For microstructure profiles plotted along the dimension of cortical depth see **Table 1 - supplemental table 3**.

| | qT1 | nQSM | pQSM | qT1 | nQSM | pQSM |
|---|---|---|---|---|---|---|
| | contralateral to missing limb | | | ipsilateral to missing limb | | |
| hand outer | 2107.4 | -0.017 | 0.015 | 2081.1 | -0.012 | 0.016 |
| hand middle | 1678.5 | -0.013 | 0.021 | 1705.2 | -0.007 | 0.017 |
| hand inner | 1499.5 | -0.011 | 0.022 | 1508.2 | -0.008 | 0.019 |
| face outer | 2142.4 | -0.017 | 0.021 | 2140.1 | -0.019 | 0.020 |
| face middle | 1794.3 | -0.019 | 0.017 | 1750.3 | -0.018 | 0.020 |
| face inner | 1559.9 | -0.014 | 0.015 | 1527.2 | -0.010 | 0.016 |
