## Supplementary material for "Cortical sensory aging is layer-specific": Table 1-supplemental figure 1

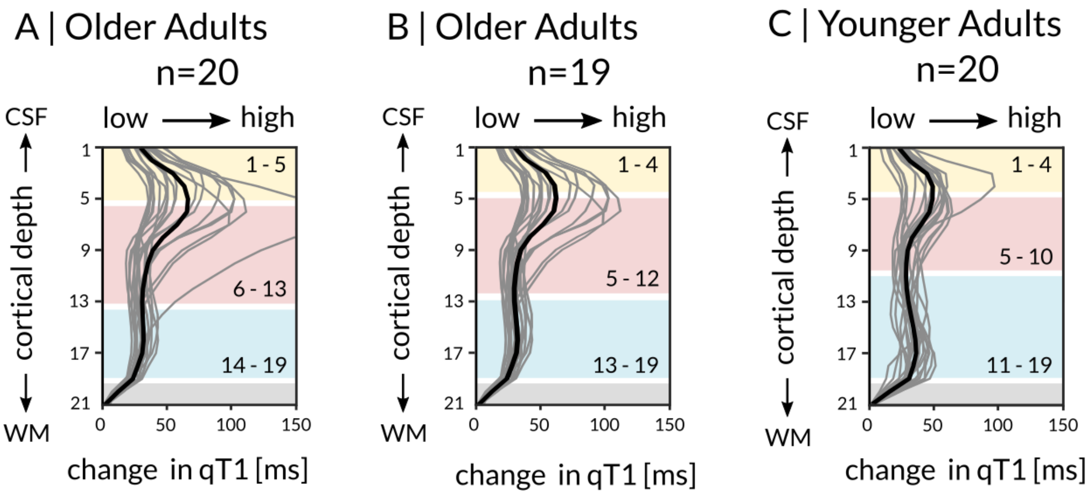


**Table 1 - supplemental figure 1. Anatomically-relevant cortical layer compartments. (A)** Cortical layer compartments (cream, outer layer; light pink, middle layer; light blue, inner layer) included in statistical analyses. The layer definition (i.e., minima and maxima of the first derivative of raw qT1; group mean plotted in black, individual data plotted in gray) was based on the left BA 3b hand area (identified by vibrotactile stimulation to the five fingertips of the right hand) using the full sample of older adults (i.e. n=20). **(B)** Cortical layer compartments after removing one outlier (participant 40) from the full sample, leaving n=19 older adults to define the cortical layer compartments. (C) Cortical layer compartments of younger adults shown as reference.
