## Supplementary material for "Cortical sensory aging is layer-specific": Table 1-supplemental figure 2

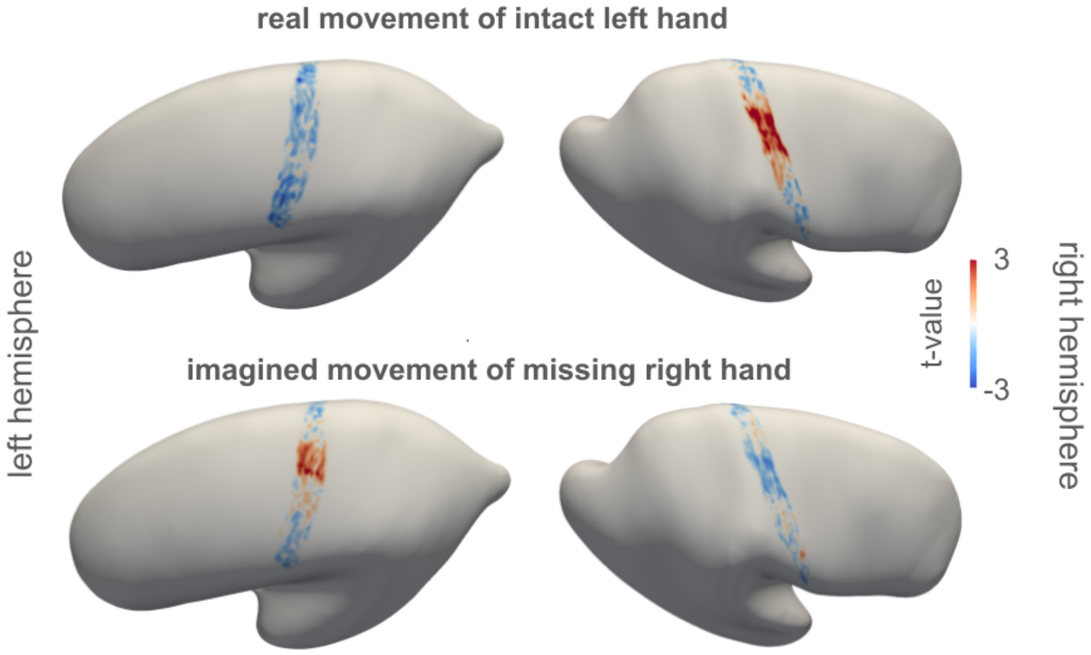


**Table 1 - supplemental figure 2. Functional localizers used to locate the hand area in area 3b of a healthy adult (male, age=52 years) with congenital arm loss on the right side.** Whereas real movement of the intact left hand induced strong activation in area 3b of the contralateral right hemisphere (colored in red), imagined movement of the missing right hand induced strong activation in area 3b of the contralateral left hemisphere (colored in red). Critically, the cluster of highest t-values (colored in red) overlaps an area where we, based on anatomical landmarks[1,2,3], know the hand area to be located.
