## Supplementary material for "Cortical sensory aging is layer-specific": Table 1-supplemental figure 3

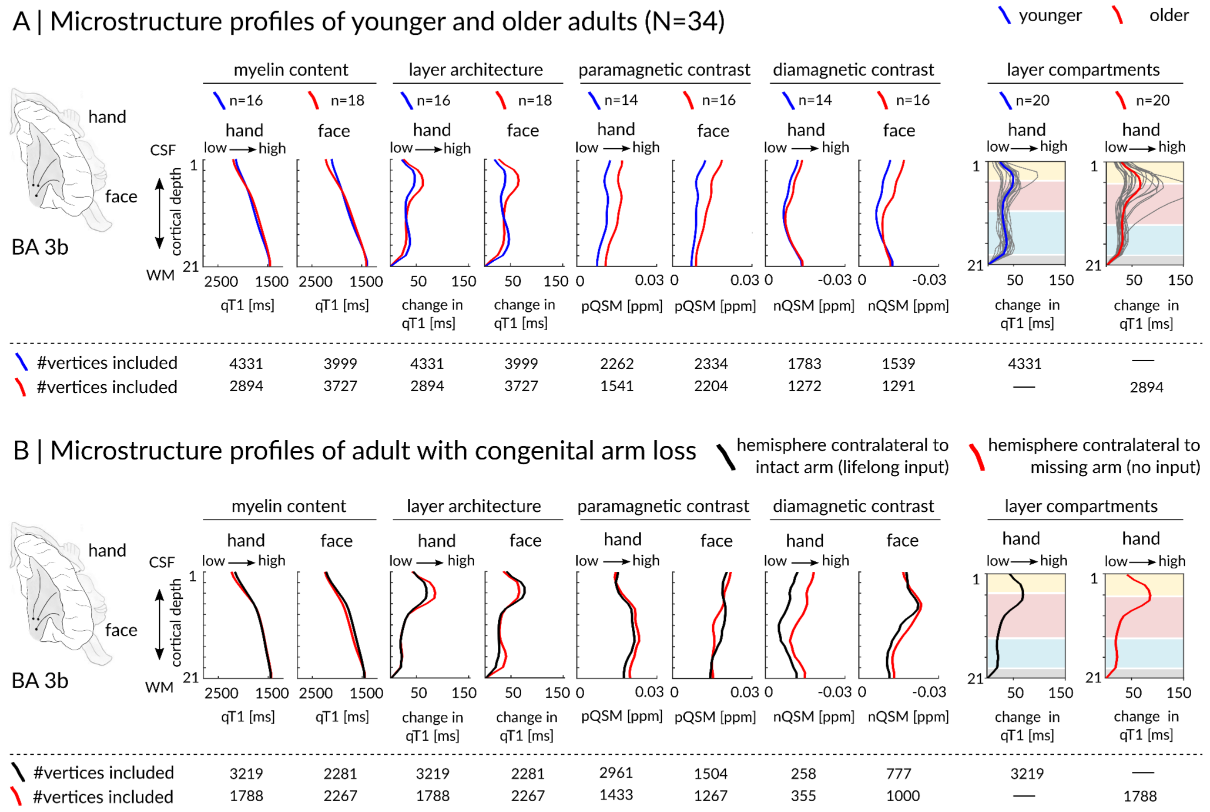


**Table 1 - supplemental figure 3. Structural SI Layer Architecture in Younger Adults, Older Adults and with Congenital Arm Loss. (A)** Microstructural profiles for younger adults (blue) and older adults (red), extracted from the left hemisphere (contralateral to finger and tongue movements). qT1 values are given in milliseconds (ms), nQSM and pQSM values in parts per million (ppm). Three layer compartments were extracted based on localizing maxima and minima of the first derivative of raw qT1 values (outer compartment: cream, middle compartment: light red, inner compartment: light blue). For qT1 and nQSM, lower values indicate higher substance concentration. **(B)** A healthy adult (male, age=52 years) with congenital arm loss on the right side. Microstructural profiles for the participant with congenital arm loss were extracted from the hemisphere contralateral (colored in red) and ipsilateral (colored in black) to the missing arm. The cortex of the hand area contralateral to the missing arm (identified via mental imagery of finger movements[6]) is thinnest (hand contralateral: 1.82 mm; hand ipsilateral: 1.88 mm; face contralateral: 1.96 mm; face ipsilateral: 1.93 mm). Layer-specific cortical thickness extraction reveals a thinner middle compartment (colored in light red) for the hand area contralateral compared to ipsilateral to the missing limb (hand contralateral: outer=0.47 mm, middle=0.72 mm, inner=0.63 mm; hand ipsilateral: outer=0.38 mm, middle=0.83 mm, inner=0.67 mm). Note that layer-specific thickness values of the one-hander are in a plausible data range (taking the range of two-handed participants for the hand area as reference: outer=0.38-0.58 mm, middle=0.53-0.83 mm, inner=0.60-1.17 mm).
