## Supplementary material for "Cortical sensory aging is layer-specific": Table 1-supplemental figure 4

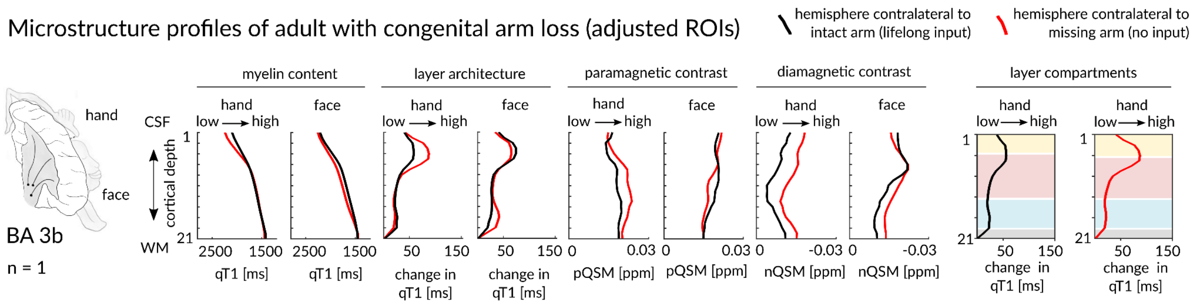


| | myelin content: hand | myelin content: face | layer architecture: hand | layer architecture: face | paramagnetic contrast: hand | paramagnetic contrast: face | diamagnetic contrast: hand | diamagnetic contrast: face | layer compartments: hand | layer compartments: hand |
|---|---|---|---|---|---|---|---|---|---|---|
| (black) #vertices included | 1652 | 1672 | 1652 | 1672 | 1558 | 1164 | 94 | 508 | 1652 | — |
| (red) #vertices inlcuded | 1661 | 1687 | 1661 | 1687 | 1337 | 938 | 324 | 749 | — | 1661 |

**Table 1 - supplemental figure 4. Microstructural profiles of the SI hand and face areas for the participant with congenital arm loss after controlling for map size**. Larger differences between ROI sizes contralateral (coloured in red) and ipsilateral (coloured in black) to the missing arm for pQSM and nQSM values are due to differences in the number of positive and negative vertices per hemisphere (because a vertex can either be positive or negative). qT1 values are given in milliseconds (ms), nQSM and pQSM values are given in part per million (ppm). Three anatomically-relevant layer compartments were extracted based on localizing maxima and minima of the first derivative of raw qT1 values (outer compartment: light yellow, middle compartment: light red, inner compartment: light blue). For qT1 and nQSM, lower values indicate higher substance concentration.
