## Supplementary material for "Cortical sensory aging is layer-specific": Figure 2-supplemental figure 1

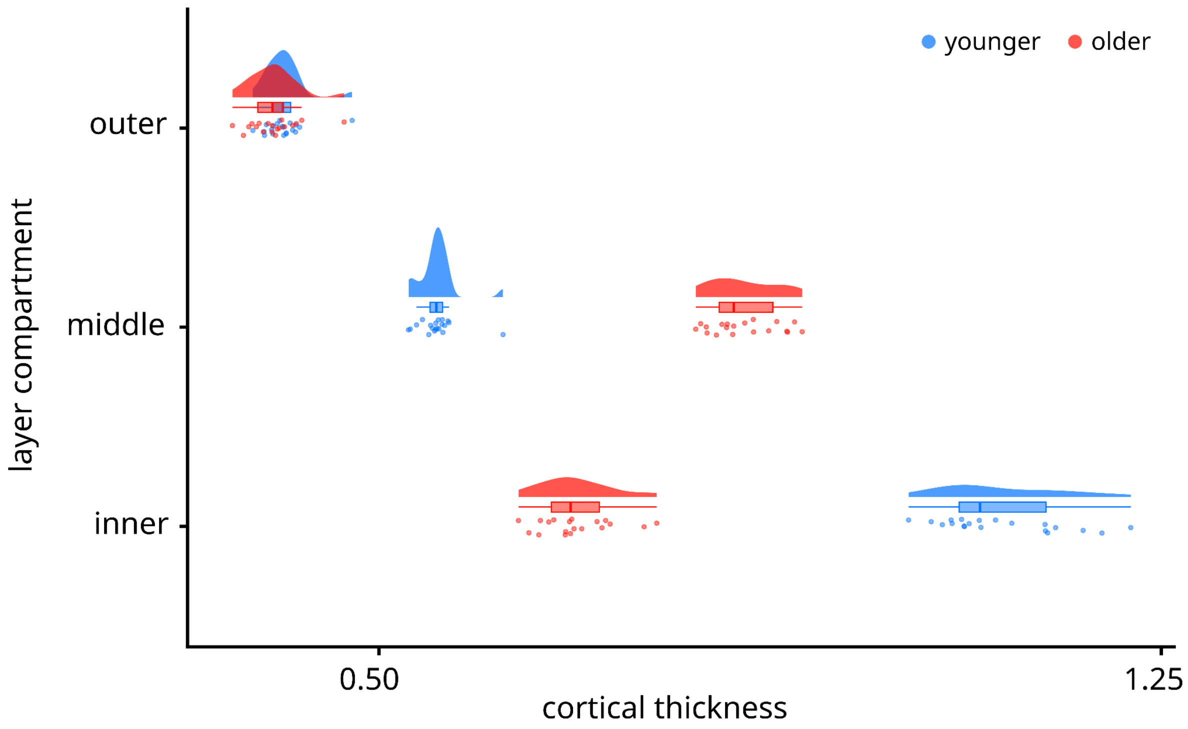


**Figure 2 - supplemental figure 1. Distribution of Cortical Thickness in Individual Layers of Human SI.** Individual data shown as colored dots: younger adults in blue, older adults in red. Box plots are drawn within the interquartile range (box), medians are shown as vertical lines, whiskers connect the minimum and the maximum with the lower and the upper quartiles.
