## Supplementary material for "Cortical sensory aging is layer-specific": Figure 2-supplemental figure 2

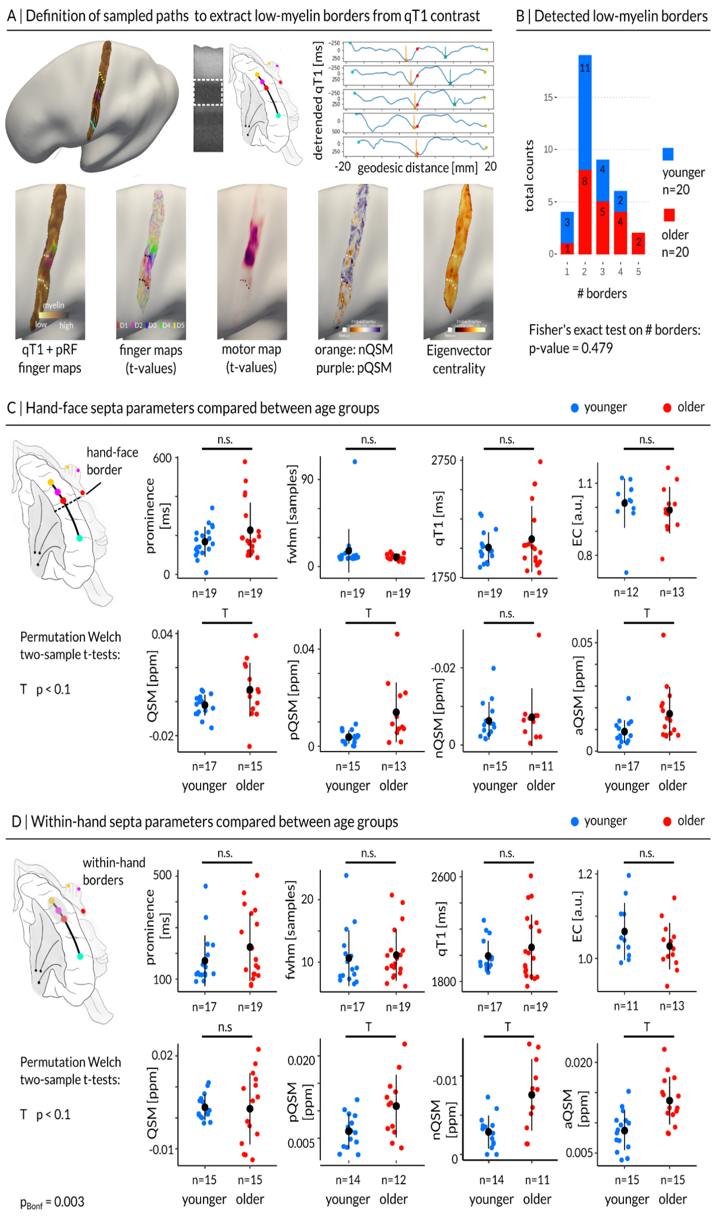


**Figure 2 - supplemental figure 2. Structural border results using an automated detection approach. (A)** Low-myelin borders (vertical lines) were detected based on detrended qT1 values sampled along multiple geodesic paths connecting the face representation (dots coloured in cyan) with the little finger representation (dots coloured in yellow). The analysis was based on qT1 values extracted from the middle layer compartment (dotted white line in myelin stain [remodeled according to Dinse et al.[4]]). Additional seeds were placed at the thumb representation (anchor to calculate geodesic distances, dots coloured in red) and the index finger representation (dots coloured in purple). Geodesic paths were sampled along the inferior-to-superior axis. Five paths were extracted from anterior to posterior within BA 3b using equal spaces between neighboring paths. Detected low myelin borders in BA 3b were back-projected to cortical surfaces (white dots in enlarged surface plots) and are shown together with different contrasts (from left to right): middle qT1 (lighter areas indicate lower myelin content) together with population receptive field (pRF) center location maps of individual fingers, finger activation maps (t-values) extracted from vibro-tactile blocked design paradigm, hand activation map (t-values) extracted from motor blocked design paradigm (i.e. individual finger movements), quantitative susceptibility map (QSM) indicative of diamagnetic (negative values coloured in orange, nQSM) and paramagnetic (positive values coloured in purple, pQSM) areas, connectivity map (Eigenvector centrality values, lighter colors indicate high connectivity). **(B)** Total counts of detected low myelin borders for younger (coloured in blue) and older (coloured in red) adults. Fisher's exact test indicated no difference in the number of detected borders between age groups. **(C)** Comparison of hand-face low-myelin border composition between age groups. There were no significant (n.s.) differences between age groups with respect to border prominence, full width at half maximum, qT1 intensity, Eigenvector centrality (EC), signed QSM values (QSM), positive QSM values (pQSM), negative QSM values (nQSM) or aQSM values. For n=2 participants no hand-face border could be detected. **(D)** Comparison of within-hand low-myelin border composition between age groups. There were no significant differences between age groups. Trends above Bonferroni-corrected threshold of p=0.003 (correcting for 16 tests) are marked by a T.
