## Supplementary material for "Cortical sensory aging is layer-specific": Figure 2-supplemental figure 3

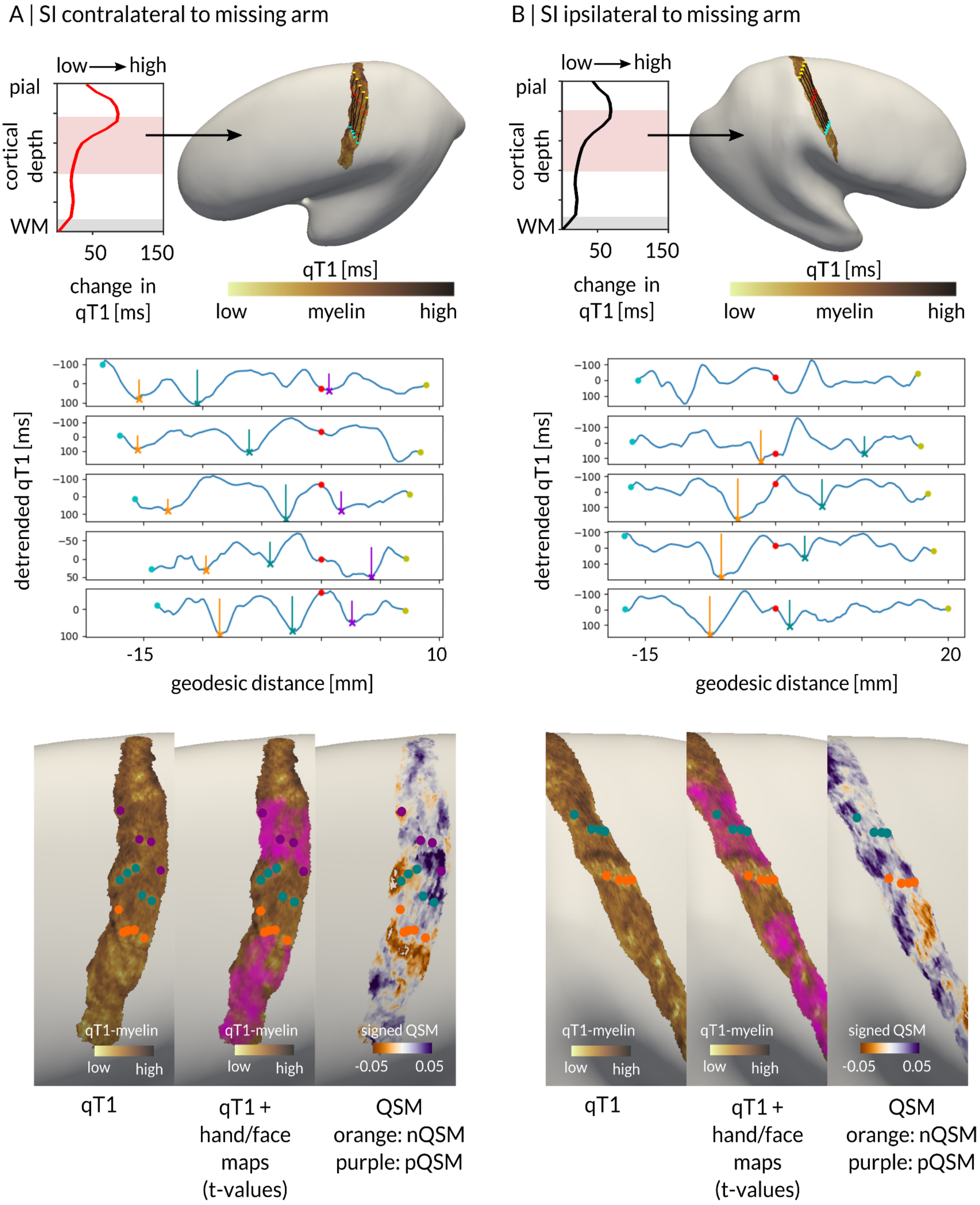


**Figure 2 - supplemental figure 3. Structural border results in SI (i.e., BA 3b) for the participant with congenital arm loss using an automated detection approach. (A)** Detected low-myelin borders in the hemisphere contralateral to the missing arm. The analysis was based on qT1 values extracted from the middle layer compartment (highlighted in light red) which were sampled from inferior to superior (connecting the upper face representation [dots coloured in cyan] with the superior border of the hand representation [dots coloured in yellow]) and from anterior to posterior (along multiple geodesic paths). Additional seeds were placed at the inferior border of the hand representation (anchor to calculate geodesic distances, dots coloured in red). Detected low myelin borders in the detrended qT1 signal (vertical coloured lines) in BA 3b were back-projected to cortical surfaces (coloured dots in enlarged surface plots) and are shown together with different contrasts (from left to right): middle qT1 map (lighter areas indicate lower myelin content), middle qT1 map together with hand and face activation maps (t-values, magenta coloured areas) localized by motor movements, quantitative susceptibility map (QSM) indicative of diamagnetic (negative values coloured in orange, nQSM) and paramagnetic (positive values coloured in purple, pQSM) areas. **(B)** Same as in (A) but calculated for the hemisphere ipsilateral to the missing arm.
