## Supplementary material for "Cortical sensory aging is layer-specific": Figure 2-supplemental figure 4

### A | BA 3b masks of younger adults

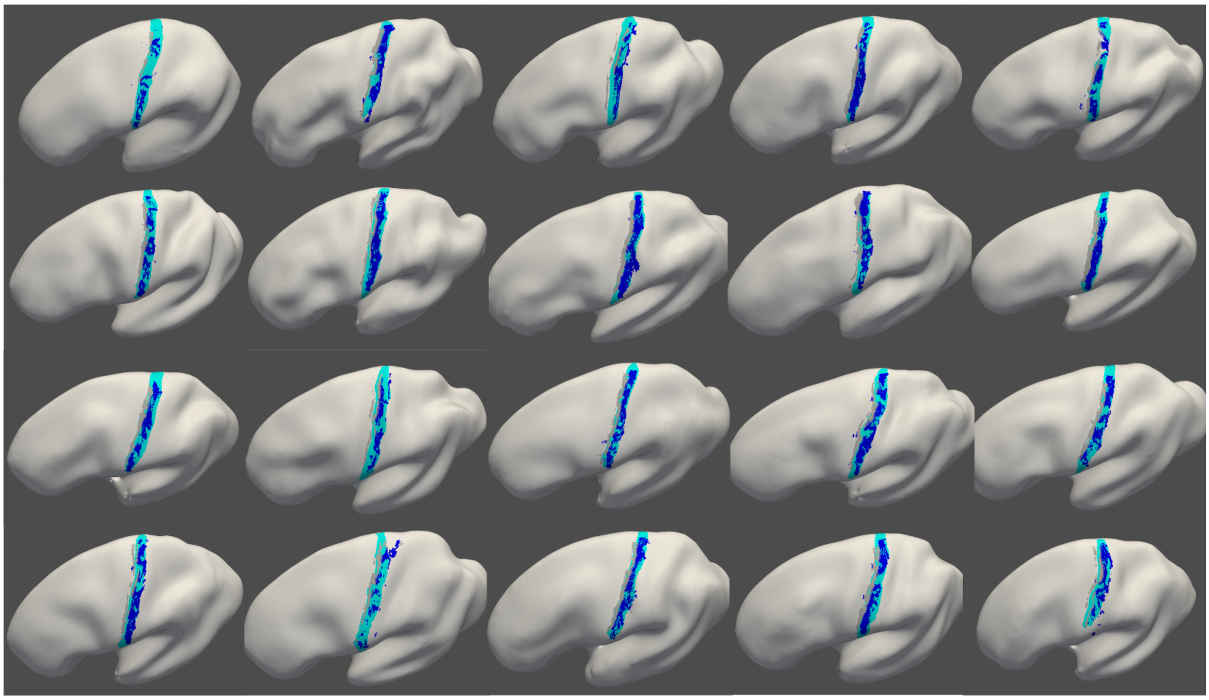

### B | BA 3b masks of older adults

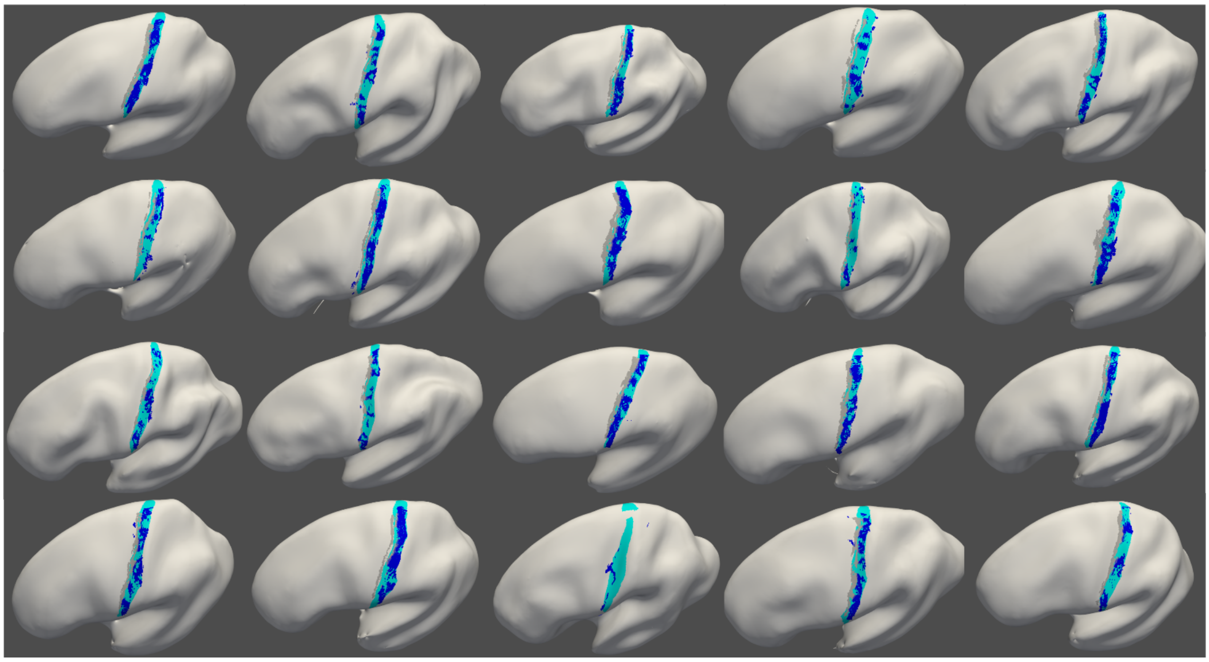

**Figure 2 - supplemental figure 4. Individual BA 3b masks in reference to co-registered Freesurfer labels. (A)** Shown are manual BA 3b masks of younger adults (which were used to delineate the region of interest) on individual inflated cortical surfaces (cyan) together with co-registered Freesurfer labels of BA 3b (dark blue) and BA 3a (gray). **(B)** Manual BA 3b masks of older adults (cyan) together with co-registered Freesurfer labels of BA 3b (dark blue) and BA 3a (gray).
