## Supplementary material for "Cortical sensory aging is layer-specific": Figure 2-supplemental table 1

**Figure 2 - supplemental table 1. Comparison of myelin, calcium, iron and mineralization content between age groups, fingers and layer compartments.** Permutation mixed-effects ANOVAs ("type III" sum of square) with between-subjects factor age (levels: younger, older) and within-subjects factors layer (levels: inner, middle, outer) and finger (D1, D2, D3, D4, D5) were performed on residual qT1 (myelin), nQSM (calcium), pQSM (iron) and aQSM (mineralization) values after controlling for finger map size (F=test statistic, df=degrees of freedom, p=parametric p-value, $df_{GG}$=Greenhouse-Geiser corrected degrees of freedom, $p_{GG}$=Greenhouse-Geiser corrected parametric p-value, $\eta_G^2$=effect size estimator generalized Eta-squared, $p_{perm}$=permutation p-value using the method by Kherad-Pajouh and Renaud[5] for non-spherical data with 100000 permutations). Please note that the minimum value of $p_{perm}$ is given by 1/number of permutations. For qT1 analysis n=2 participants (1 older and 1 younger), for nQSM n=12 participants (5 younger, 7 older), for pQSM n=4 participants (2 younger, 2 older), for aQSM n=2 participants (1 younger, 1 older) were excluded because of missing finger maps. Significant effects with Bonferroni-corrected p < 0.0125 (correcting for 4 ANOVAs) are marked by *. Trends above Bonferroni-corrected threshold are marked by a T.

| effect | *F* | *DFn, DFd* | *p* | *$DFn_{GG}$, $DFd_{GG}$* | *$p_{GG}$* | $\eta_G^2$ | *$p_{perm}$* |
|---|---|---|---|---|---|---|---|
| **qT1 (n=38)** | | | | | | | |
| age | 3.3 | 1, 36 | 0.077 T | 1.00, 36.00 | 0.077 T | 0.049 | 0.076 T |
| layer | 286.1 | 2,72 | $5.5x10^{-35}$ * | 1.05, 37.87 | $1.5x10^{-19}$ * | 0.690 | $<10^{-5}$ * |
| finger | 2.3 | 4, 144 | 0.059 T | 3.16, 113.64 | 0.075 T | 0.006 | 0.057 T |
| age x layer | 5.1 | 2, 72 | 0.009 * | 1.05, 37.87 | 0.028 T | 0.038 | 0.008 * |
| age x finger | 0.1 | 4, 144 | 0.976 | 3.16, 113.64 | 0.955 | $3.1^{-4}$ | 0.976 |
| layer x finger | 1.2 | 8, 288 | 0.295 | 3.57, 128.47 | 0.311 | 0.002 | 0.298 |
| age x layer x finger | 0.9 | 8, 288 | 0.541 | 3.57, 128.47 | 0.473 | 0.002 | 0.545 |
| **nQSM (n=22)** | | | | | | | |
| age | 4.0 | 1, 20 | 0.060 T | 1.00, 20.00 | 0.060 T | 0.039 | 0.059 T |
| layer | 7.1 | 1, 40 | 0.002 * | 1.44, 28.71 | 0.007 * | 0.070 | 0.002 * |
| finger | 0.2 | 4, 80 | 0.930 | 4.00, 80.00 | 0.930 | 0.003 | 0.929 |
| age x layer | 5.8 | 2, 40 | 0.006 * | 1.44, 28.71 | 0.014 T | 0.058 | 0.006 * |
| age x finger | 0.4 | 4, 80 | 0.806 | 4.00, 80.00 | 0.806 | 0.006 | 0.806 |
| layer x finger | 1.0 | 8, 160 | 0.462 | 4.58, 91.59 | 0.436 | 0.014 | 0.462 |

| | | | | | | | |
|---|---|---|---|---|---|---|---|
| age x layer x finger | 0.6 | 8, 160 | 0.795 | 4.58, 91.59 | 0.702 | 0.008 | 0.797 |
| **pQSM (n=30)** | | | | | | | |
| age | 35.3 | 1, 28 | $2.1x10^{-6}$ * | 1.00, 28.00 | $2.1x10^{-6}$ * | 0.262 | $<10^{-5}$ * |
| layer | 11.2 | 2, 56 | $8.2x10^{-5}$ * | 2.00, 56.00 | $8.2x10^{-5}$ * | 0.046 | $9.0x10^{-5}$ * |
| finger | 1.5 | 4, 112 | 0.205 | 2.36, 66.19 | 0.227 | 0.019 | 0.204 |
| age x layer | 1.4 | 2, 56 | 0.257 | 2.00, 56.00 | 0.257 | 0.006 | 0.258 |
| age x finger | 1.2 | 4, 112 | 0.331 | 2.36, 66.19 | 0.325 | 0.015 | 0.334 |
| layer x finger | 0.4 | 8, 224 | 0.892 | 4.45, 124.52 | 0.794 | 0.004 | 0.892 |
| age x layer x finger | 1.0 | 8, 224 | 0.463 | 4.45, 124.52 | 0.435 | 0.008 | 0.467 |
| **aQSM (n=32)** | | | | | | | |
| age | 43.8 | 1, 30 | $2.5x10^{-7}$ * | 1.00, 30.00 | $2.5x10^{-7}$ * | 0.291 | $<10^{-5}$ * |
| layer | 8.2 | 2, 60 | $7.3x10^{-4}$ * | 2.00, 60.00 | $7.3x10^{-4}$ * | 0.040 | $8.7x10^{-4}$ * |
| finger | 1.5 | 4, 120 | 0.194 | 2.74, 82.18 | 0.213 | 0.015 | 0.195 |
| age x layer | 0.4 | 2, 60 | 0.649 | 2.00, 60.00 | 0.649 | 0.002 | 0.650 |
| age x finger | 1.1 | 4, 120 | 0.342 | 2.74, 82.18 | 0.336 | 0.011 | 0.344 |
| layer x finger | 1.4 | 8, 240 | 0.179 | 4.53, 135.92 | 0.217 | 0.012 | 0.178 |
| age x layer x finger | 0.3 | 8, 240 | 0.962 | 4.53, 135.92 | 0.892 | 0.003 | 0.962 |
