## Supplementary material for "Cortical sensory aging is layer-specific": Figure 2-supplemental table 2

**Figure 2 - supplemental table 2. Comparison of qT1, nQSM, pQSM and aQSM values between age groups, fingers and layer compartments using the layer definition of younger adults for both age groups.** Permutation mixed-effects ANOVAs ("type III" sum of square) with between-subjects factor age (levels: younger, older) and within-subjects factors layer (levels: inner, middle, outer) and finger (D1, D2, D3, D4, D5) were performed on residual qT1 (myelin), nQSM (calcium), pQSM (iron) and aQSM (mineralization) values after controlling for finger map size (F=test statistic, df=degrees of freedom, p=parametric p-value, $df_{GG}$=Greenhouse-Geiser corrected degrees of freedom, $p_{GG}$=Greenhouse-Geiser corrected parametric p-value, $\eta_G^2$=effect size estimator generalized Eta-squared, $p_{perm}$=permutation p-value using the method by Kherad-Pajouh and Renaud[5] for non-spherical data with 100000 permutations. Please note that the minimum value of $p_{perm}$ is given by 1/number of permutations. For qT1 analysis n=2 participants (1 older, 1 younger), for nQSM n=13 participants (5 younger, 8 older), for pQSM n=4 participants (2 younger, 2 older), for aQSM n=2 participants (1 younger, 1 older) were excluded because of missing finger maps. Significant effects with Bonferroni-corrected p < 0.0125 (correcting for 4 ANOVAs) are marked by *. Trends above Bonferroni-corrected threshold are marked by a T.

| effect | *F* | *DFn, DFd* | *p* | *$DFn_{GG}$, $DFd_{GG}$* | *$p_{GG}$* | $\eta_G^2$ | *$p_{perm}$* |
|---|---|---|---|---|---|---|---|
| **qT1 (n=38)** | | | | | | | |
| age | 0.1 | 1, 36 | 0.765 | 1.00, 36.00 | 0.765 | 0.002 | 0.771 |
| layer | 277.4 | 2, 72 | $1.5 \times 10^{-34}$ * | 1.06, 38.31 | $1.6 \times 10^{-19}$ * | 0.654 | $<10^{-5}$ * |
| finger | 2.4 | 4, 144 | 0.056 T | 3.06, 110.07 | 0.074 T | 0.006 | 0.055 T |
| age x layer | 2.8 | 2, 72 | 0.070 T | 1.06, 38.31 | 0.102 | 0.019 | 0.069 T |
| age x finger | 0.2 | 4, 144 | 0.940 | 3.06, 110.07 | 0.902 | 0.001 | 0.940 |
| layer x finger | 1.2 | 8, 288 | 0.275 | 3.70, 133.30 | 0.297 | 0.002 | 0.276 |
| age x layer x finger | 0.7 | 8, 288 | 0.648 | 3.70, 133.30 | 0.551 | 0.001 | 0.650 |
| **nQSM (n=21)** | | | | | | | |
| age | 11.2 | 1, 19 | 0.003 * | 1.00, 19.00 | 0.003 * | 0.091 | 0.003 * |
| layer | 15.1 | 2, 38 | $1.5 \times 10^{-5}$ * | 1.36, 25.81 | 0.002 * | 0.147 | $3.0 \times 10^{-5}$ * |
| finger | 0.3 | 4, 76 | 0.878 | 4.00, 76.00 | 0.878 | 0.005 | 0.878 |
| age x layer | 2.0 | 2, 38 | 0.146 | 1.36, 25.81 | 0.163 | 0.023 | 0.147 |
| age x finger | 1.0 | 4, 76 | 0.419 | 4.00, 76.00 | 0.419 | 0.016 | 0.421 |
| layer x finger | 1.6 | 8, 152 | 0.123 | 4.10, 77.84 | 0.176 | 0.025 | 0.124 |

| | | | | | | | |
|---|---|---|---|---|---|---|---|
| age x layer x finger | 0.8 | 8, 152 | 0.645 | 4.10, 77.84 | 0.562 | 0.012 | 0.645 |
| **pQSM (n=30)** | | | | | | | |
| age | 37.9 | 1, 28 | $1.2x10^{-6}$ * | 1.00, 28.00 | $1.2x10^{-6}$ * | 0.270 | $<10^{-5}$ * |
| layer | 5.9 | 2, 56 | 0.005 * | 2.00, 56.00 | 0.005 * | 0.022 | 0.005 * |
| finger | 1.4 | 4, 112 | 0.227 | 2.44, 68.41 | 0.243 | 0.020 | 0.228 |
| age x layer | 0.1 | 2, 56 | 0.917 | 2.00, 56.00 | 0.917 | $3.3x10^{-4}$ | 0.917 |
| age x finger | 1.1 | 4, 112 | 0.354 | 2.44, 68.41 | 0.343 | 0.015 | 0.356 |
| layer x finger | 0.5 | 8, 224 | 0.876 | 4.41, 123.50 | 0.775 | 0.004 | 0.878 |
| age x layer x finger | 1.0 | 8, 224 | 0.421 | 4.41, 123.50 | 0.404 | 0.008 | 0.423 |
| **aQSM (n=32)** | | | | | | | |
| age | 45.6 | 1, 30 | $1.7x10^{-7}$ * | 1.00, 30.00 | $1.7x10^{-7}$ * | 0.310 | $<10^{-5}$ * |
| layer | 6.2 | 2, 60 | 0.004 * | 2.00, 60.00 | 0.004 * | 0.026 | 0.004 * |
| finger | 1.2 | 4, 120 | 0.298 | 2.87, 86.00 | 0.300 | 0.014 | 0.300 |
| age x layer | 0.5 | 2, 60 | 0.596 | 2.00, 60.00 | 0.596 | 0.002 | 0.598 |
| age x finger | 0.8 | 4, 120 | 0.528 | 2.87, 86.00 | 0.492 | 0.009 | 0.527 |
| layer x finger | 1.8 | 8, 240 | 0.083 T | 4.71, 141.23 | 0.127 | 0.014 | 0.082 T |
| age x layer x finger | 0.4 | 8, 240 | 0.918 | 4.71, 141.23 | 0.836 | 0.003 | 0.919 |
