## Supplementary material for "Cortical sensory aging is layer-specific": Figure 2-supplemental table 3

**Figure 2 - supplemental table 3. Comparison of qT1, nQSM, pQSM and aQSM values between age groups, body parts (hand, face) and layer compartments.** Permutation mixed-effects ANOVAs ("type III" sum of square) with between-subjects factor age (levels: younger, older) and within-subjects factors layer (levels: inner, middle, outer) and body part (hand, face) were performed on residual qT1 (myelin), nQSM (calcium), pQSM (iron) and aQSM (mineralization) values after controlling for body part map size (F=test statistic, df=degrees of freedom, p=parametric p-value, $df_{GG}$=Greenhouse-Geiser corrected degrees of freedom, $p_{GG}$=Greenhouse-Geiser corrected parametric p-value, $\eta_G^2$=effect size estimator generalized Eta-squared, $p_{perm}$=permutation p-value using the method by Kherad-Pajouh and Renaud[5] for non-spherical data with 100000 permutations. Please note that the minimum value of $p_{perm}$ is given by 1/number of permutations. Significant effects with Bonferroni-corrected p < 0.0125 (correcting for 4 ANOVAs) are marked by *. Trends above Bonferroni-corrected threshold are marked by a T.

| effect | *F* | *DFn, DFd* | *p* | *$DFn_{GG}$, $DFd_{GG}$* | *$p_{GG}$* | $\eta_G^2$ | *$p_{perm}$* |
|---|---|---|---|---|---|---|---|
| **qT1 (n=34)** | | | | | | | |
| age | 1.1 | 1, 32 | 0.312 | 1.00, 32.00 | 0.312 | 0.022 | 0.316 |
| layer | 416.8 | 2,64 | $2.0\times10^{-37}$ * | 1.04, 33.25 | $1.3\times10^{-20}$ * | 0.778 | $<10^{-5}$ * |
| body part | 0.1 | 1, 32 | 0.735 | 1.00, 32.00 | 0.735 | $1.2\times10^{-4}$ | 0.737 |
| age x layer | 15.3 | 2, 64 | $3.7\times10^{-6}$ * | 1.04, 33.25 | $3.7\times10^{-4}$ * | 0.114 | $<10^{-5}$ * |
| age x body part | 1.5 | 1, 32 | 0.231 | 1.00, 32.00 | 0.231 | 0.001 | 0.232 |
| layer x body part | 5.3 | 2, 64 | 0.007 * | 1.21, 38.84 | 0.021 T | 0.002 | 0.007 * |
| age x layer x body part | 2.1 | 2, 64 | 0.132 | 1.21, 38.84 | 0.153 | 0.001 | 0.131 |
| **nQSM (n=30)** | | | | | | | |
| age | 32.8 | 1, 28 | $3.7\times10^{-6}$ * | 1.00, 28.00 | $3.7\times10^{-6}$ * | 0.313 | $<10^{-5}$ * |
| layer | 36.9 | 2, 56 | $6.0\times10^{-11}$ * | 1.23, 34.50 | $1.4\times10^{-7}$ * | 0.315 | $<10^{-5}$ * |
| body part | 2.7 | 1, 28 | 0.113 | 1.00, 28.00 | 0.113 | 0.008 | 0.112 |
| age x layer | 2.3 | 2, 56 | 0.107 | 1.23, 34.5 | 0.131 | 0.028 | 0.106 |
| age x body part | 8.5 | 1, 28 | 0.007 * | 1.00, 28.0 | 0.007 * | 0.026 | 0.006 * |
| layer x body part | 4.4 | 2, 56 | 0.016 T | 1.34, 37.4 | 0.032 T | 0.027 | 0.017 T |
| age x layer x body part | 3.1 | 2, 56 | 0.053 T | 1.34, 37.4 | 0.075 T | 0.019 | 0.052 T |

| | | | | | | | |
|---|---|---|---|---|---|---|---|
| **pQSM (n=30)** | | | | | | | |
| age | 45.3 | 1, 28 | $2.6x10^{-7}$ * | 1.00, 28.00 | $2.6x10^{-7}$ * | 0.478 | $<10^{-5}$ * |
| layer | 32.0 | 2, 56 | $5.4x10^{-10}$ * | 1.11, 31.01 | $1.7x10^{-6}$ * | 0.229 | $<10^{-5}$ * |
| body part | 18.3 | 1, 28 | $2.0x10^{-4}$ * | 1.00, 28.00 | $2.0x10^{-4}$ * | 0.053 | $1.4x10^{-4}$ * |
| age x layer | 16.2 | 2, 56 | $2.8x10^{-6}$ * | 1.11, 31.01 | $2.3x10^{-4}$ * | 0.131 | $<10^{-5}$ * |
| age x body part | 0.01 | 1, 28 | 0.939 | 1.00, 28.00 | 0.939 | $1.8x10^{-5}$ | 0.939 |
| layer x body part | 7.6 | 2, 56 | 0.001 * | 1.29, 36.14 | 0.006 * | 0.023 | 0.001 * |
| age x layer x body part | 0.8 | 2, 56 | 0.440 | 1.29, 36.14 | 0.396 | 0.003 | 0.446 |
| **aQSM (n=30)** | | | | | | | |
| age | 83.2 | 1, 28 | $7.1x10^{-10}$ * | 1.00, 28.00 | $7.1x10^{-10}$ * | 0.622 | $<10^{-5}$ * |
| layer | 97.3 | 2, 56 | $6.0^{-19}$ * | 1.27, 35.46 | $7.7^{-13}$ * | 0.463 | $<10^{-5}$ * |
| body part | 14.6 | 1, 28 | $6.8x10^{-4}$ * | 1.00, 28.00 | $6.8x10^{-4}$ * | 0.046 | $4.7x10^{-4}$ * |
| age x layer | 21.7 | 2, 56 | $1.1x10^{-7}$ * | 1.27, 35.46 | $1.3x10^{-5}$ * | 0.161 | $<10^{-5}$ * |
| age x body part | 0.2 | 1, 28 | 0.657 | 1.00, 28.00 | 0.657 | 0.001 | 0.659 |
| layer x body part | 10.5 | 2, 56 | $1.4x10^{-4}$ * | 1.42, 39.63 | $8.4x10^{-4}$ * | 0.038 | $9.0x10^{-5}$ * |
| age x layer x body part | 4.3 | 2, 56 | 0.018 T | 1.42, 39.63 | 0.032 T | 0.016 | 0.017 T |
