## Supplementary material for "Cortical sensory aging is layer-specific": Figure 2-supplemental table 4

**Figure 2 - supplemental table 4. Post-hoc comparisons to follow up significant main effects, significant interaction effects and trends on qT1, nQSM, pQSM and aQSM related to cortical fields (hand, face) and layer compartments.** Bootstrap Welch paired-sample t-tests were performed on qT1 (myelin) (n=16 younger adults, n=18 older adults), nQSM (calcium) (n=14 younger adults, n=16 older adults), pQSM (iron) (n=14 younger adults, n=16 older adults) and aQSM (mineralization) (n=14 younger adults, n=16 older adults) values to follow up significant main effects of body part (tested differences: hand-face) and interactions between body part and layer (tested differences: hand outer - face outer, hand middle - face middle, hand inner - face inner), Bootstrap Welch two-sample t-tests were performed to follow up significant interaction effects between age (younger, older) and body part (tested differences: younger hand - older hand, younger face - older face) as well as between age, body part and layer (y=younger, ol=older; o=outer, m=middle, i=inner). Results are given as mean difference values (Mean) and standard errors (SE) in milliseconds (for qT1) or in ppm (for QSM-based estimates), test statistic (t), degrees of freedom (df), p-value (p), bootstrapped p-value ($p_{boot}$), 95% confidence interval (CI), bootstrapped 95% confidence interval ($CI_{boot}$). Number of bootstrap samples=100000, minimum value of $p_{boot}$=1/number of bootstrap samples. Significant effects with Bonferroni-corrected p < 0.002 (correcting for 25 post-hoc tests) are marked by *. Trends above Bonferroni-corrected threshold are marked by a T.

| comparison | Mean ± SE | *t* | *df* | *p* | $p_{boot}$ | *CI* | $CI_{boot}$ |
|---|---|---|---|---|---|---|---|
| **qT1 (n=34)** | | | | | | | |
| face outer > hand outer | 5.4 ± 12.3 | 0.4 | 33 | 0.665 | 0.652 | -19.7, 30.5 | -17.9, 30.0 |
| face middle > hand middle | 16.1 ± 7.4 | 2.2 | 33 | 0.036 T | 0.037 T | 1.1, 31.1 | 1.7, 30.2 |
| face inner < hand inner | -11.8 ± 7.5 | -1.6 | 33 | 0.128 | 0.114 | -27.1, 3.6 | -26.5, 2.5 |
| **nQSM (n=30)** | | | | | | | |
| younger face > older face | 0.004 ± $6.3\times10^{-4}$ | 6.8 | 20.7 | $9.9\times10^{-4}$ * | $<10^{-5}$ * | $3.0\times10^{-3}$, $5.7\times10^{-3}$ | $3.1\times10^{-3}$, $5.5\times10^{-3}$ |
| younger hand > older hand | 0.003 ± $6.5\times10^{-4}$ | 4.1 | 19.8 | $6.0\times10^{-4}$ * | $5.6\times10^{-4}$ * | $1.2\times10^{-3}$, $4.0\times10^{-3}$ | $1.4\times10^{-3}$, $3.9\times10^{-3}$ |
| y face o > ol face o | 0.005 ± $1.0\times10^{-3}$ | 5.2 | 23.9 | $2.7\times10^{-5}$ * | $6.0\times10^{-5}$ * | $3.2\times10^{-3}$, $7.4\times10^{-3}$ | $3.3\times10^{-3}$, $7.2\times10^{-3}$ |
| y face m > ol face m | 0.005 ± $8.1\times10^{-4}$ | 6.7 | 21.0 | $1.3\times10^{-6}$ * | $<10^{-5}$ * | $3.7\times10^{-3}$, $7.1\times10^{-3}$ | $3.9\times10^{-3}$, $6.9\times10^{-3}$ |
| y face i > o face i | 0.002 ± $7.0\times10^{-4}$ | 3.2 | 28.0 | 0.003 T | 0.004 T | $8.3\times10^{-4}$, $3.7\times10^{-3}$ | $9.4\times10^{-4}$, $3.6\times10^{-3}$ |
| y hand o > ol hand o | 0.002 ± $1.2\times10^{-3}$ | 1.6 | 17.5 | 0.129 | 0.102 | $-6.2\times10^{-4}$, $4.4\times10^{-3}$ | $-2.8\times10^{-4}$, $4.3\times10^{-3}$ |
| y hand m > ol hand m | 0.004 ± $8.2\times10^{-4}$ | 4.3 | 19.8 | $3.4\times10^{-4}$ * | $1.8\times10^{-4}$ * | $1.8\times10^{-3}$, $5.3\times10^{-3}$ | $2.0\times10^{-3}$, $5.1\times10^{-3}$ |
| y hand i > ol hand i | 0.002 ± $9.3\times10^{-4}$ | 2.6 | 23.3 | 0.016 T | 0.007 T | $4.9\times10^{-4}$, $4.3\times10^{-3}$ | $7.3\times10^{-4}$, $4.2\times10^{-3}$ |
| **pQSM (n=30)** | | | | | | | |
| face < hand | -0.001 ± $3.4\times10^{-4}$ | -4.4 | 29 | $1.5\times10^{-4}$ * | $1.5\times10^{-4}$ * | $-2.2\times10^{-3}$, $-7.9\times10^{-4}$ | $-2.1\times10^{-3}$, $-8.1\times10^{-4}$ |

| | | | | | | | |
|---|---|---|---|---|---|---|---|
| face outer < hand outer | $-8.8\times10^{-5} \pm 6.9\times10^{-4}$ | -0.1 | 29 | 0.900 | 0.915 | $-1.5\times10^{-3}$, $1.3\times10^{-3}$ | $-1.4\times10^{-3}$, $1.3\times10^{-3}$ |
| face middle < hand middle | $-0.002 \pm 2.9\times10^{-4}$ | -7.3 | 29 | $4.8\times10^{-8}$ * | $<10^{-5}$ * | $-2.7\times10^{-3}$, $-1.5\times10^{-3}$ | $-2.6\times10^{-3}$, $-1.5\times10^{-3}$ |
| face inner < hand inner | $-0.002 \pm 4.0\times10^{-4}$ | -5.6 | 29 | $4.3\times10^{-6}$ * | $<10^{-5}$ * | $-3.1\times10^{-3}$, $-1.4\times10^{-3}$ | $-3.1\times10^{-3}$, $-1.5\times10^{-3}$ |
| **aQSM (n=30)** | | | | | | | |
| face < hand | $-0.001 \pm 2.6\times10^{-4}$ | -3.8 | 29 | $6.0\times10^{-4}$ * | $2.4\times10^{-4}$ * | $-1.5\times10^{-3}$, $-4.7\times10^{-4}$ | $-1.5\times10^{-3}$, $-5.0\times10^{-4}$ |
| face outer > hand outer | $3.4\times10^{-4} \pm 4.8\times10^{-4}$ | 0.7 | 29 | 0.490 | 0.491 | $-6.5\times10^{-4}$, $1.3\times10^{-3}$ | $-5.9\times10^{-4}$, $1.3\times10^{-3}$ |
| face middle < hand middle | $-1.6\times10^{-3} \pm 3.1\times10^{-4}$ | -5.0 | 29 | $2.3\times10^{-5}$ * | $<10^{-5}$ * | $-2.2\times10^{-3}$, $-9.4\times10^{-4}$ | $-2.2\times10^{-3}$, $-1.0\times10^{-3}$ |
| face inner < hand inner | $-1.7\times10^{-3} \pm 3.6\times10^{-4}$ | -4.8 | 29 | $3.9\times10^{-5}$ * | $<10^{-5}$ * | $-2.5\times10^{-3}$, $-1.0\times10^{-3}$ | $-2.4\times10^{-3}$, $-1.1\times10^{-3}$ |
| y face o < ol face o | $-9.2\times10^{-3} \pm 9.3\times10^{-4}$ | -9.9 | 23.1 | $8.0\times10^{-10}$ * | $<10^{-5}$ * | $-1.1\times10^{-2}$, $-7.3\times10^{-3}$ | $-1.1\times10^{-2}$, $-7.6\times10^{-3}$ |
| y face m < ol face m | $-5.5\times10^{-3} \pm 6.5\times10^{-4}$ | -8.5 | 22.6 | $1.7\times10^{-8}$ * | $<10^{-5}$ * | $-6.9\times10^{-3}$, $-4.2\times10^{-3}$ | $-6.7\times10^{-3}$, $-4.3\times10^{-3}$ |
| y face i < ol face i | $-3.1\times10^{-3} \pm 5.5\times10^{-4}$ | -5.6 | 25.6 | $7.3\times10^{-6}$ * | $<10^{-5}$ * | $-4.2\times10^{-3}$, $-1.9\times10^{-3}$ | $-4.1\times10^{-3}$, $-2.0\times10^{-3}$ |
| y hand o < ol hand o | $-7.0\times10^{-3} \pm 1.1\times10^{-3}$ | -6.5 | 18.2 | $3.6\times10^{-6}$ * | $<10^{-5}$ * | $-9.7\times10^{-3}$, $-5.0\times10^{-3}$ | $-9.7\times10^{-3}$, $-5.4\times10^{-3}$ |
| y hand m < ol hand m | $-5.9\times10^{-3} \pm 8.0\times10^{-4}$ | -7.4 | 22.8 | $1.6\times10^{-7}$ * | $<10^{-5}$ * | $-7.6\times10^{-3}$, $-4.3\times10^{-3}$ | $-7.4\times10^{-3}$, $-4.4\times10^{-3}$ |
| y hand i < ol hand i | $-3.8\times10^{-3} \pm 7.3\times10^{-4}$ | -5.2 | 21.1 | $3.7\times10^{-5}$ * | $<10^{-5}$ * | $-5.3\times10^{-3}$, $-2.3\times10^{-3}$ | $-5.2\times10^{-3}$, $-2.4\times10^{-3}$ |
