## Supplementary material for "Cortical sensory aging is layer-specific": Figure 2-supplemental table 5

**Figure 2 - supplemental table 5. Comparison of qT1, nQSM, pQSM and aQSM values between age groups and layer compartments.** Permutation mixed-effects ANOVAs ("type III" sum of square) with between-subjects factor age (levels: younger, older) and within-subjects factor layer (levels: inner, middle, outer) were performed on qT1 (myelin), nQSM (calcium), pQSM (iron) and aQSM (mineralization) values (F=test statistic, df=degrees of freedom, p=parametric p-value, $df_{GG}$=Greenhouse-Geiser corrected degrees of freedom, $p_{GG}$=Greenhouse-Geiser corrected parametric p-value, $\eta_G^2$=effect size estimator generalized Eta-squared, $p_{perm}$=permutation p-value using the method by Kherad-Pajouh & Renaud[5] for non-spherical data with 100000 permutations, minimum value of $p_{perm}$=1/number of permutations). Significant effects with Bonferroni-corrected p < 0.0125 (correcting for 4 ANOVAs) are marked by *.

| effect | *F* | *DFn, DFd* | *p* | *DFn<sub>GG</sub>, DFd<sub>GG</sub>* | *p<sub>GG</sub>* | $\eta_G^2$ | $p_{perm}$ |
|---|---|---|---|---|---|---|---|
| **qT1 (n=40)** | | | | | | | |
| age | 2.01 | 1, 38 | 0.164 | 1.00, 38.00 | 0.164 | 0.034 | 0.167 |
| layer | 338.01 | 2, 76 | $1.5\times10^{-38}$ * | 1.06, 40.46 | $1.1\times10^{-21}$ * | 0.747 | $<10^{-5}$ * |
| age x layer | 7.24 | 2, 76 | 0.001 * | 1.06, 40.46 | 0.009 * | 0.059 | $7.5\times10^{-4}$ * |
| **nQSM (n=34)** | | | | | | | |
| age | 11.01 | 1, 32 | 0.002 * | 1.00, 32.00 | 0.002 * | 0.135 | 0.002 * |
| layer | 15.34 | 2, 64 | $3.6\times10^{-6}$ * | 1.16, 37.24 | $2.0\times10^{-4}$ * | 0.207 | $<10^{-5}$ * |
| age x layer | 0.20 | 2, 64 | 0.822 | 1.16, 37.24 | 0.698 | 0.003 | 0.824 |
| **pQSM (n=34)** | | | | | | | |
| age | 22.68 | 1, 32 | $4.0\times10^{-5}$ * | 1.00, 32.00 | $4.0\times10^{-5}$ * | 0.301 | $5.0\times10^{-5}$ * |
| layer | 12.76 | 2, 64 | $2.2\times10^{-5}$ * | 1.36, 43.45 | $2.7\times10^{-4}$ * | 0.135 | $3.0\times10^{-5}$ * |
| age x layer | 1.06 | 2, 64 | 0.351 | 1.36, 43.45 | 0.330 | 0.013 | 0.353 |
| **aQSM (n=34)** | | | | | | | |
| age | 26.70 | 1, 32 | $1.2\times10^{-5}$ * | 1.00, 32.00 | $1.2\times10^{-5}$ * | 0.349 | $2.0\times10^{-5}$ * |
| layer | 17.65 | 2, 64 | $7.8\times10^{-7}$ * | 1.55, 49.71 | $9.2\times10^{-6}$ * | 0.164 | $<10^{-5}$ * |
| age x layer | 1.16 | 2, 64 | 0.321 | 1.55, 49.71 | 0.312 | 0.013 | 0.323 |
