## Supplementary material for "Cortical sensory aging is layer-specific": Figure 2-supplemental table 6

**Figure 2 - supplemental table 6. Post-hoc comparisons to follow up significant layer-specific age differences in qT1, nQSM, pQSM and aQSM values.** Bootstrap Welch paired-sample t-tests were performed on qT1 (myelin), nQSM (calcium), pQSM (iron) and aQSM (mineralization) values to follow up significant main effects of layer (tested differences: outer-middle, middle-inner), Bootstrap Welch two-sample t-tests were performed to follow up significant main effects of layer (inner, middle, outer; tested differences; outer-middle, middle-inner) and significant interaction effects between age (younger, older) and layer (tested differences: younger outer-older outer, younger middle-older middle, younger inner-older inner). Results are given as mean difference values (Mean) and standard errors (SE) in milliseconds (for qT1) or ppm (for QSM-based estimates), test statistic (t), degrees of freedom (df), p-value (p), bootstrapped p-value ($p_{boot}$), 95% confidence interval (CI), bootstrapped 95% confidence interval ($CI_{boot}$). Number of bootstrap samples=100000, minimum value of $p_{boot}$=1/number of bootstrap samples. Significant effects with Bonferroni-corrected p < 0.0045 (correcting for 11 post-hoc tests) are marked by *.

| comparison | Mean ± SE | *t* | *df* | *p* | $p_{boot}$ | *CI* | $CI_{boot}$ |
|---|---|---|---|---|---|---|---|
| **qT1 (n=40)** | | | | | | | |
| outer > middle | 279.8 ± 22.3 | 12.6 | 39.0 | $2.8 \times 10^{-15}$ * | $<10^{-5}$ * | 234.8, 324.9 | 240.2, 325.8 |
| middle > inner | 236.9 ± 8.6 | 27.6 | 39.0 | $3.2 \times 10^{-27}$ * | $<10^{-5}$ * | 219.5, 254.2 | 221.6, 254.8 |
| younger outer < older outer | -41.2 ± 61.3 | -0.7 | 25.0 | 0.508 | 0.467 | -167.3, 85.0 | -164.7, 69.7 |
| younger middle > older middle | 91.2 ± 25.4 | 3.6 | 30.8 | 0.001 * | 0.002 * | 39.3, 143.1 | 41.6, 139.2 |
| younger inner > older inner | 88.7 ± 19.3 | 4.6 | 36.8 | $4.8 \times 10^{-5}$ * | $4.0 \times 10^{-5}$ * | 49.7, 127.8 | 51.6, 125.2 |
| **nQSM (n=34)** | | | | | | | |
| outer < middle | $-2.9 \times 10^{-3} \pm 3.2 \times 10^{-4}$ | -9.2 | 33.0 | $1.4 \times 10^{-10}$ * | $<10^{-5}$ * | $-3.5 \times 10^{-3}$, $-2.3 \times 10^{-3}$ | $-3.5 \times 10^{-3}$, $-2.3 \times 10^{-3}$ |
| middle > inner | $3.4 \times 10^{-4} \pm 5.4 \times 10^{-4}$ | 0.6 | 33.0 | 0.541 | 0.569 | $-7.7 \times 10^{-4}$, $1.4 \times 10^{-3}$ | $-7.5 \times 10^{-4}$, $1.3 \times 10^{-3}$ |
| **pQSM (n=34)** | | | | | | | |
| outer > middle | $5.0 \times 10^{-4} \pm 6.3 \times 10^{-4}$ | 0.8 | 33 | 0.428 | 0.392 | $-7.7 \times 10^{-4}$, $1.8 \times 10^{-3}$ | $-6.6 \times 10^{-4}$, $1.8 \times 10^{-3}$ |
| middle > inner | $2.2 \times 10^{-3} \pm 3.5 \times 10^{-4}$ | 6.3 | 33 | $4.5 \times 10^{-7}$ * | $<10^{-5}$ * | $1.5 \times 10^{-3}$, $2.9 \times 10^{-3}$ | $1.6 \times 10^{-3}$, $2.9 \times 10^{-3}$ |

| | | | | | | | |
|---|---|---|---|---|---|---|---|
| **aQSM (n=34)** | | | | | | | |
| outer > middle | $1.1x10^{-3}$ ± $4.5x10^{-4}$ | 2.5 | 33 | 0.019 T | 0.024 T | $1.9x10^{-4}$, $2.0x10^{-3}$ | $2.3x10^{-4}$, $2.0x10^{-3}$ |
| middle > inner | $1.4x10^{-3}$ ± $3.1x10^{-4}$ | 4.5 | 33 | $8.5x10^{-5}$ * | <$10^{-5}$ * | $7.7x10^{-4}$, $2.0x10^{-3}$ | $8.2x10^{-4}$, $2.0x10^{-3}$ |
