## Supplementary material for "Cortical sensory aging is layer-specific": Figure 2-supplemental table 7

**Figure 2 - supplemental table 7. Comparison of qT1, nQSM, pQSM and aQSM values of the SI hand area between age groups and layer compartments using the layer definition of younger adults for both age groups.** Permutation mixed-effects ANOVAs ("type III" sum of square) with between-subjects factor age (levels: younger, older) and within-subjects factor layer (levels: inner, middle, outer) were performed on qT1 (myelin) (n=20 younger adults, n=20 older adults), nQSM (calcium) (n=18 younger adults, n=16 older adults), pQSM (iron) (n=18 younger adults, n=16 older adults) and aQSM (mineralization) (n=18 younger adults, n=16 older adults) values (F=test statistic, df=degrees of freedom, p=parametric p-value, $df_{GG}$=Greenhouse-Geiser corrected degrees of freedom, $p_{GG}$=Greenhouse-Geiser corrected parametric p-value, $\eta_G^2$=effect size estimator generalized Eta-squared, $p_{perm}$=permutation p-value using the method by Kherad-Pajouh and Renaud[5] for non-spherical data with 100000 permutations, minimum value of $p_{perm}$=1/number of permutations). Significant effects with Bonferroni-corrected p < 0.0125 (correcting for 4 ANOVAs) are marked by *. Trends above Bonferroni-corrected threshold are marked by a T.

| effect | *F* | *DFn, DFd* | *p* | *DFn<sub>GG</sub>, DFd<sub>GG</sub>* | $p_{GG}$ | $\eta_G^2$ | $p_{perm}$ |
|---|---|---|---|---|---|---|---|
| **qT1 (n=40)** | | | | | | | |
| age | 0.04 | 1, 38 | 0.834 | 1.00, 38.00 | 0.834 | 0.001 | 0.837 |
| layer | 328.62 | 2, 76 | $3.9 \times 10^{-38}$ * | 1.08, 41.05 | $9.9 \times 10^{-22}$ * | 0.712 | $<10^{-5}$ * |
| age x layer | 4.14 | 2, 76 | 0.020 T | 1.08, 41.05 | 0.046 T | 0.030 | 0.016 T |
| **nQSM (n=34)** | | | | | | | |
| age | 14.49 | 1, 32 | $6.0 \times 10^{-4}$ * | 1.00, 32.00 | $6.0 \times 10^{-4}$ * | 0.188 | $5.8 \times 10^{-4}$ * |
| layer | 18.82 | 2, 64 | $3.7 \times 10^{-7}$ * | 1.15, 36.76 | $5.6 \times 10^{-5}$ * | 0.224 | $<10^{-5}$ * |
| age x layer | 1.94 | 2, 64 | 0.152 | 1.15, 36.76 | 0.171 | 0.029 | 0.151 |
| **pQSM (n=34)** | | | | | | | |
| age | 27.86 | 1, 32 | $8.8 \times 10^{-6}$ * | 1.00, 32.00 | $8.8 \times 10^{-6}$ * | 0.347 | $2.0 \times 10^{-5}$ * |
| layer | 11.58 | 2, 64 | $5.1 \times 10^{-5}$ * | 1.28, 40.86 | $6.7 \times 10^{-4}$ * | 0.123 | $3.0 \times 10^{-5}$ * |
| age x layer | 0.61 | 2, 64 | 0.547 | 1.28, 40.86 | 0.610 | 0.007 | 0.549 |
| **aQSM (n=34)** | | | | | | | |
| age | 30.70 | 1, 32 | $4.1 \times 10^{-6}$ * | 1.00, 32.00 | $4.1 \times 10^{-6}$ * | 0.390 | $2.0 \times 10^{-5}$ * |
| layer | 17.10 | 2, 64 | $1.1 \times 10^{-6}$ * | 1.31, 42.03 | $4.4 \times 10^{-5}$ * | 0.152 | $<10^{-5}$ * |
| age x layer | 1.14 | 2, 64 | 0.325 | 1.31, 42.03 | 0.308 | 0.012 | 0.326 |
