## Supplementary material for "Cortical sensory aging is layer-specific": Figure 2-supplemental table 8

**Figure 2 - supplemental table 8. Post-hoc comparisons to follow up significant main and interaction effects of age on qT1, nQSM, pQSM and aQSM values of the SI hand area using the layer definition of younger adults for both age groups.** Bootstrap Welch paired-sample t-tests were performed on qT1 (myelin) (n=20 younger adults, n=20 older adults), nQSM (calcium) (n=18 younger adults, n=16 older adults), pQSM (iron) (n=18 younger adults, n=16 older adults) and aQSM (mineralization) (n=18 younger adults, n=16 older adults) values to follow up significant main effects of layer (tested differences: outer-middle, middle-inner), Bootstrap Welch two-sample t-tests were performed to follow up significant main effects of layer (inner, middle, outer; tested differences: outer - middle, middle - inner) and significant interaction effects between age (younger, older) and layer (tested differences: younger outer - older outer, younger middle - older middle, younger inner - older inner). Results are given as mean difference values (Mean) and standard errors (SE) in milliseconds (for qT1) or in ppm (for QSM-based estimates), test statistic (t), degrees of freedom (df), p-value (p), bootstrapped p-value ($p_{boot}$), 95% confidence interval (CI), bootstrapped 95% confidence interval ($CI_{boot}$). Number of bootstrap samples=100000, minimum value of $p_{boot}$=1/number of bootstrap samples. Significant effects with Bonferroni-corrected p < 0.0045 (correcting for 11 post-hoc tests) are marked by *. Trends above Bonferroni-corrected threshold are marked by a T.

| comparison | Mean ± SE | *t* | *df* | *p* | $p_{boot}$ | *CI* | $CI_{boot}$ |
|---|---|---|---|---|---|---|---|
| **qT1 (n=40)** | | | | | | | |
| outer > middle | 250.8 ± 17.4 | 14.4 | 39.0 | $3.1 \times 10^{-17}$ * | $<10^{-5}$ * | 215.7, 286.0 | 219.5, 286.2 |
| middle > inner | 256.9 ± 12.7 | 20.3 | 39.0 | $2.6 \times 10^{-22}$ * | $<10^{-5}$ * | 231.3, 282.5 | 235.6, 284.2 |
| younger outer < older outer | -69.8 ± 63.4 | -1.1 | 24.6 | 0.282 | 0.232 | -200.5, 60.9 | -196.7, 45.0 |
| younger middle > older middle | 4.7 ± 33.4 | 0.1 | 25.5 | 0.89 | 0.912 | -64.0, 73.4 | -61.9, 65.6 |
| younger inner > older inner | 42.2 ± 19.0 | 2.2 | 37.1 | 0.033 T | 0.034 T | 3.7, 80.7 | 5.7, 77.9 |
| **nQSM (n=34)** | | | | | | | |
| outer < middle | $-2.7 \times 10^{-3} \pm 2.6 \times 10^{-4}$ | -10.6 | 33 | $3.8 \times 10^{-12}$ * | $<10^{-5}$ * | $-3.2 \times 10^{-3}$, $-2.2 \times 10^{-3}$ | $-3.2 \times 10^{-3}$, $-2.2 \times 10^{-3}$ |
| middle < inner | $-4.1 \times 10^{-4} \pm 6.0 \times 10^{-4}$ | -0.7 | 33 | 0.500 | 0.461 | $-1.6 \times 10^{-3}$, $8.1 \times 10^{-4}$ | $-1.6 \times 10^{-3}$, $6.9 \times 10^{-4}$ |
| **pQSM (n=34)** | | | | | | | |
| outer > middle | $6.4 \times 10^{-4} \pm 6.3 \times 10^{-4}$ | 1.0 | 33 | 0.322 | 0.282 | $-6.5 \times 10^{-4}$, $1.9 \times 10^{-3}$ | $-5.4 \times 10^{-4}$, $1.9 \times 10^{-3}$ |
| middle > inner | $2.0 \times 10^{-3} \pm 3.1 \times 10^{-4}$ | 6.5 | 33 | $2.3 \times 10^{-7}$ * | $<10^{-5}$ * | $1.4 \times 10^{-3}$, $2.7 \times 10^{-3}$ | $1.4 \times 10^{-3}$, $2.6 \times 10^{-3}$ |
| **aQSM (n=34)** | | | | | | | |
| outer > middle | $1.0 \times 10^{-3} \pm 4.3 \times 10^{-4}$ | 2.4 | 33 | 0.023 T | 0.028 T | $1.5 \times 10^{-4}$, $1.9 \times 10^{-3}$ | $1.9 \times 10^{-4}$, $1.9 \times 10^{-3}$ |
| middle > inner | $1.4 \times 10^{-3} \pm 2.7 \times 10^{-4}$ | 5.3 | 33 | $8.7 \times 10^{-6}$ * | $<10^{-5}$ * | $8.6 \times 10^{-4}$, $1.9 \times 10^{-3}$ | $9.0 \times 10^{-4}$, $1.9 \times 10^{-3}$ |
