## Supplementary material for "Cortical sensory aging is layer-specific": Figure 3-supplemental figure 1

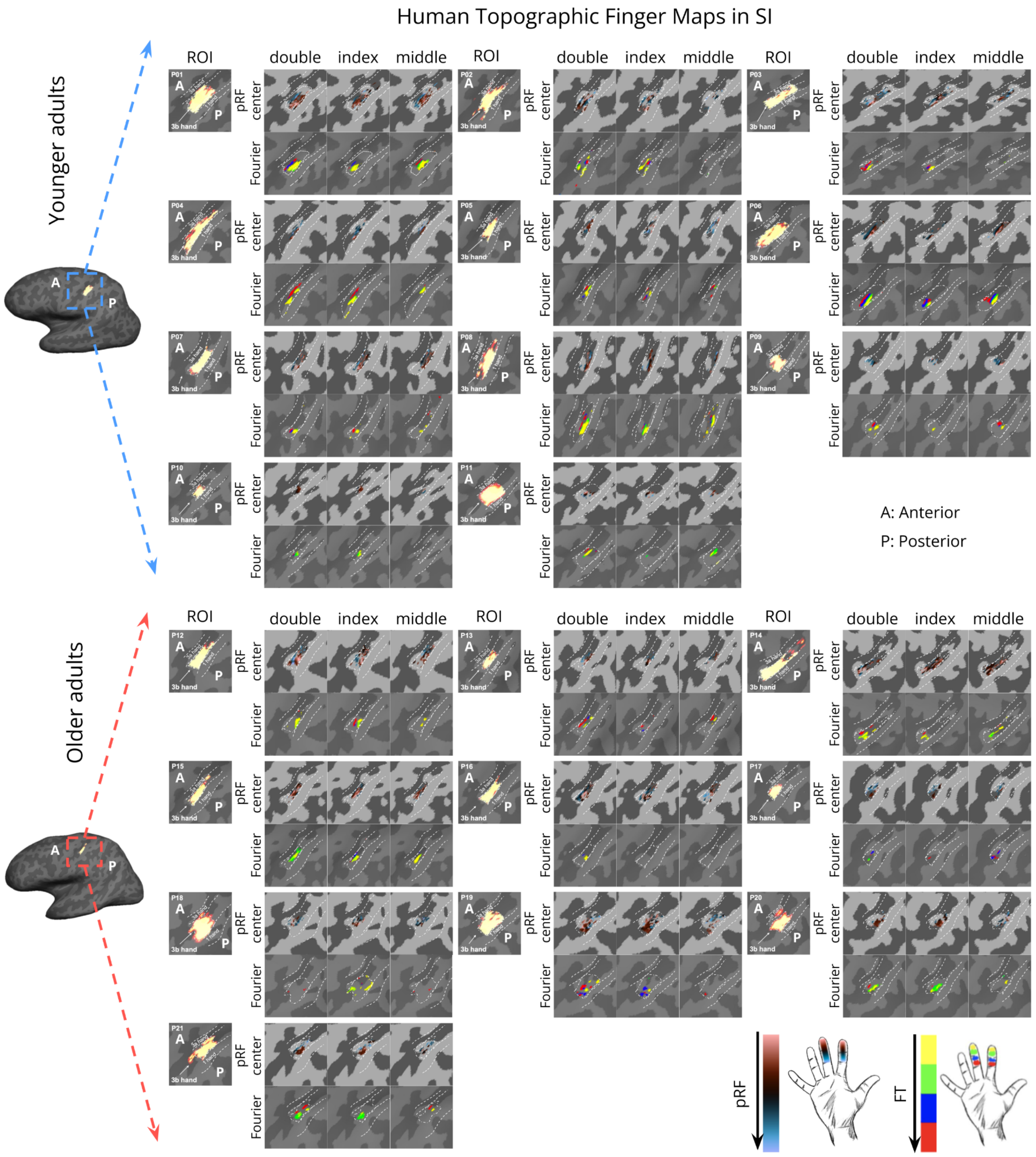


**Figure 3 - supplemental figure 1. Topographic maps of the index finger and the middle finger representation in area 3b of younger adults (n=11) and older adults (n=10)**. Each map is plotted onto the individual surface. Shown are topographic maps extracted using population receptive field (pRF) modeling (first rows in each panel) and Fourier-based mapping (second rows in each panel). ROI = Region-of-interest (hand area of contralateral area 3b).
