## Supplementary material for "Cortical sensory aging is layer-specific": Figure 3-supplemental figure 2

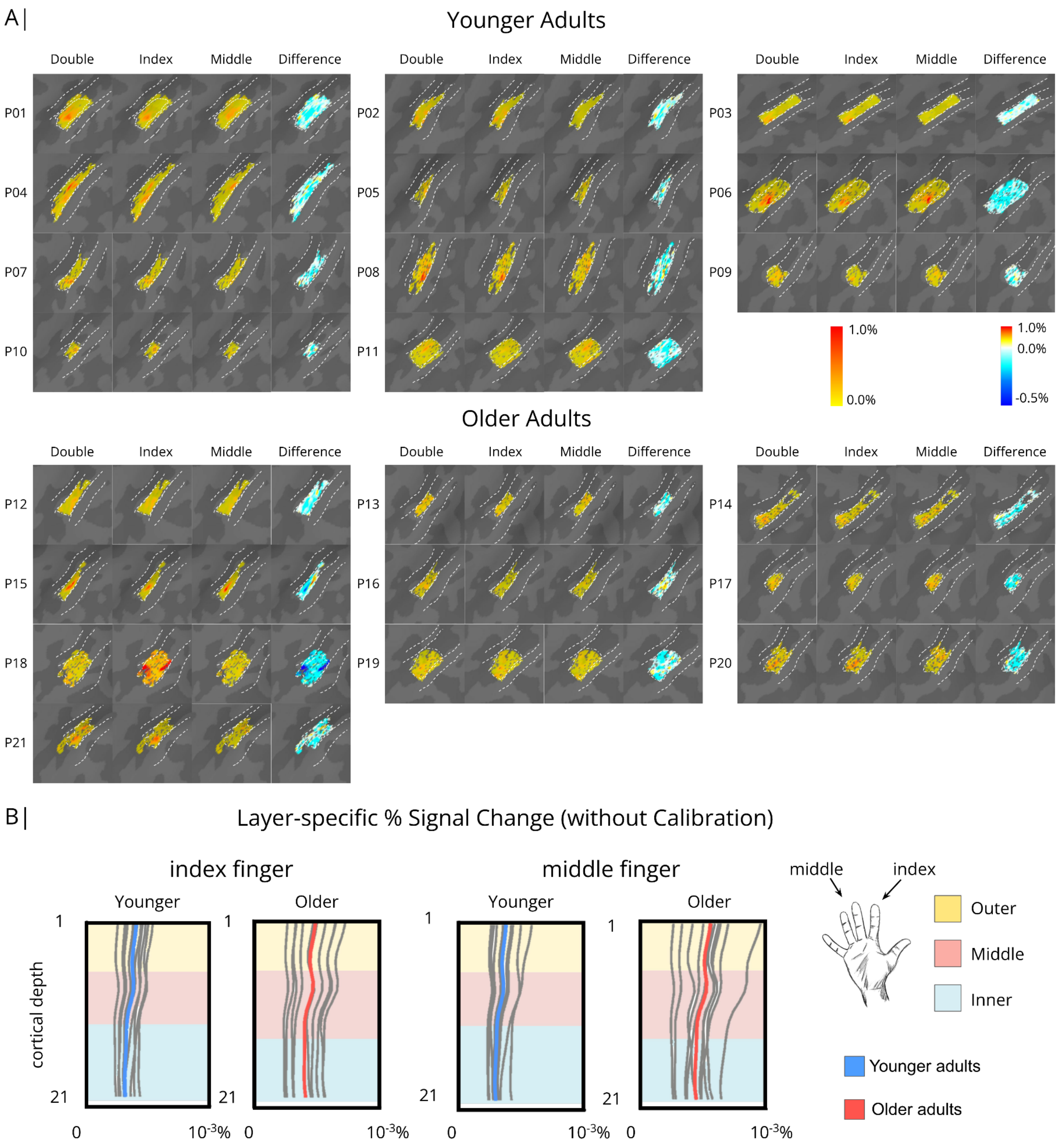


**Figure 3 - supplemental figure 2. Mesoscale functional architecture of finger representations in area 3b. (A)** % signal change map for double finger condition (index and middle finger), index finger condition, middle finger condition, and the inhibition maps (Difference=index finger condition+middle finger condition-double finger condition) during coactivation for younger (n=11) and older (n=10) adults. Each map was plotted onto the individual surface. (**B)** Layer-specific %signal change of index finger and middle finger representations in SI extracted at different cortical depths for younger (blue) and older (red) adults. Note that the calculation was performed on the BOLD signals before calibration.
