## Supplementary material for "Cortical sensory aging is layer-specific": Figure 3-supplemental table 1

**Figure 3 - supplemental table 1. Bayesian independent-sample t-tests were performed on younger (n=11) and older (n=10) adults.** The alternative hypothesis $H_1$ is specified as young < old, and the null hypothesis $H_0$ is specified as no difference between younger and older adults.

| | Central peak | | | | |
|---|---|---|---|---|---|
| | Younger adults n=11 | Older adults n=10 | | | |
| Condition | Mean±SD | Mean±SD | $BF_{+0}$ | error% | 95% Credible Interval |
| index | 0.002±3.548×10$^{-4}$ | 0.002±6.445×10$^{-4}$ | 3.146 | ~ 3.333×10$^{-5}$ | -1.454, -0.054 |
| middle | 0.002±3.795×10$^{-4}$ | 0.002±7.747×10$^{-4}$ | 15.533 | ~ 6.461×10$^{-4}$ | -1.695, -0.089 |

| | Signal decay | | | | |
|---|---|---|---|---|---|
| | Younger adults n=11 | Older adults n=10 | | | |
| Condition | Mean±SD | Mean±SD | $BF_{+0}$ | error% | 95% Credible Interval |
| index | 0.002±4.824×10$^{-4}$ | 0.003±8.493×10$^{-4}$ | 2.769 | ~ 2.756×10$^{-5}$ | -1.380, -0.047 |
| middle | 0.002±5.162×10$^{-4}$ | 0.003±0.001 | 13.405 | ~ 5.665×10$^{-4}$ | -1.675, -0.096 |
