## Supplementary material for "Cortical sensory aging is layer-specific": Figure 3-supplemental table 2

**Figure 3 - supplemental table 2. Bayesian independent-sample t tests were performed on younger (n=11) and older (n=10) adults.** The alternative hypothesis $H_1$ is specified as young < old, and the null hypothesis $H_0$ is specified as no difference between younger and older adults.

| | | **Younger adults** n=11 | **Older adults** n=10 | | | |
|---|---|---|---|---|---|---|
| | | Mean ± SD | Mean ± SD | $BF_{+0}$ | error% | 95% Credible Interval |
| % signal change (%) | index | 0.118 ± 0.024 | 0.149 ± 0.091 | 0.982 | ~ $1.854\times10^{-6}$ | -1.173, -0.026 |
| | middle | 0.111 ± 0.036 | 0.127 ± 0.033 | 0.924 | ~ $9.772\times10^{-7}$ | -1.151, -0.025 |
| pRF size (σ) | index | 0.427 ± 0.107 | 0.633 ± 0.205 | 11.47 | ~ $4.490\times10^{-4}$ | -1.998, -0.189 |
| | middle | 0.448 ± 0.150 | 0.514 ± 0.244 | 0.70 | ~ $8.650\times10^{-7}$ | -1.048, -0.019 |
