## Supplementary material for "Cortical sensory aging is layer-specific": Figure 3-supplemental table 3

**Figure 3 - supplemental table 3. Bayesian independent sample t-tests were performed on younger (n=11) and older (n=10) adults.** For pRF size (σ) difference between double finger condition and the average of single finger conditions, the alternative hypothesis is $H_1$ is specified as $\sigma_{older} < \sigma_{younger}$. For % signal change difference between double finger condition and the sum of single finger conditions, the alternative hypothesis $H_1$ is specified as $\%_{older} < \%_{younger}$. The null hypothesis indicates no difference between younger and older adults on the pRF size difference and % signal change difference.

| | **Younger adults** n=11 | **Older adults** n=10 | | | |
|---|---|---|---|---|---|
| | Mean ± SD | Mean ± SD | $BF_{+0}$ | error% | 95% Credible Interval |
| pRF size (σ) difference | 0.152 ± 0.144 | 0.196 ± 0.099 | 0.734 | ~ $6.290 \times 10^{-7}$ | -0.663, -0.007 |
| % signal change (%) difference | 0.081 ± 0.022 | 0.124 ± 0.096 | 1.424 | ~ $8.959 \times 10^{-5}$ | -1.330, -0.040 |
