## Supplementary material for "Cortical sensory aging is layer-specific": Figure 4-supplemental figure 1

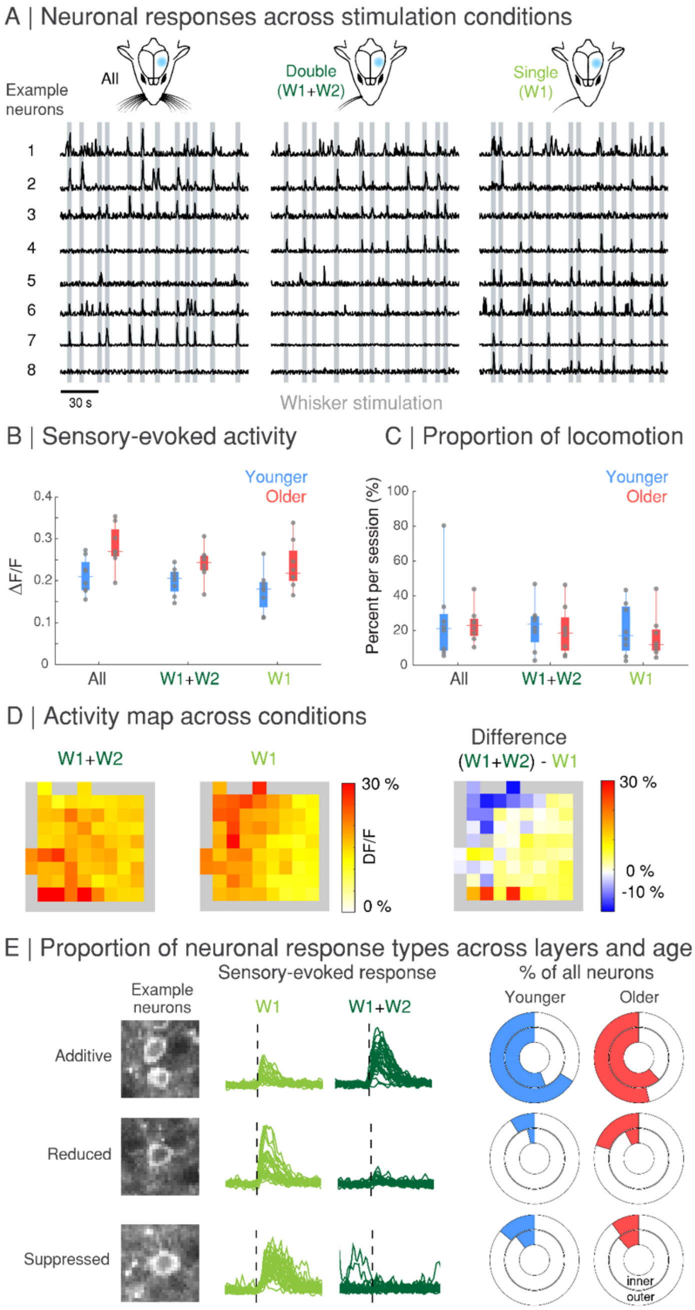


**Figure 4 - supplemental figure 1. Age-related functional changes in mouse barrel cortex following single whisker and double whisker coactivation. (A)** Calcium activity from example neurons (1-8) for the three stimulation conditions, all whiskers (All), two neighboring whiskers (W1+W2; double stimulation condition), and single whisker (W1; single stimulation condition). **(B)** Average ΔF/F responses across all neurons within a field-of-view for younger adult mice (1519 neurons from n=8 mice) and older adult mice (1958 neurons from n=8 mice) during sensory-evoked airpuff stimulation of all whiskers as well as double stimulation and single stimulation conditions. Significant effect of age ($F_{(2,42)}$=18.18, p < 0.001), with older adult mice showing larger sensory-evoked excitatory neuronal responses (effect of stimulation condition, $F_{(2,42)}$=3.22, p=0.050; no significant interaction, $F_{(2,42)}$=0.38, p=0.689; two-way mixed-effects ANOVA, constrained (type III) sum of squares; Tukey-Kramer correction). **(C)** Percentage of time that mice spent running during experimental sessions did not differ significantly across age (p=0.525) or stimulation condition (p=0.556; with no interaction (p=0.978) two-way mixed-effects ANOVA, constrained (type III) sum of squares), indicating that locomotion alone is unlikely to contribute to the differences in sensory-evoked activity observed across age (see **Fig. 4A**). **(D)** Sensory-evoked ΔF/F for an example field-of-view, averaged across 50 × 50 pixel spatial bins during double whisker (left) and single whisker (middle) stimulation and the difference of these two conditions (right). **(E)** Example neurons and sensory-evoked ΔF/F that show additive, reduced, or suppressed responses to single (W1) or double (W1+W2) whisker stimulation; percentage of all neurons (younger adult mice [total 1519 neurons from n=8 mice; 1081 outer layer neurons, 438 inner layer neurons] and older adult mice [total 1958 neurons from n=8 mice; 1446 outer layer neurons, 512 inner layer neurons]) showing these response types across outer (layer II/III, outer ring) and inner (layer V, inner ring) cortical layers.
