## Supplementary material for "Cortical sensory aging is layer-specific": Figure 4-supplemental figure 2

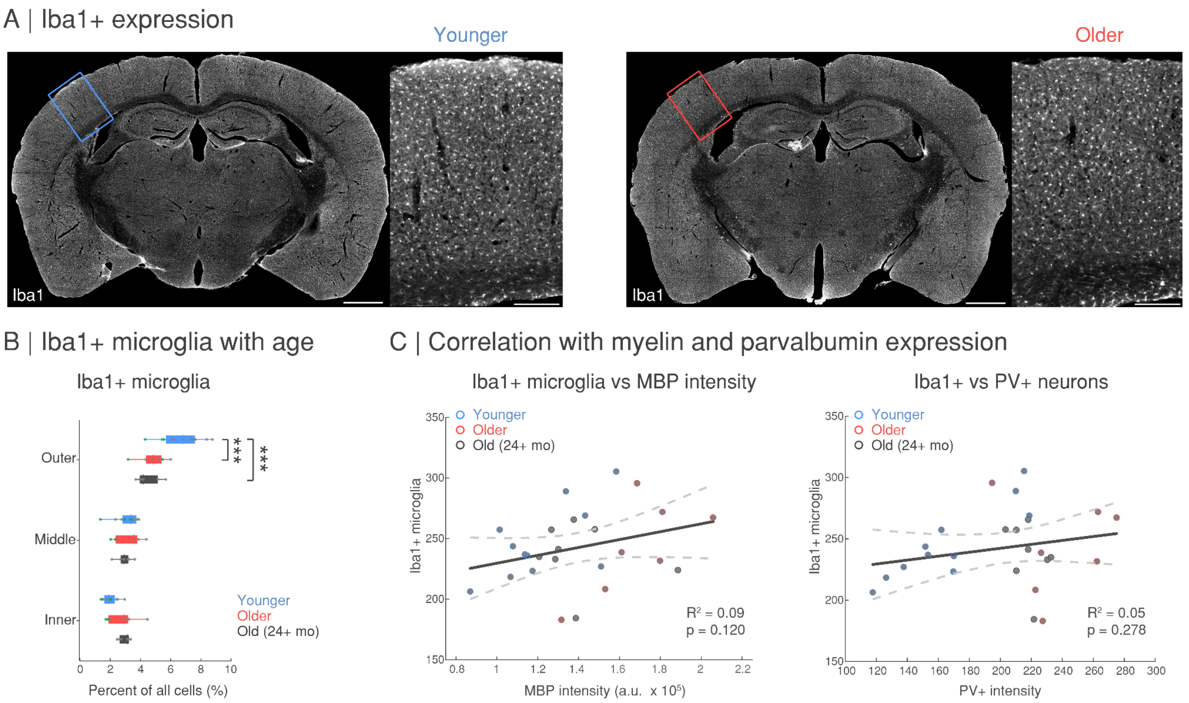


**Figure 4 - supplemental figure 2. Age-related changes in Iba1 immunohistochemistry as a readout of microglial density in mouse barrel cortex. (A)** Representative Iba1 expression in a younger adult and older adult mouse. **(B)** The percentage of Iba1+ microglia (normalized to the DAPI total cell count) across age and outer (layer II/III), middle (layer IV) and inner (layer V/VI) cortical layers from the somatosensory barrel cortex. Significant effect of layer ($F_{(2,69)}$=78.25, p < 0.001) but not age ($F_{(2,69)}$=2.85, p=0.064), and a significant interaction ($F_{(4,69)}$=10.80, p < 0.001, two-way mixed-effects ANOVA, asterisk indicates significance level of *p < 0.05, **p < 0.01, ***p < 0.001, values from Tukey-Kramer corrections). Data from mice with cranial window implantations (5 total, 3 younger and 2 older adult mice) are indicated as green dots. **(C)** Linear regression $R^2$ and 95% confidence intervals (dashed lines) for the number of Iba1+ microglia (average count per sample, summed across cortical depth) and intensity of myelin basic protein (MBP; average intensity per sample, summed across cortical depth) expression (left; $R^2$=0.09, p=0.120) or the number of parvalbumin (PV+) expressing cells (average count per sample, summed across cortical depth; right; $R^2$=0.05, p=0.278) per animal (n=26 mice: younger adult mice [n=11, 2-6 months], older adult mice [n=7, 12-20 months], and mice in old age [n=8, +24 months]). Scale bars in **(A)**, 1 mm on coronal section and 250 μm inset.
