## Supplementary material for "Cortical sensory aging is layer-specific": Figure 5-supplemental table 1

**Figure 5 - supplemental table 1. Comparison of functional map features and behavioral markers between age groups.** Network centrality (Eigenvector centrality), dprime of index finger (D2) 2-point discrimination, and finger discrimination averaged across all fingers are given in arbitrary units, the inverse tactile detection threshold is given in 1/[log10 0.1 mg], the inverse 2-point discrimination threshold is given in 1/mm, sensorimotor integration involving thumb and index finger (D1-D2) is given in seconds. To ensure that higher values indicate better performance, tactile detection and 2-point discrimination thresholds were reversed. We report group means (Mean) and standard deviations (SD). Independent-samples random permutation Welch t-tests were calculated to investigate group differences (t=test statistic, df=degrees of freedom, $p_{perm}$=Monte-Carlo permutation p-value, $CI_{perm}$=95% Monte-Carlo permutation confidence interval, number of permutations=100000, minimum value of $p_{perm}$=1/number of permutations). Significant differences with Bonferroni-corrected threshold of $p < 0.006$ are marked by *, trends above Bonferroni-corrected threshold are marked by a T.

| | **Younger Adults** n = 20 | **Older Adults** n = 20 | **Group Differences** | | | |
|---|---|---|---|---|---|---|
| | Mean ± SD | Mean ± SD | $t$ | *df* | $p_{perm}$ | $CI_{perm}$ |
| % signal change D2 | 0.22 ± 0.09 | 0.15 ± 0.08 | 2.41 | 36.41 | 0.021 T | 0.009, 0.121 |
| network centrality D2 | 1.03 ± 0.07 | 1.00 ± 0.06 | 1.06 | 24.29 | 0.296 | -0.021, 0.073 |
| network centrality Hand | 1.04 ± 0.05 | 1.01 ± 0.04 | 1.46 | 23.62 | 0.158 | -0.009, 0.064 |
| inverse tactile detection D2 | 0.32 ± 0.06 | 0.27 ± 0.04 | 2.96 | 35.70 | 0.005 * | 0.013, 0.079 |
| inverse 2-point discrimination threshold D2 | 0.56 ± 0.10 | 0.34 ± 0.09 | 7.12 | 34.99 | $<10^{-5}$ * | 0.130, 0.321 |
| dprime 2-point discrimination D2 | 1.20 ± 0.32 | 1.33 ± 0.31 | -1.21 | 35.00 | 0.233 | -0.325, 0.077 |
| finger discrimination hand | 1.71 ± 0.54 | 1.12 ± 0.62 | 2.84 | 27.91 | 0.008 T | 0.145, 1.027 |
| sensorimotor integration D1-D2 | 8.61 ± 2.18 | 6.36 ± 1.31 | 3.95 | 31.10 | $4.8 \times 10^{-4}$ * | 0.952, 3.561 |
