## Supplementary material for "Cortical sensory aging is layer-specific": Figure 5-supplemental table 2

**Figure 5 - supplemental table 2. Nonparametric Wilcoxon Signed-Rank tests comparing measures of hand dexterity between younger (n=21) and older (n=17) adults** using different finger(s) on (1) time in seconds used to complete the Grooved Pegboard Test, (2) the number of holes (n) filled within the given time interval in the O'Connor Dexterity Test, (3) number of pairs completed (n) within the given time interval in the Small Motor Test, and (4) accuracies on distinguishing texture in the Texture Roughness Test. D1=thumb, D2=index finger and D3=middle finger. Significant effects with p < 0.01 are marked by *, with p < 0.001 are marked by **, and with p < 0.0001 are marked by ***.

| | | **Younger Adults** n = 11 | **Older Adults** n = 10 | **Group Differences** | | |
|---|---|---|---|---|---|---|
| | | Mean ± SD | Mean ± SD | W | *p* | $r_{rb}$ |
| Grooved Pegboard Test (s) | D1 + D2 | 102.48 ± 19.77 | 142.18 ± 36.23 | 304.50 | $1.14\times10^{-4}$*** | 0.71 |
| | D1 + D2 + D3 | 67.62 ± 9.97 | 81.12 ± 12.01 | 289.50 | $5.82\times10^{-4}$*** | 0.62 |
| O'Conner Hand Dexterity Test (n) | D1 + D2 | 26.29 ± 7.76 | 21.65 ± 8.33 | 101.00 | 0.012** | -0.434 |
| | D1 + D2 + D3 | 19.24 ± 3.34 | 17.43 ± 3.39 | 91.50 | 0.006** | -0.487 |
| Small Motor Test (n) | D1 + D2 | 19.24 ± 3.38 | 17.43 ± 3.39 | 233.50 | 0.054 | 0.308 |
| | D1 + D2 + D3 | 21.18 ± 4.19 | 18.91 ± 2.70 | 242.00 | 0.032* | 0.356 |
| Texture Roughness Test (%) | D2 | 88.2 ± 7.1 | 87.7 ± 6.9 | 169.00 | 0.616 | -0.053 |
| | D2 + D3 | 87.7 ± 7.4 | 88.7 ± 5.7 | 195.00 | 0.317 | 0.092 |
